# miRNA/mRNA analysis of increased TGF-β pathways drive epithelial-mesenchymal transition and regulatory T cell differentiation

**DOI:** 10.1101/2025.06.23.661210

**Authors:** Toni Darville, Xuejun Sun, Yu Zhang, Catherine M. O’Connell, Neha Mokashi, Weiming Tang, Aakash Bhardwaj, Bryce Duncan, Charles W. Andrews, Harold Weisenfeld, Xiaojing Zheng

**Author notes:** shared first authorship (Toni Darville, Xuejun Sun). Correspondence: Xiaojing Zheng, PhD, University of North Carolina at Chapel Hill, 4018A Mary Ellen Jones Building, CB# 7509, 110 Manning Drive, Chapel Hill, NC 27599-7509, USA.

## Abstract

*Chlamydia trachomatis* genital tract infection is linked to severe reproductive complications in women, including ectopic pregnancy, infertility, and adverse pregnancy outcomes. Mouse models of infection suggest that chlamydia*-*induced dysregulation of microRNAs (miRNAs) can drive harmful cytokine responses, pathogenic epithelial-mesenchymal transition (EMT), and fibrosis. To investigate these mechanisms in humans, we profiled miRNA and mRNA expression in endometrial biopsies from women with endometrial infection (Endo+) and compared them to profiles from women with cervix-only infection (Endo-) or no infection. Ingenuity Pathway Analysis (IPA) revealed that Endo+ tissues had upregulated genes associated with innate and adaptive immune response pathways, as well as EMT regulation, while downregulated genes were linked to cell cycle control. An integrative miRNA-mRNA analysis, which combined a review of published miRNA regulation in human infections and immune responses with IPA’s miRNA target filter, identified differentially expressed miRNAs that modulate these pathways in the endometrium of Endo+ women. Functional annotation of these miRNAs showed a predominance of downregulated miRNAs that typically suppress EMT and regulatory T cell (Treg) differentiation, along with miRNAs that usually enhance Th17 responses. Comparisons with previously identified mRNA pathways in blood samples from women with endometrial *Chlamydia* infection indicated that alterations in TGF-β signaling and EMT were specific to the endometrium. Overall, the miRNA-mRNA interactions inferred from Endo+ tissue suggest increased activity in TGF-β pathways that promote enhanced EMT and Treg differentiation, while reducing Th17 activation. These changes highlight a dual potential for promoting tissue scarring while dampening inflammatory responses that could otherwise limit infection.

## Introduction

Sexually transmitted *Chlamydia trachomatis* (Ct) genital infections represent a global public health challenge due to their high prevalence and severe reproductive health consequences. These include pelvic inflammatory disease (PID), chronic pelvic pain, infertility, and ectopic pregnancy (1, 2). Infertility and ectopic pregnancy after Ct infection is primarily caused by fibrotic scarring of the oviducts. Additionally, studies have linked Ct with adverse pregnancy outcomes such as stillbirth, infant death, spontaneous abortion, preterm labor, small-for-gestational-age infants, and postpartum endometritis (3). This suggests that Ct may cause lasting endometrial damage.

In the female reproductive tract, immune cells in the vagina and cervix provide the first line of defense against infection, whereas immune cells in the endometrium play a dual role: eliminating pathogens that breach the cervical barrier and maintaining immune tolerance of the embryo during pregnancy. Regulatory T (Treg) cells in the uterus are critical for immune tolerance to foreign fetal antigens(4, 5). Their secretion of anti-inflammatory cytokines such as IL-10 and TGF-β, and expression of inhibitory molecules like CTLA-4 and PD-1, suppress maternal immune responses that could otherwise target the fetus. In contrast, Th17 cells drive proinflammatory responses that can be harmful. The balance between these cell types is crucial for a healthy pregnancy. During Ct infection, the induction of Tregs may hinder the development of effective T cell immunity (6), whereas the activation of chlamydial-specific Th17 cells appears to enhance resistance to reinfection (7). Therefore, while a higher Treg/Th17 ratio supports reproductive health, it may compromise host defense against Ct.

Animal models have also shown that pro-inflammatory signaling and immune cell recruitment are initiated upon Ct infection of host epithelial cells (8, 9). Engagement of pathogen recognition receptors (PRRs) elicits a rapid influx of neutrophils that fail to kill the bacteria while releasing tissue-damaging molecules such as reactive oxygen species and matrix metalloproteases (10–15). Eventually, chlamydia-specific IFN-γ-producing CD4 T cells, activated by dendritic cells, resolve infection. However, danger-associated molecular patterns (DAMPs) released by chlamydiae or dying epithelial cells can perpetuate inflammation through induction of production of TNFα and IL-1α by adjacent epithelium or influxing innate cells, creating a feed-forward loop of tissue damage (11, 16, 17). Furthermore, epithelial-mesenchymal transition (EMT)—a dysregulated tissue repair process controlled by microRNAs (miRNAs)—has been implicated in oviduct fibrosis in chlamydia-infected mice (18). During EMT, epithelial cells lose cell-cell contacts and acquire mesenchymal characteristics, including elevated production of extracellular matrix proteins, before transitioning into myofibroblasts.

Mouse models of chlamydial infection demonstrated that the induction of EMT involves TNFα signaling, caspase activation, and cleavage inactivation of Dicer (18, 19) an RNase III enzyme that processes RNA into microRNAs. MicroRNAs (miRNAs) are small non-coding RNAs that play a crucial role in post-transcriptional gene regulation, primarily through translational repression and mRNA degradation. However, miRNAs can also enhance gene expression and translation under certain conditions (20, 21). miRNAs are predicted to control the activity of approximately 30% to 50% of all protein-coding genes. A single miRNA can target multiple mRNAs, while multiple miRNAs can collaborate to finely tune the expression of a single mRNA target (22). Chlamydia infection in mice was associated with reduced expression of miRNAs that normally suppress EMT, fibrosis and tumorigenesis, alongside increased expression of proteins associated with EMT and fibrosis (18, 19)

Human salpingeal tissues are challenging to obtain but minimally invasive endometrial suction catheter biopsies can be collected in an office setting (23). Histological evidence of endometritis correlates with salpingitis observed via laparoscopy (24) and is associated with an increased risk of infertility in women with symptomatic or asymptomatic Ct infection (25). Histologic features resembling endometritis have also been described in surgically removed Fallopian tubes (26), supporting the use of endometrial biopsies for investigating pathogenic mechanisms underlying Ct-induced disease.

In this study, we conducted an integrated analysis of endometrial mRNA and miRNA profiles in cisgender women with high exposure to Ct. Our findings identified an infection-driven mRNA signature that highlights active innate and adaptive immune signaling pathways, along with epithelial-mesenchymal transition (EMT), in women with endometrial Ct infection compared to those with cervical Ct infection only and uninfected women. A parallel miRNA analysis revealed significant downregulation of several miRNAs that typically suppress mRNAs involved in activating TGF-β-related pathways, which drive EMT and regulatory T cell (Treg) differentiation. Additionally, a partially overlapping subset of miRNAs that typically enhance proinflammatory Th17 differentiation was also downregulated. The suppression of these miRNAs during endometrial Ct infection appears to release their regulatory effects, facilitating EMT and increasing the potential for tissue scarring. However, the miRNA-mediated shift in the Treg/Th17 balance, favoring Tregs, may simultaneously act to dampen inflammation.

## Materials and Methods

### Ethics statement

This study adhered to the Declaration of Helsinki guidelines, and all participants provided written informed consent prior to participation. The study protocols were approved by the Institutional Review Boards for Human Subjects Research at the University of North Carolina and the University of Pittsburgh.

### Study population

Endometrial biopsy samples were obtained from cisgender female participants enrolled in two independent cohorts with high exposure to Ct infection. Anaerobes and Clearance of Endometritis (ACE) Cohort: Participants were women clinically diagnosed with pelvic inflammatory disease (PID) (27, 28). Diagnostic criteria for enrollment included cervical motion tenderness, uterine tenderness, or adnexal tenderness observed during pelvic examination in sexually active young women experiencing pelvic or lower abdominal pain. T Cell Response Against Chlamydia (TRAC) Cohort: Participants were asymptomatic women identified as being at high risk for Ct infection (29). At enrollment, demographic, behavioral, and medical history data were collected. General physical and pelvic exams were conducted, and blood samples were obtained for immune studies. Participants underwent endometrial biopsy sampling using suction catheters. Additional assessments included Gram-stained vaginal smears for bacterial vaginosis using Nugent scores (30), as well as testing for Ct, *Neisseria gonorrhoeae* (Ng), and *Mycoplasma genitalium* (Mg) in cervical swabs and endometrial biopsies using nucleic acid amplification tests. Histopathological analyses of biopsies were performed (25, 31).

Participants were categorized into three primary groups based on infection status: (1) Endo+: Ct detected in both the endometrium and cervix. Endo+ women were further categorized into subgroups of women with clinical PID (Endo+, PID+), and asymptomatic women (Endo+, PID-). (2) Endo-: Ct detected in the cervix only, without endometrial infection. (3) Uninfected: No Ct infection. For miRNA analyses, women with PID-like symptoms with negative tests for Ct, Ng, and Mg were included as an additional comparison group.

### Study design and workflow

The study workflow is depicted in **Figure 1**. Endometrial biopsy samples were analyzed for mRNA and miRNA expression. Principal Component Analysis (PCA) was used to explore transcriptomic relationships among cohort participants. Differentially expressed (DE) mRNAs and miRNAs were identified between Endo+ and Endo-/Uninfected groups. Functional annotations of DE mRNAs and miRNA-mRNA pairs were conducted using Ingenuity Pathway Analysis (IPA). Endometrial mRNA data were also compared with previously published blood mRNA data from the same cohorts (32).

**Figure 1.**
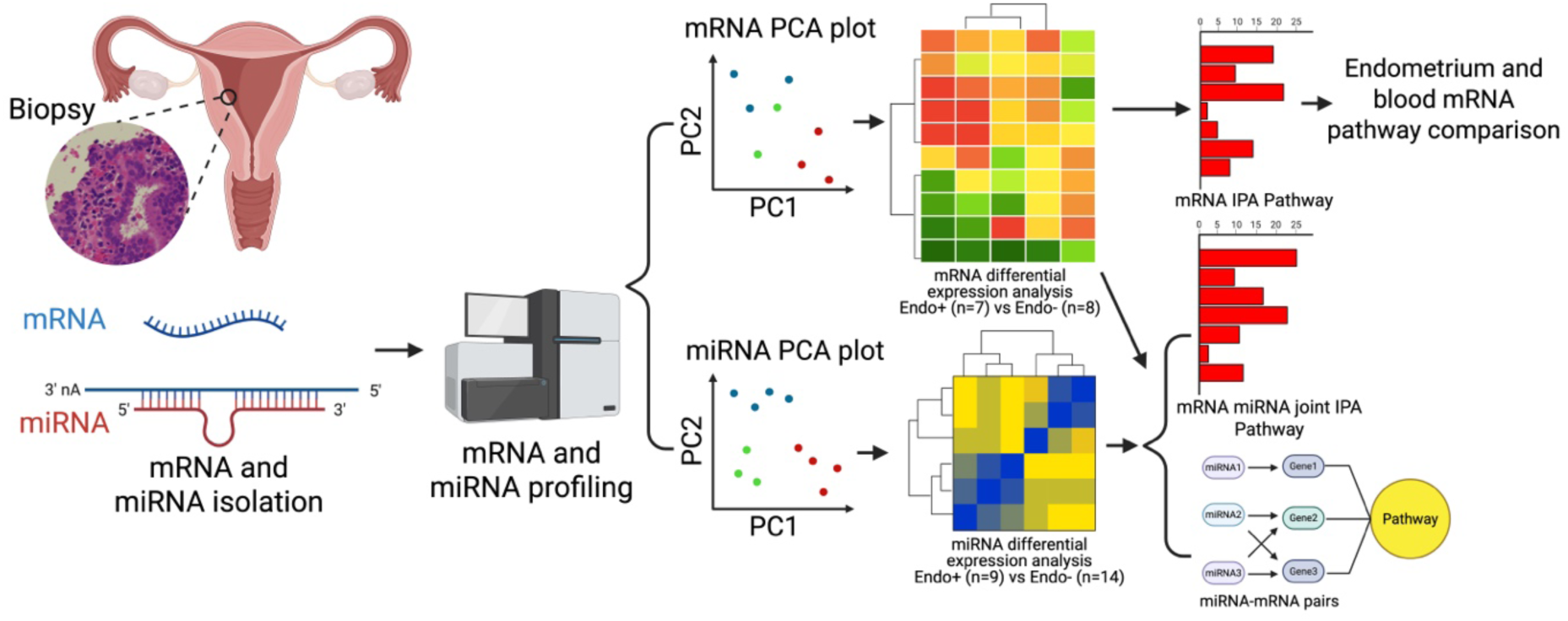
Workflow depicting sample processing, mRNA and miRNA profiling and subsequent analytical methods. Endometrial biopsy samples were processed to profile mRNA from 7 Endo (+) and 8 Endo (-) women, including 4 cervix (+) only and 4 uninfected, and miRNA from 9 Endo (+) and 14 Endo (-) women, including 5 cervix(+) only and 9 uninfected. Relationships among cohort participants based on their transcriptomes were revealed by Principal Component Analysis (PCA). Differentially expressed (DE) mRNAs and miRNAs between Endo+ and Endo-biopsies were determined and functional annotation of DE mRNA and miRNA-mRNA pairs were provided by IPA. Additionally, endometrial biopsy generated mRNA data were compared to previously published blood mRNA data generated from participants in the T Cell Response Against Chlamydia (TRAC) cohort (32).

### mRNA and miRNA data collection and processing

mRNA Extraction and Profiling: Total RNA was extracted from endometrial biopsies stored at -80°C in tubes containing RNA/DNA Shield (Zymo Research, Irvine, CA). After thawing, samples were weighed, minced, and processed for simultaneous DNA and RNA extraction using the Quick DNA/RNA™ isolation kit according to the manufacturers protocol (Zymo) with on column DNAse I treatment prior to total RNA elution. Libraries were prepared using the Ovation SoLo RNA-Seq kit (NuGen Technologies) and sequenced on an Illumina HiSeq2500 platform in the High Throughput Sequencing Facility (HTSF) at the University of North Carolina. Gene expression quantification used BBMap (v37.25) and samtools (v1.4.1). Blood mRNA pathway data had been previously generated by microarray hybridization analysis using samples obtained from ACE and TRAC participants (31, 32). miRNA Extraction and Profiling: miRNAs were isolated using HTG EdgeSeq reagents according to the user manuscript, and libraries were synthesized at the HTSF at the University of North Carolina. Barcoded samples were pooled, and sequencing was performed on an Illumina NextSeq 500 platform using a 75-cycle High Output v2 kit. Count data were generated by the EdgeSeq parser software (HTG Molecular Diagnostics, Inc.).

### Data accession

All endometrial mRNA and miRNA profiles have been deposited in the Gene Expression Omnibus database (https://www.ncbi.nlm.nih.gov/geo/) GEO accession number: GSE290615 for endometrial mRNA, and GSE289941 for endometrial miRNAs. The blood mRNA profiles were retrieved from the previous study (32) with GEO accession number GSE110106.

### Statistical analysis

Demographic Data: One-way ANOVA was used for numerical variables, while Fisher’s exact test or Chi-square test was applied for categorical variables. RNA-Seq Data: PCA with ComBat-seq (33) correction was applied to address batch effects. mRNAs with expression levels <20 in >50% of samples were excluded, leaving 16,183 mRNAs for analysis. miRNAs with expression levels below mean +3 standard deviations of negative controls were filtered, retaining 253 miRNAs. Normalization was performed using DESeq2 (v1.36.0) (34). PCA of variance by R was used to identify the inherent pattern of samples. Differential Expression Analysis: DE mRNAs and miRNAs were identified using DESeq2 with false discovery rate (FDR) <0.05. All analyses, unless specified, were completed in R (version 4.1.0). Within-sample group clustering heatmaps were generated to visualize DE miRNA expression across groups.

### Functional annotation and pathway enrichment analysis of mRNAs

Ingenuity Pathway Analysis (IPA) (QIAGEN Inc., https://digitalinsights.qiagen.com/IPA) identified enriched pathways for DE mRNAs, using Fisher’s exact test. Regulatory T cell-related mRNAs not included in IPA were sourced from Gene Ontology (GO) (35) and manually curated from literature.

### Integrated miRNA-mRNA analysis

Putative target mRNAs of DE miRNAs were identified through IPA’s “miRNA target filter” which incorporates both published experimental and high-confidence computational data. mRNA targets were further filtered to include only those that were also identified as DE mRNAs (FDR<0.05) in this study. We performed a functional annotation of the negatively correlated miRNA-mRNA pairs to identify significantly enriched pathways.

### Systematic review of miRNA functions

A systematic review identified studies examining DE miRNA functions in human immune-related processes. PubMed searches from 1990 to 2024 used the following search query: all miRNAs AND (infect OR immune OR autoimmune) AND (human OR patient OR healthy donor) AND ("T cell" OR "B cell" OR lymphocyte OR CD4 OR CD8 OR Treg OR Th1 OR Th2 OR Th17 OR Tfh OR neutrophil OR monocyte OR macrophage OR “NK cell” OR “epithelial cell” OR "dendritic cell") AND (expression OR function OR Differentiation OR activated OR activation OR inhibit OR suppress OR stimulation) NOT ("beta-cell" OR methylation OR stem OR endothelial OR pregnancy). Tumor-related and non-immune studies were excluded, leaving 361 full-text studies curated for miRNA effects on immune and epithelial cell responses including directionality of their effects.

### Comparison of DE mRNA enriched pathways detected in the endometrium versus blood

We previously identified blood DE mRNAs from women with Ct-induced pelvic inflammatory disease (PID) (Endo+) to those from asymptomatic women with cervical infection only and uninfected women (Endo-/Uninfected) (32). Using Bonferroni-adjusted P values <0.05 as the significance threshold, we compared the significantly enriched pathways for DE mRNAs from this endometrial study and the aforementioned blood study.

## Results

### Characteristics of study participants

The characteristics of study participants included in the mRNA (N = 15) and miRNA (N = 23) analyses are summarized in **Supplementary Tables 1 and 2**. Participant group sizes were determined by the quality of mRNA and miRNA isolated from tissue specimens. Most participants were young, African American, unmarried, had some college education, and were insured by Medicaid. A majority reported previous Ct infection, had anti-Ct antibodies, and did not have bacterial vaginosis (BV) by Nugent’s criteria (30) or coinfection with Ng and/or Mg. Demographic variables such as age, race, education, insurance status, and contraceptive use did not differ significantly among groups.

### Distinct gene expression profiles in Endo+ women contrast with minimal impact of cervical Ct infection

Unsupervised principal component analysis (PCA) was performed to assess whether global gene expression profiles were associated with infection status or disease extent. PCA of mRNA data (**Figure 2A**) showed that most Endo+ women formed a distinct cluster, separate from the Endo- and Uninfected groups, with one outlier in the Endo+ group. Similarly, PCA of miRNA data (**Figure 2B**) indicated that Endo+ women clustered apart from the Endo- and Uninfected groups. However, PCA did not detect differences between endometrial mRNA or miRNA expression profiles of women with cervical Ct infection or those who were uninfected. The lack of separation between Endo- and Uninfected groups suggests that cervical Ct infection does not significantly impact endometrial gene expression.

**Figure 2.**
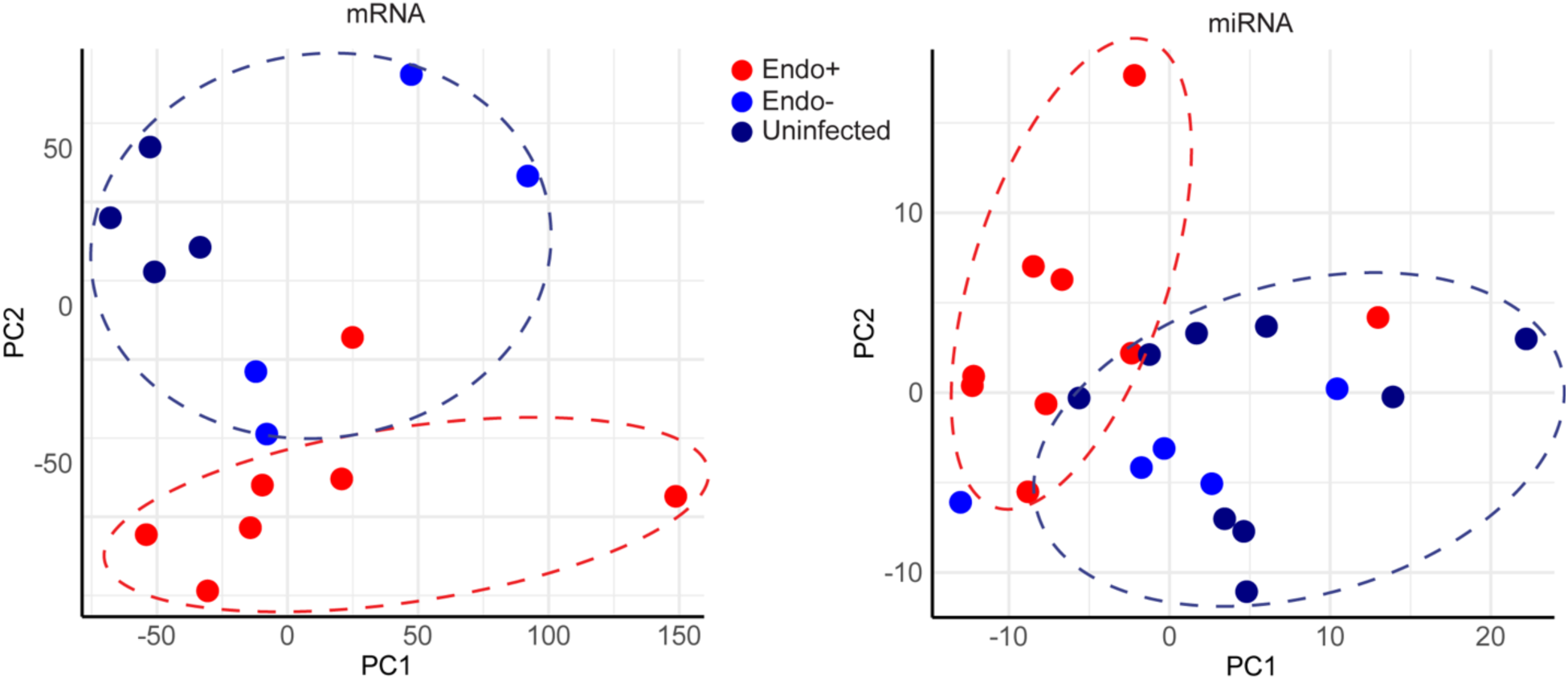
Principal component analysis (PCA) of endometrial transcriptome (A) mRNA and (B) miRNA. Each dot represents one subject, with infection status indicated by color. Red denotes endometrial infection (Endo+), light blue denotes absence of endometrial infection but positive cervical infection (Endo-), and dark blue (Uninfected). The x-axis represents the first principal component, PC1, which accounts for the largest variance of mRNA (A) or miRNA (B) expression, and the y axis, PC2, explains the second largest variance.

### Endometrial inflammation was detected in Endo+ and non-STI PID cases

Biopsied tissues from Endo+ women, both symptomatic and asymptomatic, and from women with non-STI-induced PID showed plasma cell and lymphocyte infiltrates in the endometrial stroma. Neutrophils were also observed within the basal lamina of the epithelium and in gland lumens in some Endo+ biopsies. In contrast, 6 evaluable biopsis from women with cervical Ct infection only and 9 evaulable biopsies from uninfected women either lacked inflammatory cells or contained rare mononuclear cells. The remaining biopsies had insufficient tissue for histological evaluation (**Figure 3A-D**). These pathological data align with the data from PCA analysis indicating clear separation of endometrial transcriptional profiles of Endo+ women from those with Ct infection limited to the cervix.

**Figure 3.**
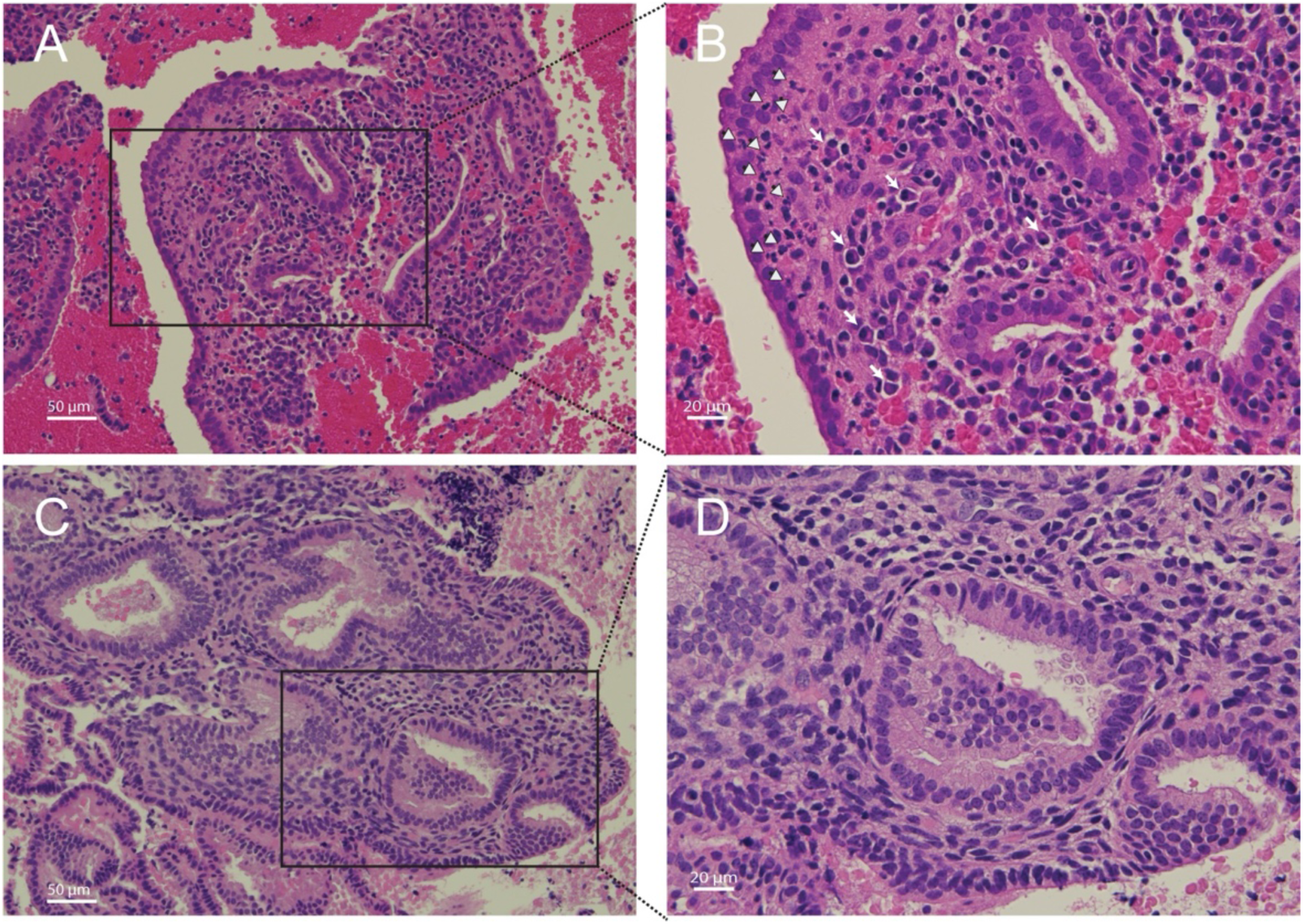
Histopathology of *Chlamydia trachomatis* endometritis. (A, 20X; B, 40X) Histological hematoxylin and eosin-stained endometrial tissue sections from a biopsy taken from a woman positive for *C. trachomatis* in their endometrium. White arrows indicate plasma cells; white arrowheads indicate neutrophils. (C, 20X; D, 40X) Endometrial biopsy section from a woman who was negative for endometrial infection but tested positive for *C. trachomatis* infection at their cervix.

### Key transcriptional differences in Ct endometritis highlight induction of innate and adaptive immune responses, EMT, and cell cycle dysregulation that is confirmed by functional analysis of miRNA-mRNA interactions

Since PCA showed minimal variance between the Endo- and Uninfected groups, their transcriptional data were combined and compared to the Endo+ group. A total of 2,000 mRNAs were significantly DE between Endo+ and Endo-/Uninfected groups (FDR < 0.05), with 1046 upregulated, and 954 downregulated genes in the Endo+ group. The top 100 upregulated and downregulated genes with FDR <0.05 are listed in **Supplementary Tables 3 and 4**, respectively and displayed in a volcano plot (**Figure 4**).

**Figure 4.**
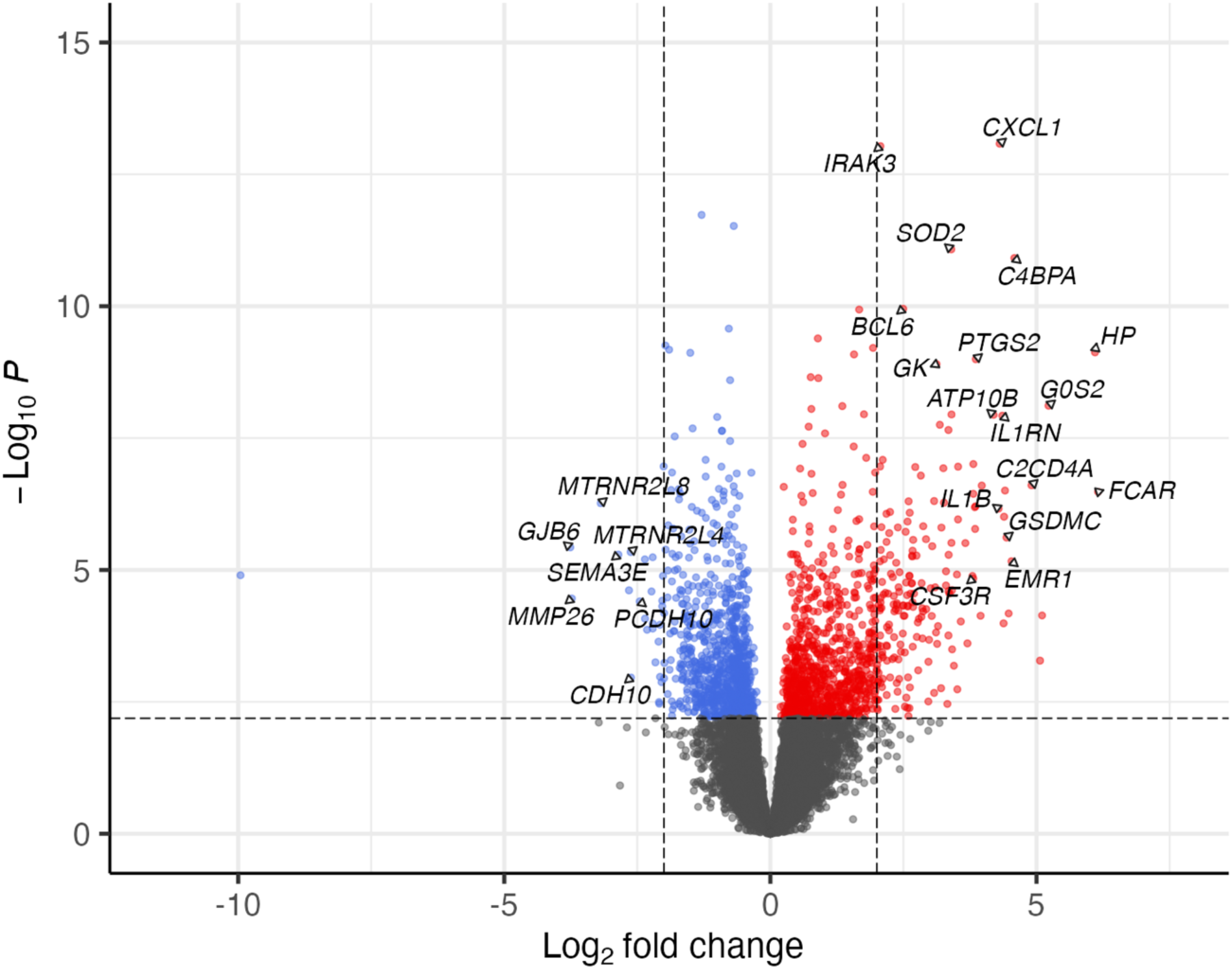
The significance and fold change of DE mRNAs (FDR<0.05) in Endo+ versus Endo-/Uninfected biopsies depicted by volcano plot. Each dot represents one mRNA gene, with red indicating upregulation and blue indicating downregulation in the Endo+ group compared to the Endo-/Uninfected group. Horizontal dashed line indicates FDR=0.05 (-log_10_P=2.2). Left vertical dashed line indicates log2 fold change=-2 (fold change= 0.25) and right vertical dashed line indicates log2 fold change=2 (fold change=4).

Several of the most strongly upregulated genes play crucial roles in local antibody production and germinal center formation. These include transcription factors and coactivators such as *BCL6* and *POU2AF1*, *CXCL13*, a key germinal center chemokine, and antibody Fc receptors including *FCAR*, the IgA receptor, and *FCRL2*, which supports B cell antibody production (**Supplementary Table 3**). This gene expression pattern was consistent with the marked plasma cell infiltrates observed in women with PID, underscoring the hallmark histology of chronic endometritis (**Figure 3A-B**) (36–40). Other highly upregulated genes are involved in the acute phase response and cytokine cascade along with secondary mediators, including chemokines, colony-stimulating factors, and prostaglandins, which amplify leukocyte recruitment and local innate immunity (*IL1B*, *IL1RN*, *IRAK3*, *GSDMC*, *C2CD4A*, *CSF3R*, *CXCL1*, *SOD2*, *PTGS2*) (**Supplementary Table 3**). *PTGS2* which encodes a cyclooxygenase enzyme that catalyzes prostaglandin synthesis and is linked to pain, was up-regulated in Endo+ women, some of whom exhibited PID symptoms.

Upregulated pathways identified by IPA included non-specific inflammatory responses (acute phase response, PPARa/RXRa activation, *IL-6*, *PI3K/AKT*, *JAK/STAT*), innate (*TREM1*, NK cell, nitrous oxide, phagosome formation, Toll-like receptor (TLR) and pattern recognition receptor (PRR) signaling, granulocyte adhesion, and HMGB-1 signaling), and adaptive immune response pathways (*IL-10*, Th1/Th2, CD40 signaling, IL-9 signaling) (**Figure 5A , Supplementary Table 5**).

**Figure 5.**
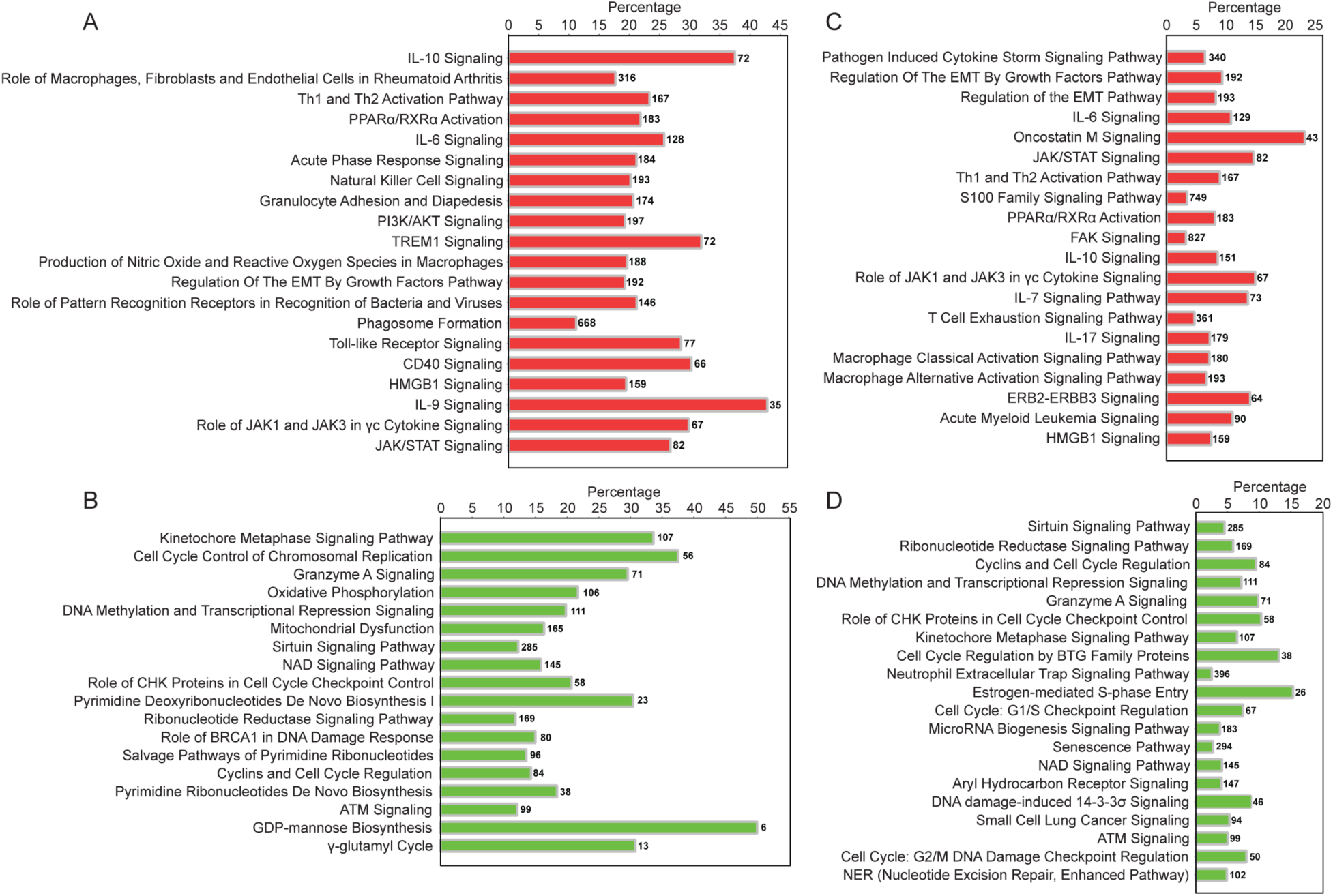
Ingenuity canonical pathways enriched by DE mRNAs and miRNAs in Endo+ compared to Endo-/Uninfected biopsies. (A) Top 20 signaling pathways enriched by significantly upregulated mRNAs in Endo+ biopsies. (B) All 18 pathways enriched by significantly downregulated mRNAs in Endo+ biopsies. (C) Top 20 signaling pathways enriched by downregulated miRNA-upregulated mRNA pairs in Endo+ biopsies. (D) Top 20 signaling pathways enriched by significantly upregulated miRNA-downregulated mRNA pairs in Endo+ biopsies. The percentage is the number of the DE genes present in each pathway, divided by the total number of genes in that pathway according to the IPA database, as listed on the right-hand side of each figure panel.

IPA also identified EMT pathway enrichment, driven by upregulation of *TGFB2*, *TGFBR2*, *SMAD3*, and *ZEB2* (**Figure 5A , Supplementary Table 5)**. *TGFB2* interacts with its receptor to activate SMAD proteins, including SMAD3. This activation promotes the expression of EMT-related genes such as *ZEB2*, a transcription factor which represses epithelial markers like E-cadherin and enhances mesenchymal markers, playing a pivotal role in EMT progression.

Downregulated genes included membrane adhesion components (*GJB6*, *CDH10*, *PCDH10*) and tissue remodeling enzymes (*MMP26*, *FBN3*, *PLOD1*), as well as genes involved in transcription regulation (*RXRG* and *SPDEF*), protein synthesis (*DHFR*, *GSTZ1*) and modification (*APRT*, *RYCR1*), and cellular metabolism (*KMO*, *NDUFB7*, *UQCRC1*) (Supplementary Table 4). Multiple genes related to histone metabolism were also decreased, potentially altering chromatin structure and impacting gene expression regulation (**Figure 4 and Supplementary Tables 4 and 6**). Downregulated pathways were enriched in genes involved with cell cycle regulation, mitochondrial dysfunction, and nucleotide biosynthesis (**Figure 5B , Supplementary Table 6**).

Consistent with the DE mRNA results (**Figure 5A**), functional enrichment analysis using IPA revealed that downregulated miRNA-mRNA interactions in Endo+ women enriched pathways such as "Regulation of EMT by Growth Factors" (**Figure 5C**, **Supplementary Table 7**), highlighting EMT promotion. Additionally, these interactions enriched pathways containing upregulated innate and adaptive immune signaling genes (**Figure 5C**). In contrast, upregulated miRNAs associated with downregulated mRNAs were enriched in pathways involving cell cycle regulation and DNA damage signaling (**Figure 5D**), aligning with mRNA findings (**Figure 5B and Supplementary Table 8**). These results suggest that Ct infection decreases the transcription of miRNAs that normally suppress EMT and immune signaling genes, potentially accelerating scarring and inflammation. Simultaneously, the infection increases the transcription of miRNAs that downregulate mRNAs involved in mucosal epithelial homeostasis, possibly further impairing uterine healing processes.

### Differential expression of miRNAs highlights EMT promotion and Treg/Th17 modulation in endometrial Ct infection

A total of 89 miRNAs were significantly DE (FDR<0.05) between Endo+ and Endo-women. Among these, 47 miRNAs were upregulated, and 42 were downregulated (**Supplementary Tables 9 and 10 ).** miRNAs with fold changes >2 and <1/2 are depicted in **Figure 6A**. The majority of (53 of 89) DE miRNAs were associated with EMT with 32 miRNAs that normally dampen EMT being downregulated and 4 that increase EMT being upregulated (**Figure 6B**), leading to an overall effect of enhancing EMT (**Table 1**).

**Figure 6.**
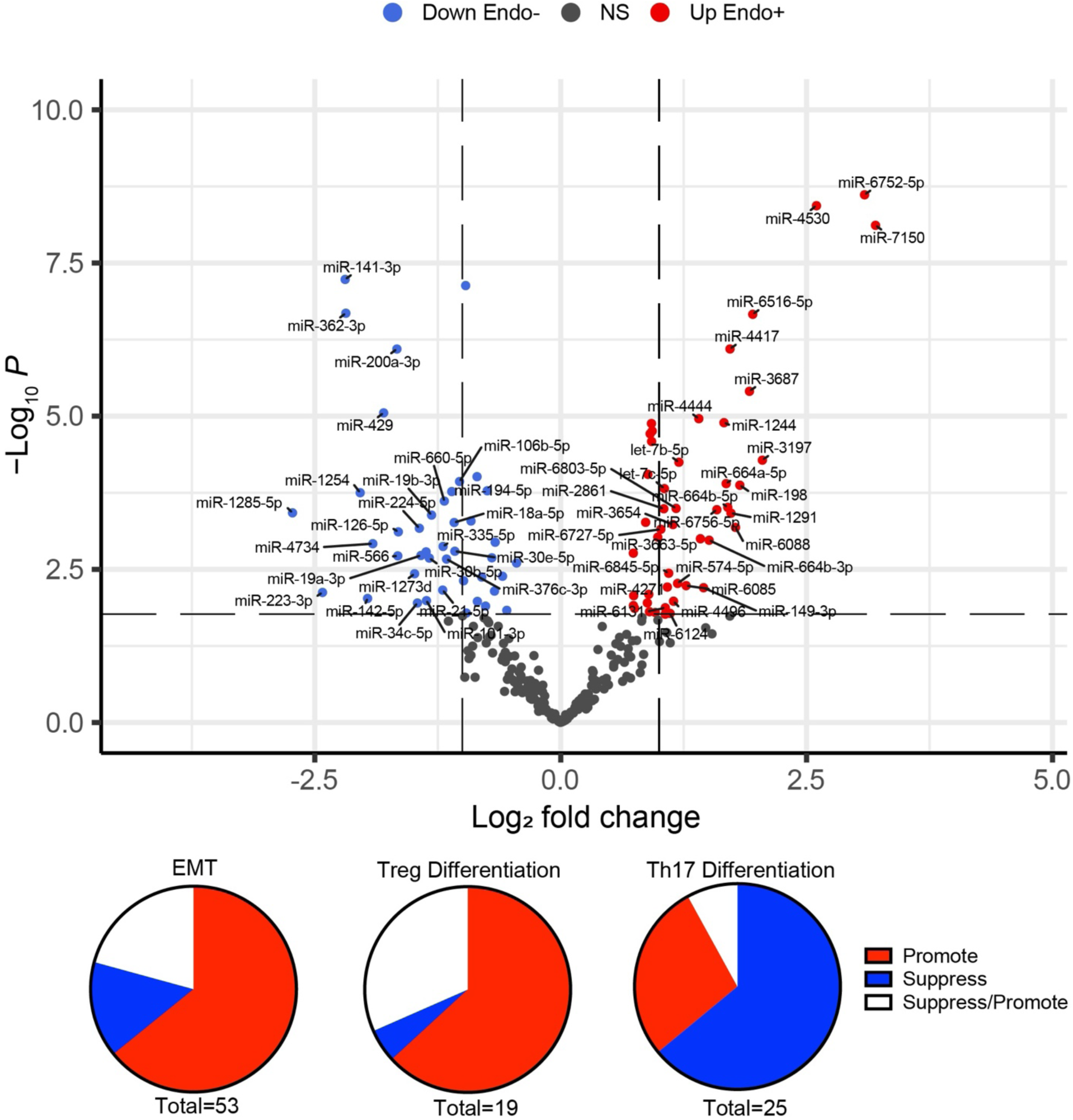
The significance and fold change of DE miRNAs (FDR<0.05) in Endo+ versus Endo-/Uninfected biopsies depicted by (A) volcano plot. Each dot represents one miRNA gene with colors indicating directions of dysregulated expression in the Endo+ group compared to the Endo-/Uninfected group. Horizontal dashed line indicates FDR=0.05 ( -log_10_P=2.2). Left vertical dashed line indicates log2 fold change=-1 (fold changes= 0.5) and right vertical dashed line indicates log2 fold change=1 (fold change=2). All significant DE miRNAs (FDR<0.05) with fold changes>2 or <0.5 are highlighted. **DE miRNAs linked to (B) EMT, (C) Tregs, and (D) Th17 differentiation are depicted by pie charts.** Red, blue and white colors indicate overall enhancing, suppressing, and enhancing/suppressing effects of miRNAs, respectively, in Endo+ women compared to Endo-/Uninfected women.

**Table 1.** DE miRNAs in Endo+ compared to Endo-associated with EMT.

| DE miRNAs | Fold Change | P value | FDR | Expression changes in Endo+ compared to Endo- | Function of miRNAs | Overall effects | Citations |
| --- | --- | --- | --- | --- | --- | --- | --- |
| miR-566 | 0.32 | 0.002 | 0.009 | Decrease (-) | ↓EMT | **Promote EMT** | (41) |
| miR-200a-3p | 0.32 | <0.001 | <0.001 | Decrease (-) | ↓EMT |  | (42) |
| miR-200c-3p | 0.59 | 0.013 | 0.039 | Decrease (-) | ↓EMT |  | (42) |
| miR-141-3p | 0.22 | <0.001 | <0.001 | Decrease (-) | ↓EMT |  | (43) |
| miR-429 | 0.29 | <0.001 | <0.001 | Decrease (-) | ↓EMT |  |  |
| miR-376a-3p | 0.51 | 0.016 | 0.048 | Decrease (-) | ↓EMT |  | (44) |
| miR-376c-3p | 0.45 | 0.002 | 0.009 | Decrease (-) | ↓EMT |  | (45) |
| miR-374b-5p | 0.39 | 0.002 | 0.008 | Decrease (-) | ↓EMT |  | (46) |
| miR-30b-5p | 0.4 | 0.002 | 0.009 | Decrease (-) | ↓EMT |  | (47) |
| miR-30c-5p | 0.68 | 0.015 | 0.045 | Decrease (-) | ↓EMT |  | (47) |
| miR-30e-5p | 0.47 | 0.002 | 0.008 | Decrease (-) | ↓EMT |  | (47) |
| miR-186-5p | 0.62 | 0.002 | 0.009 | Decrease (-) | ↓EMT |  | (48) |
| miR-130a-3p | 0.63 | 0.001 | 0.006 | Decrease (-) | ↓EMT |  | (49) |
| miR-335-5p | 0.44 | 0.001 | 0.007 | Decrease (-) | ↓EMT |  | (50) |
| miR-17-5p | 0.6 | <0.001 | 0.002 | Decrease (-) | ↓EMT |  | (51) |
| miR-18a-5p | 0.47 | 0.001 | 0.004 | Decrease (-) | ↓EMT |  | (51) |
| miR-19a-3p | 0.37 | 0.002 | 0.009 | Decrease (-) | ↓EMT |  | (51) |
| miR-19b-3p | 0.4 | <0.001 | 0.003 | Decrease (-) | ↓EMT |  | (51) |
| miR-20a-5p | 0.5 | 0.005 | 0.019 | Decrease (-) | ↓EMT |  | (51) |
| miR-20b-5p | 0.51 | <0.001 | <0.001 | Decrease (-) | ↓EMT |  | (52, 53) |
| miR-34c-5p | 0.36 | 0.011 | 0.036 | Decrease (-) | ↓EMT |  | (54) |
| miR-101-3p | 0.39 | 0.01 | 0.035 | Decrease (-) | ↓EMT |  | (55) |
| miR-194-5p | 0.46 | <0.001 | 0.002 | Decrease (-) | ↓EMT |  | (56) |
| miR-193a-3p | 0.57 | 0.004 | 0.017 | Decrease (-) | ↓EMT |  | (57) |
| miR-148a-3p | 0.53 | 0.001 | 0.004 | Decrease (-) | ↓EMT |  | (58, 59) |
| mir-148b-3p | 0.73 | 0.002 | 0.011 | Decrease (-) | ↓EMT |  | (60) |
| miR-126-5p | 0.32 | <0.001 | 0.004 | Decrease (-) | ↓EMT |  | (61, 62) |
| miR-142-5p | 0.26 | 0.009 | 0.032 | Decrease (-) | ↓EMT |  | (63) |
| miR-1285-5p | 0.15 | <0.001 | 0.003 | Decrease (-) | ↓EMT |  | (64) |
| mir-1254 | 0.24 | <0.001 | 0.002 | Decrease (-) | ↓EMT |  | (65) |
| miR-6803-5p | 2.26 | <0.001 | 0.003 | Increase (+) | ↑EMT |  | (66) |
| miR-664a-3p | 1.82 | 0.001 | 0.004 | Increase (+) | ↑EMT |  | (67) |
| miR-663b | 1.98 | 0.001 | 0.005 | Increase (+) | ↑ EMT |  | (68) |
| miR-197-3p | 1.67 | 0.012 | 0.039 | Increase (+) | ↑ EMT |  | (69) |
| miR-106b-5p | 0.49 | <0.001 | 0.001 | Decrease (-) | ↑ EMT | Suppress EMT | (70-72) |
| miR-198 | 3.54 | <0.001 | 0.001 | Increase (+) | ↓ EMT |  | (73) |
| miR-30a-3p | 1.67 | 0.008 | 0.027 | Increase (+) | ↓ EMT |  | (74) |
| miR-320a | 1.9 | <0.001 | <0.001 | Increase (+) | ↓ EMT |  | (75) |
| miR-320c | 1.88 | <0.001 | <0.001 | Increase (+) | ↓ EMT |  | (76) |
| miR-320d | 1.85 | <0.001 | 0.001 | Increase (+) | ↓ EMT |  | (77) |
| let-7c-5p | 2.08 | <0.001 | 0.002 | Increase (+) | ↓ EMT |  | (78) |
| miR-4429 | 1.9 | <0.001 | <0.001 | Increase (+) | ↓ EMT |  | (79, 80) |
| miR-130b-3p | 0.63 | 0.007 | 0.026 | Decrease (-) | ↓ ↑ EMT | Suppress/Promote EMT | (81-83) |
| miR-21-5p | 0.6 | 0.024 | 0.05 | Decrease (-) | ↓ ↑ EMT |  | (84, 85) |
| miR-27a-3p | 0.55 | <0.001 | 0.001 | Decrease (-) | ↓ ↑ EMT |  | (86, 87) |
| miR-27b-3p | 0.66 | 0.004 | 0.017 | Decrease (-) | ↓ ↑ EMT |  | (88, 89) |
| miR-224-5p | 0.37 | 0.001 | 0.004 | Decrease (-) | ↓ ↑ EMT |  | (90, 91) |
| miR-455-5p | 0.56 | 0.011 | 0.035 | Decrease (-) | ↓ ↑ EMT |  | (92, 93) |
| miR-223-3p | 0.19 | 0.008 | 0.027 | Decrease (-) | ↓ ↑ EMT |  | (94) |
| miR-149-3p | 2.74 | 0.006 | 0.024 | Increase (+) | ↓ ↑ EMT |  | (95, 96) |
| miR-574-5p | 2.28 | 0.005 | 0.021 | Increase (+) | ↓ ↑ EMT |  | (97, 98) |
| miR-146a-5p | 2.09 | 0.017 | 0.048 | Increase (+) | ↓ ↑ EMT |  | (99, 100) |
| miR-320b | 1.91 | <0.001 | <0.001 | Increase (+) | ↓ ↑ EMT |  | (75, 101) |

IPA’s target filter and literature review identified target mRNAs linked to TGF-β signaling, crucial for EMT induction. These included *JAK1/STAT3* (103), Smad-dependent and (104) independent *TGF-β* pathways, and the autocrine signaling network for EMT involving *TGF-β* and *ZEB2* (104). Transcription factors (e.g., *ZEB2*, *FOXO1*) and pathways like PI3K-AKT were upregulated, emphasizing EMT promotion. Among the 25 most significantly downregulated miRNAs in Endo+ samples, seven (miR-141-3p, miR-200a-3p, miR-106b-5p, miR-224-5p, miR-20a-5p, miR-21-5p, and miR-223-3p) were identified through literature review as directly suppressing *TGF-β* production. Consequently, their downregulation in Endo+ women likely enhances *TGF-β* activity, which may suppress chronic inflammation but also promote EMT, potentially contributing to adverse pathological outcomes.

Among the 89 differentially expressed (DE) miRNAs, 32 were associated with T cell regulation, including subclasses such as Tregs, Th17, and Th1. *TGF-β* plays a key role in the differentiation of both Tregs and Th17 cells. However, the presence of additional cytokines, particularly interleukin-6 (*IL-6*), shifts differentiation toward Th17 cells. Specifically, low *TGF-β* concentrations combined with *IL-6* promote Th17 differentiation, whereas high *TGF-β* levels suppress *IL-6* signaling and favor Treg differentiation (105).

Twenty miRNAs were linked to Treg differentiation, with 12 downregulated miRNAs typically functioning to suppress Tregs (**Figure 6C**, **Table 2**). Notably, miR-146a-5p, which promotes Tregs, was upregulated, while miR-148a-3p, another Treg-enhancing miRNA, was reduced. These expression patterns suggest an overall increase in Treg differentiation in Endo+ women.

For Th17 differentiation, 25 miRNAs were identified (**Figure 6D**, **Table 3**). Of these, 14 miRNAs that typically enhance Th17 cells were significantly reduced, 10 of which also function to suppress Tregs. Additionally, two miRNAs that normally inhibit Th17 differentiation were upregulated. Conversely, six miRNAs that suppress Th17 differentiation were downregulated, and miR-149-3p, which promotes Th17 cells, was increased. Overall, 16 of the 25 DE miRNAs are predicted to reduce Th17 differentiation, while 7 are associated with increased Th17 differentiation.

**Table 3.**
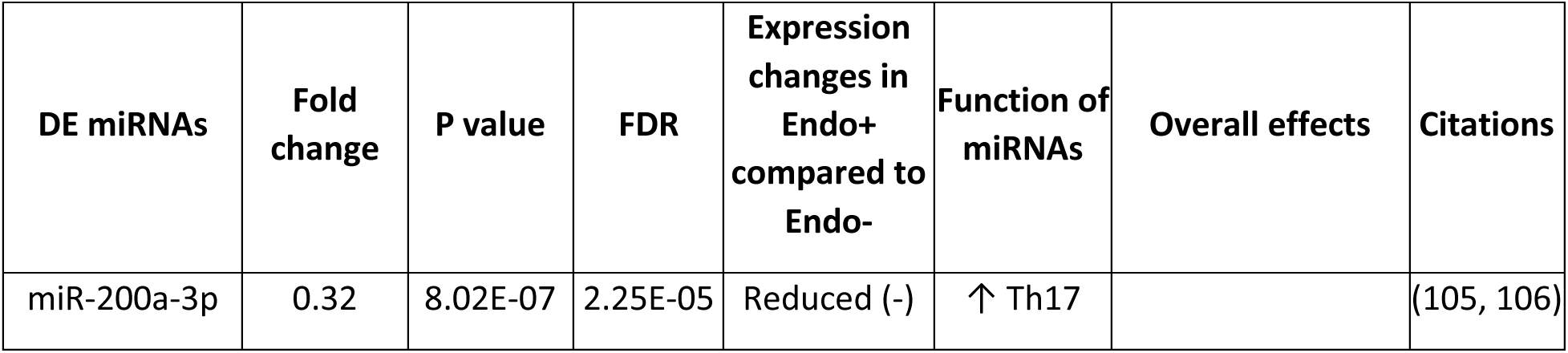
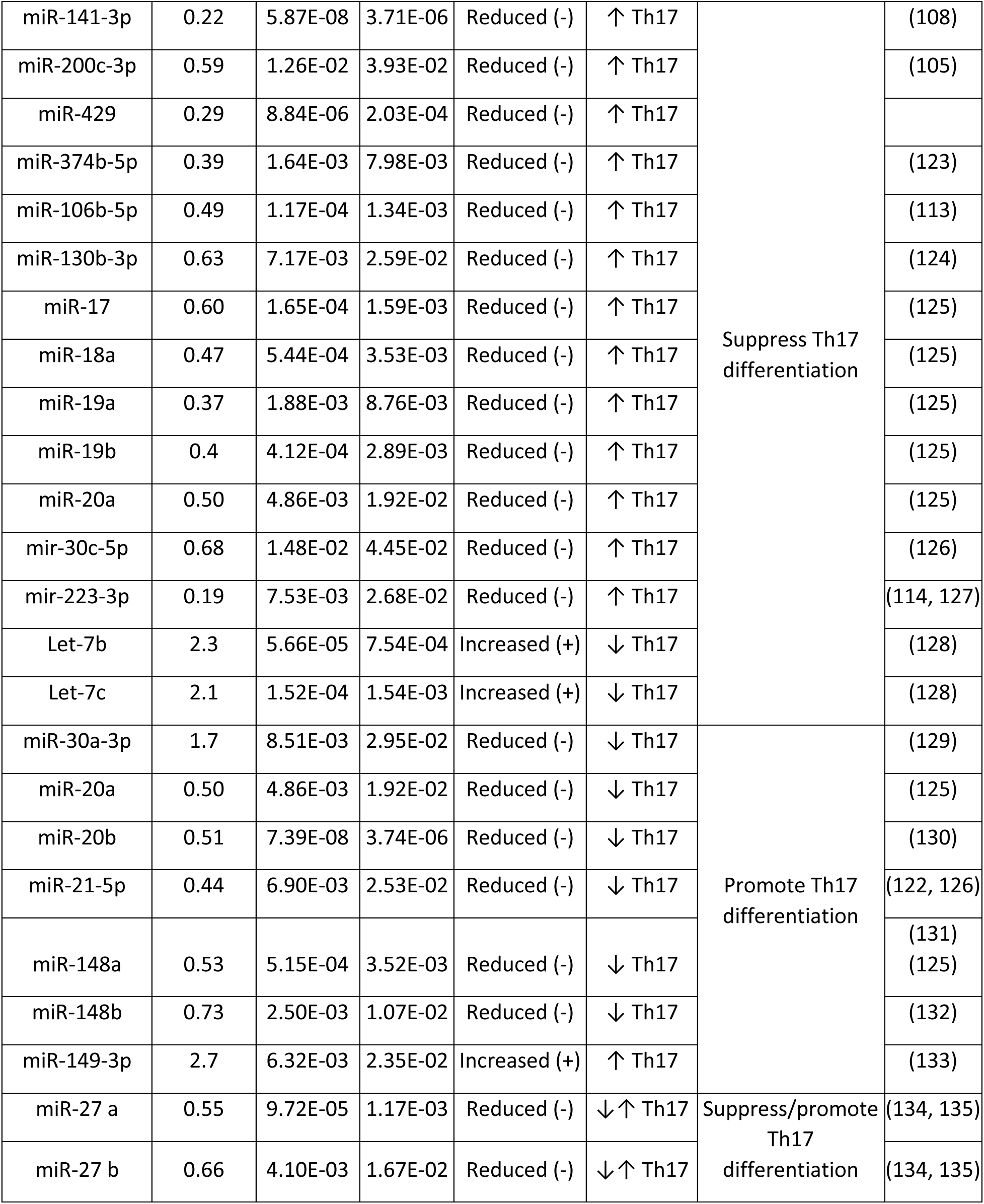
DE miRNAs in Endo+ compared to Endo-, associated with Th17 cell differentiation.

| DE miRNAs | Fold change | P value | FDR | Expression changes in Endo+ compared to Endo- | Function of miRNAs | Overall effects | Citations |
| --- | --- | --- | --- | --- | --- | --- | --- |
| miR-200a-3p | 0.32 | 8.02E-07 | 2.25E-05 | Reduced (-) | ↑ Th17 |  | (105, 106) |
| miR-141-3p | 0.22 | 5.87E-08 | 3.71E-06 | Reduced (-) | ↑ Th17 | Suppress Th17 differentiation | (108) |
| miR-200c-3p | 0.59 | 1.26E-02 | 3.93E-02 | Reduced (-) | ↑ Th17 |  | (105) |
| miR-429 | 0.29 | 8.84E-06 | 2.03E-04 | Reduced (-) | ↑ Th17 |  |  |
| miR-374b-5p | 0.39 | 1.64E-03 | 7.98E-03 | Reduced (-) | ↑ Th17 |  | (123) |
| miR-106b-5p | 0.49 | 1.17E-04 | 1.34E-03 | Reduced (-) | ↑ Th17 |  | (113) |
| miR-130b-3p | 0.63 | 7.17E-03 | 2.59E-02 | Reduced (-) | ↑ Th17 |  | (124) |
| miR-17 | 0.60 | 1.65E-04 | 1.59E-03 | Reduced (-) | ↑ Th17 |  | (125) |
| miR-18a | 0.47 | 5.44E-04 | 3.53E-03 | Reduced (-) | ↑ Th17 |  | (125) |
| miR-19a | 0.37 | 1.88E-03 | 8.76E-03 | Reduced (-) | ↑ Th17 |  | (125) |
| miR-19b | 0.4 | 4.12E-04 | 2.89E-03 | Reduced (-) | ↑ Th17 |  | (125) |
| miR-20a | 0.50 | 4.86E-03 | 1.92E-02 | Reduced (-) | ↑ Th17 |  | (125) |
| mir-30c-5p | 0.68 | 1.48E-02 | 4.45E-02 | Reduced (-) | ↑ Th17 |  | (126) |
| mir-223-3p | 0.19 | 7.53E-03 | 2.68E-02 | Reduced (-) | ↑ Th17 |  | (114, 127) |
| Let-7b | 2.3 | 5.66E-05 | 7.54E-04 | Increased (+) | ↓ Th17 |  | (128) |
| Let-7c | 2.1 | 1.52E-04 | 1.54E-03 | Increased (+) | ↓ Th17 |  | (128) |
| miR-30a-3p | 1.7 | 8.51E-03 | 2.95E-02 | Reduced (-) | ↓ Th17 | Promote Th17 differentiation | (129) |
| miR-20a | 0.50 | 4.86E-03 | 1.92E-02 | Reduced (-) | ↓ Th17 |  | (125) |
| miR-20b | 0.51 | 7.39E-08 | 3.74E-06 | Reduced (-) | ↓ Th17 |  | (130) |
| miR-21-5p | 0.44 | 6.90E-03 | 2.53E-02 | Reduced (-) | ↓ Th17 |  | (122, 126) |
| miR-148a | 0.53 | 5.15E-04 | 3.52E-03 | Reduced (-) | ↓ Th17 |  | (131)<br>(125) |
| miR-148b | 0.73 | 2.50E-03 | 1.07E-02 | Reduced (-) | ↓ Th17 |  | (132) |
| miR-149-3p | 2.7 | 6.32E-03 | 2.35E-02 | Increased (+) | ↑ Th17 |  | (133) |
| miR-27 a | 0.55 | 9.72E-05 | 1.17E-03 | Reduced (-) | ↓ ↑ Th17 | Suppress/promote Th17 differentiation | (134, 135) |
| miR-27 b | 0.66 | 4.10E-03 | 1.67E-02 | Reduced (-) | ↓ ↑ Th17 |  | (134, 135) |

Several target mRNAs were linked to promoting Treg differentiation over Th17 cells. These include transcription factors such as *FOXO1*, *RUNX1*, *IRF1*, and *BATF*, as well as key *TGF-β* pathway genes (*TGFBR2*, *SMAD3*, *STAT5*, *STAT3*, and *IL-10R*). Collectively, these findings indicate a shift toward increased Treg activity and reduced Th17 differentiation. This shift may help limit tissue-damaging neutrophil responses but could come at the cost of impaired Th17/Th1-mediated protection against reinfection (7).

A within-sample group clustering heatmap of DE miRNAs (**Figure 7**) revealed distinct expression patterns across groups: Endo+ women with and without symptoms, Endo-, Uninfected, and women with non-STI-induced pelvic pain. One cluster of miRNAs was downregulated in Endo+ women (symptomatic and asymptomatic) compared to Endo-, Uninfected, and women with non-STI-induced pelvic pain. Literature review suggests these miRNAs typically suppress EMT (**Table 1**), implying their downregulation in Endo+ women may promote EMT, regardless of symptom presence. Notably, this downregulation was not observed in non-STI pelvic pain cases, suggesting a link to active Ct infection (**Figure 7**).

**Figure 7.**
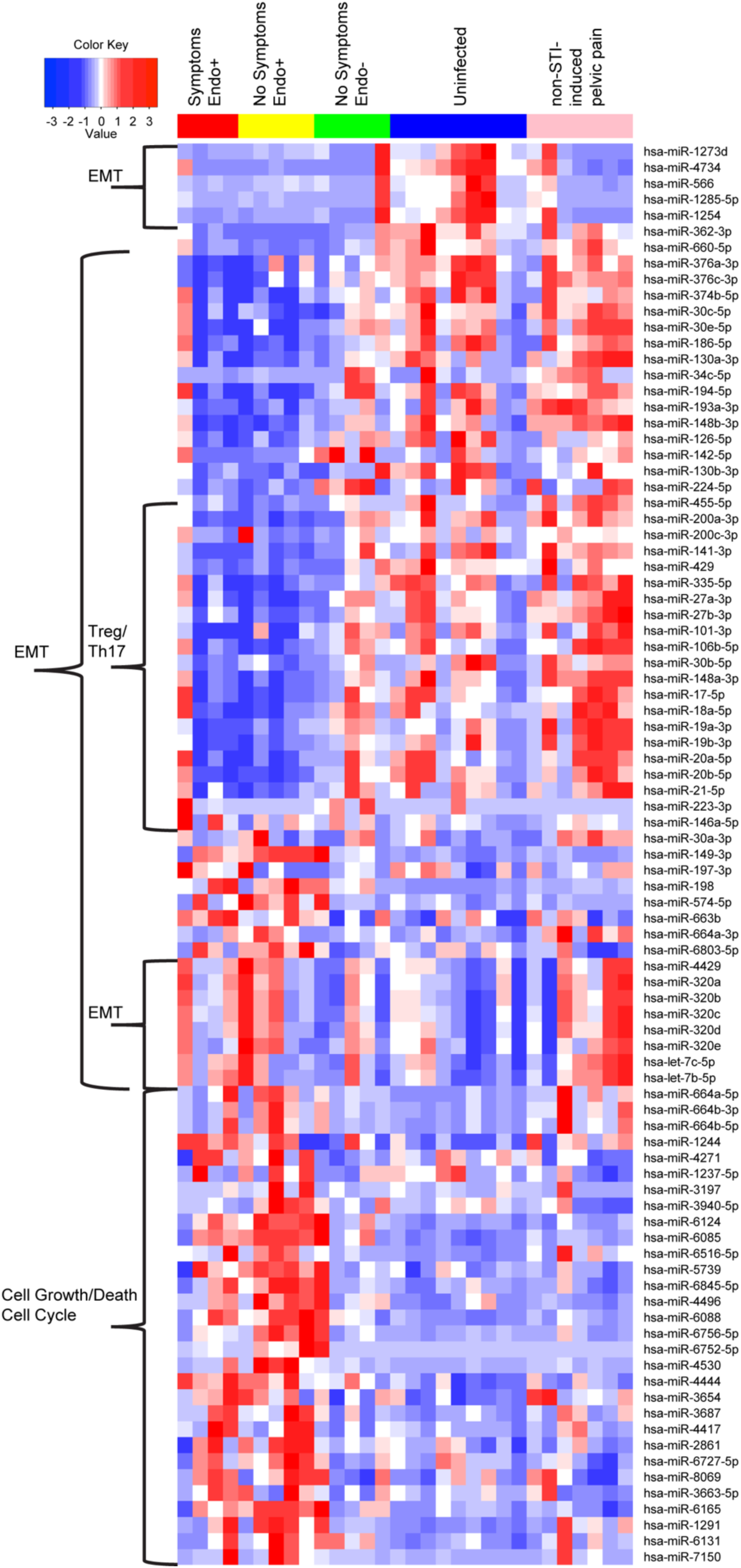
A within-sample group hierarchical clustering heatmap of DE miRNAs revealed distinct expression patterns across groups. Heatmap columns indicate individual patients with red: Endo+ with pelvic pain; yellow: Endo+ without symptoms; green: Endo-without symptoms; blue: uninfected without symptoms; pink: pelvic pain without sexually transmitted infection (STI). Heatmap rows indicate genes. Blue: low expression miRNAs, white: intermediately expressed miRNAs, and red: highly expressed miRNAs.

However, a minor cluster of miRNAs (mir-1273, mir-1285-5p, mir-1254, mir-4734, and mir-566) that normally dampen mRNAs promoting EMT (**Table 1**, **Figure 7**), was downregulated in Endo+, 4 of 5 Endo-, and 5 of 7 non-STI pelvic pain cases with recent prior Ct infection, indicating potential enhancement of EMT processes irrespective of current Ct ascension or symptom status. Each of these miRNAs have been studied in various cancer types, and reportedly have both positive (miR-1273 (137), miR-4734) and negative roles (miR-1285-5p (138), miR-1254 (139), miR-566 (140)) in promoting cell proliferation, invasion, and metastasis. Furthermore, four miRNAs (miR-320b (102, 141), let-7c-5p and let-7b-5p (79)) reported to induce EMT and promote oncogenesis were upregulated in Endo+ women and those with non-STI-induced pelvic pain (**Table 1**, **Figure 7**).

### Distinct immune and fibrosis-related pathways were detected in endometrial vs. blood mRNA profiles in women with chlamydial endometritis

In a previous study (32), we compared blood-derived mRNA profiles from women with Ct-induced pelvic inflammatory disease (PID) (Endo+) to those from asymptomatic women with cervical infection only and uninfected women (Endo-/Uninfected). A comparison of the top enriched pathways identified in blood with those found in the endometrium of Endo+ women revealed notable differences (**Table 4**). Shared pathways related to innate immune responses, including granulocyte adhesion, *TREM1* signaling, Toll-like receptor (TLR) signaling, integrin signaling, *IL-1*, *IL-10*, and *IL-6* response pathways, as well as *JAK/STAT* signaling. Endometrium--specific responses included natural killer (NK) cell signaling pathways, *IL-23*, *STAT3*, and Th17 activation pathways which can lead to chronic inflammation if excessively activated (142), multiple adaptive T cell response pathways (Th1/Th17, Th2, and *IL-7*), and fibrosis-related pathways, *TGF-β* signaling and EMT. These sharply contrasted with the blood profiles of Endo+ women, where TCR signaling pathways—including mTOR, which integrates immune signals and metabolic cues for T cell maintenance and activation, and *ICOS*-*ICOSL* signaling which delivers the co-stimulatory signals that promote T cell activation and differentiation—were significantly downregulated, and activation of EMT pathways was undetected. Conversely, type I interferon signaling pathways were prominent in blood-borne profiles, but displayed minimal enrichment in the endometrium.

**Table 4.**
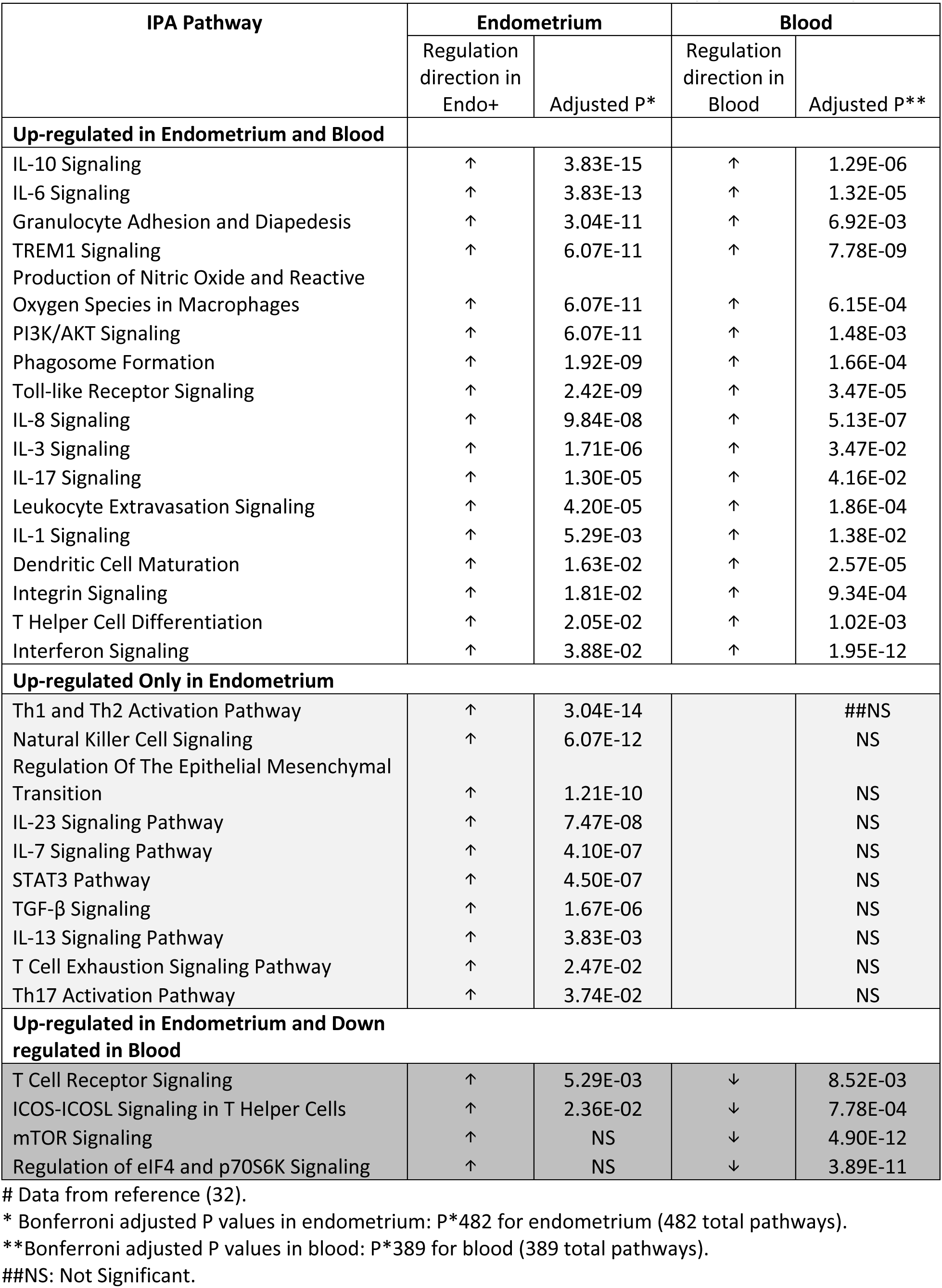
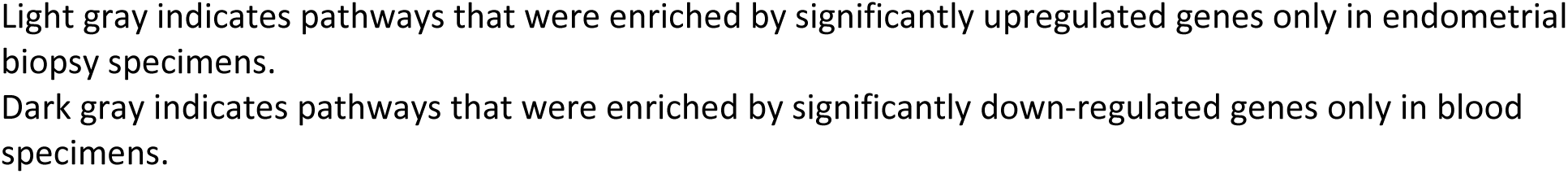
Comparison of pathways enriched by significantly differentially expressed (DE) mRNAs in Endo+ versus Endo-women, as determined from either endometrial biopsy or blood samples#.

|  | Regulation<br>direction in<br>Endo+ | Adjusted P* | Regulation<br>direction in<br>Blood | Adjusted P** |
| --- | --- | --- | --- | --- |
| **Up-regulated in Endometrium and Blood** |  |  |  |  |
| IL-10 Signaling | ↑ | 3.83E-15 | ↑ | 1.29E-06 |
| IL-6 Signaling | ↑ | 3.83E-13 | ↑ | 1.32E-05 |
| Granulocyte Adhesion and Diapedesis | ↑ | 3.04E-11 | ↑ | 6.92E-03 |
| TREM1 Signaling | ↑ | 6.07E-11 | ↑ | 7.78E-09 |
| Production of Nitric Oxide and Reactive<br>Oxygen Species in Macrophages | ↑ | 6.07E-11 | ↑ | 6.15E-04 |
| PI3K/AKT Signaling | ↑ | 6.07E-11 | ↑ | 1.48E-03 |
| Phagosome Formation | ↑ | 1.92E-09 | ↑ | 1.66E-04 |
| Toll-like Receptor Signaling | ↑ | 2.42E-09 | ↑ | 3.47E-05 |
| IL-8 Signaling | ↑ | 9.84E-08 | ↑ | 5.13E-07 |
| IL-3 Signaling | ↑ | 1.71E-06 | ↑ | 3.47E-02 |
| IL-17 Signaling | ↑ | 1.30E-05 | ↑ | 4.16E-02 |
| Leukocyte Extravasation Signaling | ↑ | 4.20E-05 | ↑ | 1.86E-04 |
| IL-1 Signaling | ↑ | 5.29E-03 | ↑ | 1.38E-02 |
| Dendritic Cell Maturation | ↑ | 1.63E-02 | ↑ | 2.57E-05 |
| Integrin Signaling | ↑ | 1.81E-02 | ↑ | 9.34E-04 |
| T Helper Cell Differentiation | ↑ | 2.05E-02 | ↑ | 1.02E-03 |
| Interferon Signaling | ↑ | 3.88E-02 | ↑ | 1.95E-12 |
| **Up-regulated Only in Endometrium** |  |  |  |  |
| Th1 and Th2 Activation Pathway | ↑ | 3.04E-14 |  | ##NS |
| Natural Killer Cell Signaling | ↑ | 6.07E-12 |  | NS |
| Regulation Of The Epithelial Mesenchymal<br>Transition | ↑ | 1.21E-10 |  | NS |
| IL-23 Signaling Pathway | ↑ | 7.47E-08 |  | NS |
| IL-7 Signaling Pathway | ↑ | 4.10E-07 |  | NS |
| STAT3 Pathway | ↑ | 4.50E-07 |  | NS |
| TGF-β Signaling | ↑ | 1.67E-06 |  | NS |
| IL-13 Signaling Pathway | ↑ | 3.83E-03 |  | NS |
| T Cell Exhaustion Signaling Pathway | ↑ | 2.47E-02 |  | NS |
| Th17 Activation Pathway | ↑ | 3.74E-02 |  | NS |
| **Up-regulated in Endometrium and Down**<br>**regulated in Blood** |  |  |  |  |
| T Cell Receptor Signaling | ↑ | 5.29E-03 | ↓ | 8.52E-03 |
| ICOS-ICOSL Signaling in T Helper Cells | ↑ | 2.36E-02 | ↓ | 7.78E-04 |
| mTOR Signaling | ↑ | NS | ↓ | 4.90E-12 |
| Regulation of eIF4 and p70S6K Signaling | ↑ | NS | ↓ | 3.89E-11 |

## Discussion

The findings from this study improve understanding of how Ct infection modulates the molecular landscape of the human endometrium, emphasizing the interplay between immune responses, EMT, and regulatory miRNA activity. These results provide insight into how Ct infection may contribute to adverse pregnancy outcomes, including stillbirth and preterm labor, as well as the Ct-induced complication of chronic pelvic pain.

The integrated transcriptomic and miRNA analyses revealed distinct molecular signatures in the endometrial tissues of women with endometrial Ct infection (Endo+), highlighting active innate and adaptive immune signaling pathways alongside EMT promotion. The transcriptional patterns of upregulated genes associated with the acute phase response as well as genes that promote antibody-producing cells and cell-mediated immunity correlate with histological findings of endometritis, marked by a predominance of plasma cell infiltrates mixed with subepithelial neutrophils and lymphocytes. Other upregulated genes in Endo+ women, such as *TGFB2*, *TGFBR2*, and *SMAD3*, collectively drive EMT through Smad-dependent and independent mechanisms, while transcription factors like *ZEB2* suppress epithelial markers to facilitate mesenchymal transitions. The significant downregulation of EMT-suppressing miRNAs, including miR-141-3p and miR-200a-3p, further supports this shift toward EMT, aligning with prior observations in murine models (18, 19, 143). These molecular alterations are consistent with enhanced tissue remodeling, fibrosis, and scarring—hallmarks of the Ct-induced complications of tubal factor infertility and ectopic pregnancy. The additional detection of upregulated miRNAs that depress gene pathways engaged in nucleotide synthesis and cell cycle control indicate a further mechanism whereby miRNA dysregulation could inhibit healing of infected endometrial tissue. While this study focused on endometrial biopsy tissues, it is highly likely that similar mechanisms are at work in the delicate oviduct. EMT and fibrosis in the oviduct could impair critical reproductive processes, including gamete transport, fertilization, early embryo development, immune defense, and structural integrity—all of which are essential for successful reproduction.

The enrichment of *TGF-β* signaling and its downstream transcription factors, such as *ZEB2*, in Endo+ samples underscores the role of this pathway in driving EMT. Downregulated miRNAs that typically suppress *TGF-β* signaling, including miR-141-3p and miR-200a-3p, likely amplify this effect. These molecular changes are consistent with previous studies in mouse models, which demonstrated that dysregulated EMT contributes to fibrosis and oviductal scarring. Igietseme et al. (18) showed that *Chlamydia muridarum*-induced TNFα signaling and caspase activation were linked to the downregulation of miRNAs that typically inhibit EMT and fibrosis. Similarly, alterations in *TGF-β*, *TNF*, *ZEB*, and the miR-200 family observed in Endo+ women were also identified in infertile mice, with multiple differentially expressed miRNAs involved in EMT overlapping between the two studies (143, 144) demonstrated significant downregulation of miRNAs that suppress fibrosis, including members of the miR-200 family in mice with oviduct pathology following *C. muridarum* infection. This overlap in miRNA dysregulation between human and mouse studies suggests a conserved *Chlamydia*-induced EMT mechanism across species. The downregulation of these miRNAs, combined with the activation of *TGF-β* signaling, likely creates a microenvironment conducive to tissue remodeling and scarring. This process may represent a critical step in the progression from infection to tubal factor infertility.

A notable finding was the miRNA-driven shift in the Treg/Th17 balance. Among the 89 DE miRNAs, many were linked to immune cell regulation, particularly promoting Treg differentiation and function. The downregulation of miRNAs that typically suppress Tregs (e.g., miR-200a-3p, miR-374b-5p) suggests an enhanced Treg response, potentially favoring immune tolerance at the expense of effective Th1/Th17-mediated immunity. This interpretation is bolstered by the upregulation of miR-146a-5p, known to support Treg function, and the increased expression of Treg-associated mRNAs, such as TGFBR2 and Smad3.

Conversely, the downregulation of Th17-promoting miRNAs, such as miR-21-5p, miR-19b-3p, and miR-223-3p, indicates a reduction in Th17 activity. Since Th17 cells enhance neutrophil recruitment, decreased Th17 activity could mitigate tissue-damaging neutrophil responses, key drivers of chlamydial disease pathogenesis (15). However, given Ct’s intracellular developmental cycle, and its production of Chlamydial Protease Activation Factor (CPAF), which impairs neutrophil function (145), neutrophils have limited efficacy in eliminating Ct, and reduced neutrophil activation may have little impact on infection resolution. However, Th17 cells exhibit plasticity, transitioning between Th17 and Th1 phenotypes, and have been strongly associated with resistance to chlamydial reinfection (7). Thus, the observed shift in the Treg/Th17 balance has contrasting implications. Enhanced Treg/Th17 activity may reduce chronic inflammation and preserve endometrial function; while simultaenously, impeding development of protective immunity, potentially contributing to the high rates of reinfection observed in some cohort participants (7). This delicate trade-off underscores the challenges of therapeutically targeting these pathways, as interventions could disrupt the balance between immune suppression and effective defense.

Comparative analyses between endometrial and previously published blood mRNA profiles highlight tissue-specific responses to Ct infection. While blood profiles revealed activation of interferon-mediated signaling and downregulation of adaptive T cell pathwys, endometrial responses were dominated by activation of T cells (e.g., Treg, Th1, Th17), EMT, and *TGF-β*-driven immune regulation. These differences underscore the localized nature of Ct-induced endometrial inflammation and fibrosis, which may not be readily detectable through systemic biomarkers. This localized EMT induction aligns with the histological evidence of fibrosis observed in Ct-affected Fallopian tubes, suggesting a direct link between Ct gene regulation in the upper genital tract and long-term reproductive sequelae. These findings underscore the importance of localized studies for understanding the distinct molecular landscape of Ct-infected reproductive tissues.

We acknowledge several limitations of this study. The sample size for the endometrial transcriptome was constrained by the difficulty of obtaining sufficient high-quality endometrial tissue for mRNA and miRNA profiling. Additionally, the cross-sectional design of the study limits our ability to assess the temporal progression of infection and associated pathology. Future studies with larger sample sizes and longitudinal designs are needed to validate these findings and provide a more comprehensive understanding into the progression of Ct-induced pathology and the persistence of molecular changes post-infection. Extending these analyses to other sexually transmitted infections could determine whether similar miRNA-mediated mechanisms drive reproductive tract inflammation and scarring across different pathogens.

Despite these limitations, our study provides a new and comprehensive view of molecular changes in the endometrium during Ct infection, highlighting critical pathways that drive tissue remodeling, fibrosis, and immune dysregulation. The findings also raise concerns about asymptomatic Endo+ women, who exhibited similar molecular profiles to symptomatic individuals, suggesting that subclinical endometrial inflammation may contribute to reproductive morbidity, which supports prior studies indicating that asymptomatic women with Ct-induced endometritis have an increased risk for infertility (25).

In summary, this study demonstrates that endometrial Ct infection induces a miRNA-mediated regulatory network that promotes EMT and alters immune cell balance, favoring Treg differentiation over Th17 responses. These molecular changes likely contribute to tissue scarring and impaired immune protection, offering new insights into the pathogenesis of Ct-induced reproductive morbidity. Targeting these pathways may provide a foundation for innovative therapeutic strategies to mitigate the long-term consequences of Ct infection.

## Acknowledgment

We thank the women who agreed to participate in this study; Ingrid Macio, Melinda Petrina, Carol Priest, Abi Jett, and Lorna Rabe, for their efforts in the clinic and the microbiology laboratory; and the staff at the Allegheny County Health Department STD Clinic. We gratefully acknowledge the technical support from the UNC High Throughput Sequencing Facility.

## Footnotes

This work was supported by the National Institute of Allergy and Infectious Diseases U19AI144181 and R01AI170959.

