## Supplemental Table 1-10 for "miRNA/mRNA analysis of increased TGF-β pathways drive epithelial-mesenchymal transition and regulatory T cell differentiation": Supp Table 1.docx

| **Supp Table 1. Characteristics of the individuals from which samples were obtained for mRNA data analysis.** | | | | | | |
| --- | --- | --- | --- | --- | --- | --- |
| **Characteristics** | **Number (%)** | | | | | **P value** |
|  |  | **Endo+ (N=7)** | | **Endo- (N=8)** | |  |
|  | **Total** | **Symptom+** | **Symptom-** | **Cervix+** | **Uninfected** |  |
| **Number of women per group** | 15 | 4 | 3 | 4 | 4 |  |
| **Age, y, median (CI)** | 21  (19-24.9) | 22.5  (18-27) | 21  (20-22) | 25.5  (19-33) | 20  (18-22) | 0.47 |
| **Race/Ethnicity** |  |  |  |  |  | 0.31 |
| African American | 11  (73.3%) | 4  (100%) | 1  (33.3%) | 2  (50%) | 4  (100%) |  |
| White | 2  (13.3%) | 0 | 1  (33.3%) | 1  (25%) | 0 |  |
| Multiracial | 2  (13.3%) | 0 | 1  (33.3%) | 1  (25%) | 0 |  |
| **Marital status** |  |  |  |  |  | 1.00 |
| Single | 14 (93.3%) | 4  (100%) | 3  (100%) | 4  (100%) | 3  (75%) |  |
| Living with partner | 1  (6.7%) | 0 | 0 | 0 | 1  (25%) |  |
| **Education** |  |  |  |  |  | 0.39 |
| Less than high school graduate | 2  (13.3%) | 0 | 0 | 0 | 2  (50%) |  |
| High school graduate or GED degree | 3  (20%) | 2  (50%) | 0 | 0 | 1  (25%) |  |
| Some college work | 6  (40%) | 1  (25%) | 1  (33.3%) | 3  (75%) | 1  (25%) |  |
| College graduate | 2  (13.3%) | 1  (25%) | 1  (33.3%) | 0 | 0 |  |
| Vocational training | 2  (13.3%) | 0 | 1  (33.3%) | 1  (25%) | 0 |  |
| **Ct exposure** | 10 |  |  |  |  |  |
| Positive antibody titer to Ct* | 9  (90%) | ND | 3  (100%) | 3  (75%) | 3  (75%) | NA |
| Patient reported past infection | 8  (80%) | ND | 2  (66%) | 3  (75%) | 3  (75%) | NA |
| **Insurance** |  |  |  |  |  | 0.14 |
| Private | 3  (20%) | 1  (25%) | 2  (66.7%) | 0 | 0 |  |
| Medicaid | 6  (40%) | 1  (25%) | 0 | 2  (50%) | 3  (75%) |  |
| Other insurance | 2  (13.3%) | 0 | 0 | 2  (50%) | 0 |  |
| None | 4  (26.7%) | 2  (50%) | 1  (33.3%) | 0 | 1  (25%) |  |
| Smoking | 6  (40%) | 2  (50%) | 0 | 2  (50%) | 2  (50%) | 0.57 |
| Marijuana use | 6  (40%) | 2  (50%) | 0 | 2  (50%) | 2  (50%) | 0.57 |
| **Birth control** |  |  |  |  |  |  |
| OCPs | 3  (20%) | 0 | 1  (33.3%) | 2  (50%) | 0 | 0.23 |
| DMPA | 0 | 0 | 0 | 0 | 0 | \ |
| Intrauterine device | 0 | 0 | 0 | 0 | 0 | \ |
| Condoms | 8  (53.3%) | 3  (75%) | 0 | 3  (75%) | 2  (50%) | 0.27 |
| Abstinence | 0 | 0 | 0 | 0 | 0 | \ |
| Coitus interruptus | 7  (46.7%) | 2  (50%) | 0 | 2  (50%) | 3  (75%) | 0.34 |
| **Bacterial vaginosis at enrollment** |  |  |  |  |  | 0.81 |
| Negative bacterial vaginosis (Nugent Score=0~3) | 8  (72.7%) | 1  (50%) | 1  (100%) | 3  (75%) | 3  (75%) |  |
| Intermediate bacterial vaginosis (Nugent Score=4~6) | 2  (18.2%) | 1  (50%) | 0 | 1  (25%) | 0 |  |
| Positive bacterial vaginosis (Nugent Score=7~10) | 1  (9.1%) | 0 | 0 | 0 | 1 |  |
| *: antibody titers to Ct performed by microimmunofluorescence testing (REF Russell paper); + titer: > 1:16. | | | | | | |
| Data are number (%) of subjects, unless otherwise indicated. | | | | | | |
| Abbreviations: GED, general education development; OCP, oral contraceptive pills; DMPA, depot medroxyprogesterone; ND, not done; NA, not applicable. | | | | | | |
