## Supplemental Table 1-10 for "miRNA/mRNA analysis of increased TGF-β pathways drive epithelial-mesenchymal transition and regulatory T cell differentiation": Supp Table 2.docx

| **Supp Table 2. Characteristics of the individuals from which samples were obtained for miRNA data analysis.** | | | | | | |
| --- | --- | --- | --- | --- | --- | --- |
| **Characteristics** | **Number (%)** | | | | | **P value** |
|  |  | **Endo+ (N=9)** | | **Endo- (N=14)** | |  |
|  | **Total** | **Symptom+** | **Symptom-** | **Cervix+** | **Uninfected** |  |
| **Number of women per group** | 23 | 4 | 5 | 5 | 9 |  |
| **Age, y, median (CI)** | 20  (19-22) | 20  (18-22) | 20  (18-23) | 19.5  (19-29) | 21.5  (19-23.5) | 0.87 |
| **Race/Ethnicity** |  |  |  |  |  | 0.10 |
| African American | 13 (56.5%) | 2  (50%) | 5  (100%) | 1  (25%) | 5  (50%) |  |
| White | 5  (21.7%) | 2  (50%) | 0 | 2  (50%) | 1  (10%) |  |
| Multiracial | 5  (21.7%) | 0 | 0 | 1  (25%) | 4  (40%) |  |
| **Marital status** |  |  |  |  |  | 0.18 |
| Single | 19 (82.6%) | 2  (50%) | 5  (100%) | 3  (75%) | 9  (90%) |  |
| Living with partner | 4  (17.4%) | 2  (50%) | 0 | 1  (25%) | 1  (10%) |  |
| **Education** |  |  |  |  |  | 0.41 |
| Less than high school graduate | 3  (13.0%) | 1  (25%) | 0 | 1  (25%) | 1  (10%) |  |
| High school graduate or GED degree | 8  (34.8%) | 1  (25%) | 3  (60%) | 0 | 4  (40%) |  |
| Some college work | 9  (39.1%) | 2  (50%) | 2  (40%) | 1  (25%) | 4  (40%) |  |
| College graduate | 1  (4.3%) | 0 | 0 | 0 | 1  (10%) |  |
| Vocational training | 2  (8.7%) | 0 | 0 | 2  (50%) | 0 |  |
| **Ct exposure** | 19 |  |  |  |  |  |
| Positive antibody titer to Ct* | 16  (89%) | ND | 4  (80%) | 4  (80%) | 8  (89%) |  |
| Patient reported past infection | 10  (53%) | ND | 2  (40%) | 2  (40%) | 6  (67%) |  |
| **Insurance** |  |  |  |  |  | 0.88 |
| Private | 5  (21.7%) | 1  (25%) | 1  (20%) | 0 | 3  (30%) |  |
| Medicaid | 13 (56.5%) | 3  (75%) | 3  (60%) | 2  (50%) | 5  (50%) |  |
| Other insurance | 1  (4.3%) | 0 | 0 | 1  (25%) | 0 |  |
| None | 4  (17.4%) | 0 | 1  (20%) | 1  (25%) | 2  (30%) |  |
| Smoking | 12 (52.2%) | 2  (50%) | 2  (40%) | 2  (50%) | 6  (60%) | 0.93 |
| Marijuana use | 10 (43.5%) | 1  (25%) | 4  (80%) | 1  (25%) | 4  (40%) | 0.33 |
| **Birth control** |  |  |  |  |  |  |
| OCPs | 1  (4.3%) | 0 | 0 | 0 | 1  (10%) | 1.00 |
| DMPA | 3  (13.0%) | 0 | 2  (40%) | 0 | 1  (10%) | 0.29 |
| Intrauterine device | 1  (4.3%) | 0 | 1  (20%) | 0 | 0 | 0.57 |
| Condoms | 10 (43.5%) | 1  (25%) | 3  (60%) | 1  (25%) | 5  (50%) | 0.66 |
| Abstinence | 3  (13.0%) | 0 | 2  (40%) | 0 | 1  (10%) | 0.29 |
| Coitus interruptus | 8  (34.8%) | 2  (50%) | 2  (40%) | 1  (25%) | 3  (30%) | 0.93 |
| **Bacterial vaginosis at enrollment** |  |  |  |  |  | 1.00 |
| Negative bacterial vaginosis (Nugent Score=0~3) | 10 (66.7%) | 2  (50%) | 2  (100%) | 1  (100%) | 5  (62.5%) |  |
| Intermediate bacterial vaginosis (Nugent Score=4~6) | 2  (13.3%) | 1  (25%) | 0 | 0 | 1  (12.5%) |  |
| Positive bacterial vaginosis (Nugent Score=7~10) | 3  (20%) | 1  (25%) | 0 | 0 | 2  (25%) |  |
| *: antibody titers to Ct performed by microimmunofluorescence testing (REF Russell paper); + titer: ≥ 1:16 | | | | | | |
| Data are number (%) of subjects, unless otherwise indicated. | | | | | | |
| Abbreviations: GED, general education development; OCP, oral contraceptive pills;  DMPA, depot medroxyprogesterone; ND, not done; NA, not applicable. | | | | | | |
