## Supplemental Table 1-10 for "miRNA/mRNA analysis of increased TGF-β pathways drive epithelial-mesenchymal transition and regulatory T cell differentiation": Supp Table 3.docx

| **Supp Table 3. Upregulated mRNAs in Endo+ compared to Endo- women with FDR<0.05 (Sorted by FDR values)** | | | |
| --- | --- | --- | --- |
| **Gene Symbol** | **Fold Change** | **P value** | **FDR** |
| IRAK3 | 4.2 | 9.26E-14 | 7.17E-10 |
| CXCL1 | 19.8 | 8.30E-14 | 7.17E-10 |
| SOD2 | 10.5 | 8.36E-12 | 2.59E-08 |
| C4BPA | 24.0 | 1.22E-11 | 3.15E-08 |
| MAP3K8 | 3.2 | 1.16E-10 | 2.25E-07 |
| BCL6 | 5.6 | 1.12E-10 | 2.25E-07 |
| FNBP1 | 1.9 | 4.08E-10 | 6.31E-07 |
| HP | 68.5 | 7.50E-10 | 7.91E-07 |
| SLC7A2 | 3.8 | 6.17E-10 | 7.91E-07 |
| SGPP2 | 3.0 | 8.21E-10 | 7.95E-07 |
| PTGS2 | 14.5 | 1.01E-09 | 9.24E-07 |
| GK | 8.7 | 1.25E-09 | 1.07E-06 |
| ARID5B | 1.9 | 2.31E-09 | 1.79E-06 |
| NFAT5 | 1.7 | 2.21E-09 | 1.79E-06 |
| G0S2 | 37.4 | 7.70E-09 | 5.28E-06 |
| CSGALNACT1 | 2.6 | 7.84E-09 | 5.28E-06 |
| SMAD3 | 1.7 | 8.87E-09 | 5.73E-06 |
| FLJ22447 | 10.6 | 1.13E-08 | 6.56E-06 |
| IL1RN | 20.5 | 1.19E-08 | 6.56E-06 |
| MCTP1 | 3.4 | 1.12E-08 | 6.56E-06 |
| ATP10B | 18.3 | 1.15E-08 | 6.56E-06 |
| CP | 9.1 | 1.77E-08 | 9.12E-06 |
| NCK1 | 1.6 | 1.92E-08 | 9.57E-06 |
| IL1R2 | 10.2 | 2.22E-08 | 1.02E-05 |
| FOXP1 | 2.0 | 2.58E-08 | 1.11E-05 |
| AAK1 | 1.5 | 4.08E-08 | 1.62E-05 |
| LOC646329 | 3.0 | 4.57E-08 | 1.77E-05 |
| FLJ43663 | 3.5 | 7.50E-08 | 2.83E-05 |
| ADTRP | 4.3 | 8.18E-08 | 2.95E-05 |
| TNIP3 | 14.1 | 9.82E-08 | 3.46E-05 |
| NFKBIZ | 4.2 | 1.09E-07 | 3.54E-05 |
| SLC16A10 | 11.5 | 1.10E-07 | 3.54E-05 |
| DEFB1 | 6.6 | 1.12E-07 | 3.54E-05 |
| NNMT | 9.5 | 1.18E-07 | 3.66E-05 |
| PRRG1 | 1.5 | 1.20E-07 | 3.66E-05 |
| TOX2 | 3.9 | 1.42E-07 | 4.11E-05 |
| ACER3 | 1.7 | 1.50E-07 | 4.14E-05 |
| DOC2B | 7.1 | 1.63E-07 | 4.43E-05 |
| C2CD4A | 30.0 | 2.48E-07 | 6.26E-05 |
| IL15 | 2.6 | 2.44E-07 | 6.26E-05 |
| BLK | 15.7 | 2.51E-07 | 6.26E-05 |
| WHSC1L1 | 1.2 | 2.66E-07 | 6.43E-05 |
| BCL2A1 | 21.2 | 3.12E-07 | 7.11E-05 |
| FCAR | 71.3 | 3.30E-07 | 7.20E-05 |
| NABP1 | 3.8 | 3.30E-07 | 7.20E-05 |
| TAOK3 | 1.7 | 3.49E-07 | 7.40E-05 |
| TNFAIP6 | 14.1 | 3.61E-07 | 7.46E-05 |
| INPP4B | 2.5 | 3.74E-07 | 7.62E-05 |
| PTPN12 | 1.5 | 3.92E-07 | 7.75E-05 |
| SLC6A20 | 8.4 | 4.93E-07 | 9.19E-05 |
| SNX10 | 6.1 | 4.99E-07 | 9.19E-05 |
| ZC3H12C | 1.5 | 5.18E-07 | 9.45E-05 |
| SPP1 | 9.6 | 5.29E-07 | 9.53E-05 |
| CFB | 14.5 | 6.22E-07 | 1.08E-04 |
| FCRL2 | 14.3 | 6.47E-07 | 1.09E-04 |
| IL1B | 19.5 | 6.87E-07 | 1.13E-04 |
| LITAF | 2.4 | 7.31E-07 | 1.19E-04 |
| SERPINB9 | 4.8 | 7.97E-07 | 1.27E-04 |
| POU2AF1 | 6.1 | 8.24E-07 | 1.29E-04 |
| FAM129A | 4.5 | 8.62E-07 | 1.33E-04 |
| AQP9 | 21.0 | 9.79E-07 | 1.49E-04 |
| R3HCC1L | 1.3 | 1.11E-06 | 1.62E-04 |
| LILRB2 | 11.5 | 1.10E-06 | 1.62E-04 |
| SLIT3 | 3.2 | 1.09E-06 | 1.62E-04 |
| IL4R | 3.1 | 1.18E-06 | 1.71E-04 |
| DNER | 6.0 | 1.21E-06 | 1.74E-04 |
| PIK3AP1 | 7.3 | 1.48E-06 | 2.01E-04 |
| FOXN3 | 1.7 | 1.48E-06 | 2.01E-04 |
| NCOA7 | 3.9 | 1.49E-06 | 2.01E-04 |
| GFOD1 | 4.3 | 1.54E-06 | 2.04E-04 |
| SLAMF1 | 9.3 | 1.57E-06 | 2.05E-04 |
| SLC4A10 | 5.3 | 1.56E-06 | 2.05E-04 |
| SLC19A3 | 14.4 | 1.67E-06 | 2.13E-04 |
| ST6GAL1 | 2.3 | 1.68E-06 | 2.13E-04 |
| RNF111 | 1.3 | 1.84E-06 | 2.28E-04 |
| KLF5 | 1.7 | 1.89E-06 | 2.31E-04 |
| NAMPT | 5.8 | 1.89E-06 | 2.31E-04 |
| TNFAIP3 | 4.7 | 2.00E-06 | 2.42E-04 |
| GSDMC | 21.6 | 2.42E-06 | 2.82E-04 |
| CCL2 | 7.6 | 2.61E-06 | 2.97E-04 |
| TMC5 | 6.4 | 2.69E-06 | 3.04E-04 |
| FLVCR2 | 2.8 | 2.76E-06 | 3.10E-04 |
| PM20D1 | 4.5 | 2.94E-06 | 3.25E-04 |
| LILRA6 | 12.7 | 3.09E-06 | 3.34E-04 |
| BTNL8 | 10.6 | 3.06E-06 | 3.34E-04 |
| MED13L | 1.6 | 3.17E-06 | 3.38E-04 |
| LYN | 3.5 | 3.34E-06 | 3.50E-04 |
| HIVEP3 | 2.2 | 3.52E-06 | 3.64E-04 |
| GALNT14 | 8.2 | 3.55E-06 | 3.64E-04 |
| NPAS2 | 2.7 | 3.54E-06 | 3.64E-04 |
| MAP3K5 | 3.2 | 4.04E-06 | 4.07E-04 |
| SRD5A3 | 3.2 | 4.08E-06 | 4.08E-04 |
| ALOX5AP | 4.2 | 4.17E-06 | 4.11E-04 |
| RHOH | 6.4 | 4.31E-06 | 4.20E-04 |
| PALM2 | 3.2 | 4.78E-06 | 4.57E-04 |
| MGAT4A | 2.3 | 4.82E-06 | 4.58E-04 |
| PRDM2 | 1.3 | 5.04E-06 | 4.72E-04 |
| INO80D | 1.3 | 5.05E-06 | 4.72E-04 |
| PDZK1IP1 | 10.2 | 5.16E-06 | 4.73E-04 |
| CRTC3 | 1.4 | 5.16E-06 | 4.73E-04 |
| CR1 | 4.8 | 5.22E-06 | 4.74E-04 |
| SLC38A1 | 2.0 | 5.26E-06 | 4.74E-04 |
| HAL | 6.3 | 5.34E-06 | 4.74E-04 |
| HCK | 6.7 | 5.33E-06 | 4.74E-04 |
| LILRB1 | 6.3 | 5.56E-06 | 4.88E-04 |
| OSMR | 2.8 | 5.58E-06 | 4.88E-04 |
| C10orf76 | 1.4 | 6.14E-06 | 5.19E-04 |
| LAMB3 | 4.4 | 6.29E-06 | 5.21E-04 |
| RDH10 | 3.2 | 6.27E-06 | 5.21E-04 |
| GNA15 | 4.2 | 6.86E-06 | 5.57E-04 |
| EMR1 | 23.1 | 6.83E-06 | 5.57E-04 |
| TTC39C | 1.9 | 7.00E-06 | 5.65E-04 |
| DAPP1 | 5.1 | 7.29E-06 | 5.83E-04 |
| RABGEF1 | 1.4 | 7.30E-06 | 5.83E-04 |
| USP43 | 2.0 | 7.38E-06 | 5.86E-04 |
| NT5C2 | 1.5 | 7.98E-06 | 6.24E-04 |
| LOC100130476 | 3.9 | 8.21E-06 | 6.39E-04 |
| PRKCH | 1.7 | 9.26E-06 | 7.14E-04 |
| AFF1 | 1.6 | 9.68E-06 | 7.38E-04 |
| MFSD4 | 3.9 | 9.80E-06 | 7.44E-04 |
| LOC284100 | 2.4 | 9.90E-06 | 7.48E-04 |
| MS4A1 | 9.8 | 1.06E-05 | 7.88E-04 |
| SAMSN1 | 5.4 | 1.10E-05 | 8.10E-04 |
| RBPMS | 1.9 | 1.12E-05 | 8.21E-04 |
| ANKRD33B | 3.3 | 1.14E-05 | 8.26E-04 |
| ZC3H12A | 3.9 | 1.15E-05 | 8.35E-04 |
| RAB39A | 3.2 | 1.16E-05 | 8.39E-04 |
| SLCO4A1 | 4.8 | 1.18E-05 | 8.40E-04 |
| OLR1 | 11.3 | 1.22E-05 | 8.51E-04 |
| ELMO1 | 3.0 | 1.21E-05 | 8.51E-04 |
| KDM4C | 1.3 | 1.22E-05 | 8.51E-04 |
| SRD5A3-AS1 | 3.6 | 1.23E-05 | 8.52E-04 |
| STS | 1.5 | 1.24E-05 | 8.53E-04 |
| IL18R1 | 2.3 | 1.26E-05 | 8.62E-04 |
| CYP2C18 | 6.1 | 1.30E-05 | 8.72E-04 |
| CAPN13 | 13.9 | 1.30E-05 | 8.72E-04 |
| RARRES1 | 6.1 | 1.29E-05 | 8.72E-04 |
| HIF1A | 2.0 | 1.32E-05 | 8.84E-04 |
| ERO1L | 2.0 | 1.34E-05 | 8.92E-04 |
| KANSL1L | 1.3 | 1.36E-05 | 8.95E-04 |
| CD48 | 4.9 | 1.37E-05 | 8.97E-04 |
| RGL1 | 1.5 | 1.38E-05 | 8.99E-04 |
| RASA2 | 1.7 | 1.41E-05 | 9.12E-04 |
| CSF3R | 14.0 | 1.45E-05 | 9.29E-04 |
| NUMB | 1.7 | 1.48E-05 | 9.38E-04 |
| FOSL2 | 2.1 | 1.48E-05 | 9.38E-04 |
| ALPK2 | 3.9 | 1.50E-05 | 9.41E-04 |
| TLR4 | 3.7 | 1.50E-05 | 9.41E-04 |
| PLAC8 | 3.8 | 1.62E-05 | 1.01E-03 |
| DOCK8 | 4.3 | 1.69E-05 | 1.05E-03 |
| SP140 | 5.9 | 1.73E-05 | 1.06E-03 |
| NLRP3 | 6.3 | 1.78E-05 | 1.08E-03 |
| WNK1 | 1.3 | 1.77E-05 | 1.08E-03 |
| SLC11A1 | 7.2 | 1.78E-05 | 1.08E-03 |
| CATSPERB | 6.3 | 1.80E-05 | 1.08E-03 |
| PRDM8 | 3.1 | 1.92E-05 | 1.14E-03 |
| PPP3CC | 1.5 | 1.93E-05 | 1.14E-03 |
| FLI1 | 2.1 | 1.98E-05 | 1.15E-03 |
| SOCS3 | 9.8 | 1.97E-05 | 1.15E-03 |
| ARNTL | 2.2 | 1.99E-05 | 1.15E-03 |
| MEGF11 | 3.5 | 2.00E-05 | 1.15E-03 |
| RFTN1 | 1.7 | 2.03E-05 | 1.15E-03 |
| SBSPON | 2.4 | 2.10E-05 | 1.18E-03 |
| MARCO | 8.6 | 2.13E-05 | 1.19E-03 |
| LCP2 | 5.3 | 2.13E-05 | 1.19E-03 |
| SWAP70 | 1.6 | 2.16E-05 | 1.20E-03 |
| CXCL13 | 8.1 | 2.22E-05 | 1.23E-03 |
| TMEM154 | 3.6 | 2.25E-05 | 1.24E-03 |
| TCF7L2 | 1.8 | 2.35E-05 | 1.27E-03 |
| ZCCHC6 | 1.5 | 2.34E-05 | 1.27E-03 |
| EMR3 | 10.7 | 2.41E-05 | 1.29E-03 |
| CLEC4E | 10.2 | 2.53E-05 | 1.34E-03 |
| CD58 | 2.1 | 2.59E-05 | 1.35E-03 |
| CCDC3 | 2.4 | 2.60E-05 | 1.35E-03 |
| ANXA4 | 2.6 | 2.58E-05 | 1.35E-03 |
| GRIA1 | 5.4 | 2.59E-05 | 1.35E-03 |
| TNFAIP8 | 3.0 | 2.64E-05 | 1.36E-03 |
| PSEN1 | 1.3 | 2.71E-05 | 1.38E-03 |
| CNGB1 | 4.5 | 2.71E-05 | 1.38E-03 |
| FYN | 1.8 | 2.71E-05 | 1.38E-03 |
| EYS | 1.5 | 2.73E-05 | 1.38E-03 |
| TBXAS1 | 3.0 | 2.74E-05 | 1.38E-03 |
| CCL3 | 10.3 | 2.80E-05 | 1.39E-03 |
| EPAS1 | 2.4 | 2.80E-05 | 1.39E-03 |
| DPP4 | 6.2 | 2.81E-05 | 1.39E-03 |
| CLEC5A | 4.6 | 2.81E-05 | 1.39E-03 |
| PAX5 | 6.5 | 2.80E-05 | 1.39E-03 |
| B4GALT5 | 1.8 | 2.88E-05 | 1.40E-03 |
| POU2F2 | 4.4 | 2.92E-05 | 1.42E-03 |
| ANKRD22 | 5.9 | 3.01E-05 | 1.45E-03 |
| CCRL2 | 5.0 | 3.12E-05 | 1.49E-03 |
| PTPN1 | 1.8 | 3.17E-05 | 1.51E-03 |
| EFCAB4B | 1.7 | 3.32E-05 | 1.56E-03 |
| RXFP1 | 3.1 | 3.31E-05 | 1.56E-03 |
| IL12RB2 | 3.9 | 3.34E-05 | 1.57E-03 |
| IRAK2 | 3.8 | 3.39E-05 | 1.57E-03 |
| FAM157A | 4.2 | 3.38E-05 | 1.57E-03 |
| ACSL1 | 3.0 | 3.36E-05 | 1.57E-03 |
| TC2N | 2.4 | 3.42E-05 | 1.57E-03 |
| RAPGEF6 | 1.5 | 3.50E-05 | 1.59E-03 |
| ACSL4 | 2.2 | 3.65E-05 | 1.65E-03 |
| RRAGD | 1.6 | 3.68E-05 | 1.65E-03 |
| DENND3 | 2.5 | 3.68E-05 | 1.65E-03 |
| AATF | 1.4 | 3.88E-05 | 1.72E-03 |
| FCGR2A | 6.4 | 4.10E-05 | 1.81E-03 |
| SH3TC2 | 4.2 | 4.11E-05 | 1.81E-03 |
| IL1R1 | 2.3 | 4.14E-05 | 1.82E-03 |
| ANK1 | 5.2 | 4.27E-05 | 1.86E-03 |
| LPXN | 1.9 | 4.32E-05 | 1.87E-03 |
| ANKRD44 | 2.2 | 4.46E-05 | 1.92E-03 |
| MUC4 | 5.0 | 4.50E-05 | 1.93E-03 |
| BIRC3 | 3.7 | 4.57E-05 | 1.94E-03 |
| LINC00341 | 2.5 | 4.62E-05 | 1.94E-03 |
| CCR1 | 5.2 | 4.59E-05 | 1.94E-03 |
| SKAP2 | 2.1 | 4.56E-05 | 1.94E-03 |
| NDRG1 | 2.7 | 4.62E-05 | 1.94E-03 |
| FAM157B | 4.1 | 4.61E-05 | 1.94E-03 |
| IL7 | 2.1 | 4.66E-05 | 1.94E-03 |
| IL4I1 | 5.6 | 4.69E-05 | 1.95E-03 |
| SYNE3 | 2.4 | 4.74E-05 | 1.96E-03 |
| THBD | 5.6 | 4.74E-05 | 1.96E-03 |
| SIGLEC5 | 6.0 | 4.76E-05 | 1.97E-03 |
| SYTL3 | 2.2 | 4.84E-05 | 1.98E-03 |
| FAM169B | 6.0 | 4.91E-05 | 1.99E-03 |
| CSF2RB | 6.4 | 5.01E-05 | 2.02E-03 |
| LOC100216545 | 1.4 | 5.02E-05 | 2.02E-03 |
| DRAM1 | 2.2 | 5.17E-05 | 2.07E-03 |
| PTPRC | 4.3 | 5.24E-05 | 2.09E-03 |
| GAB2 | 1.6 | 5.24E-05 | 2.09E-03 |
| CD55 | 3.7 | 5.33E-05 | 2.10E-03 |
| APBB1IP | 3.8 | 5.33E-05 | 2.10E-03 |
| TNF | 7.2 | 5.33E-05 | 2.10E-03 |
| STEAP4 | 4.2 | 5.31E-05 | 2.10E-03 |
| FAAH2 | 1.6 | 5.41E-05 | 2.11E-03 |
| IKZF5 | 1.4 | 5.44E-05 | 2.12E-03 |
| RORA | 2.1 | 5.47E-05 | 2.13E-03 |
| CTTNBP2NL | 1.6 | 5.51E-05 | 2.14E-03 |
| LILRB3 | 7.6 | 5.52E-05 | 2.14E-03 |
| RNF175 | 5.1 | 5.55E-05 | 2.14E-03 |
| WWC3 | 1.5 | 5.69E-05 | 2.19E-03 |
| C3AR1 | 4.2 | 5.86E-05 | 2.24E-03 |
| HCLS1 | 3.2 | 5.86E-05 | 2.24E-03 |
| ICAM1 | 5.3 | 5.91E-05 | 2.25E-03 |
| PSTPIP2 | 2.5 | 6.02E-05 | 2.28E-03 |
| FGR | 5.0 | 6.11E-05 | 2.31E-03 |
| FAM107B | 1.8 | 6.10E-05 | 2.31E-03 |
| CCL4 | 7.3 | 6.13E-05 | 2.31E-03 |
| LRRC8C | 1.9 | 6.20E-05 | 2.32E-03 |
| VNN3 | 22.3 | 6.67E-05 | 2.46E-03 |
| ITGAM | 2.8 | 6.98E-05 | 2.54E-03 |
| TLR2 | 3.9 | 6.96E-05 | 2.54E-03 |
| HDAC9 | 2.4 | 6.95E-05 | 2.54E-03 |
| SLC5A5 | 34.4 | 7.27E-05 | 2.63E-03 |
| TCN1 | 15.4 | 7.37E-05 | 2.64E-03 |
| CASS4 | 5.7 | 7.39E-05 | 2.64E-03 |
| AQP3 | 5.7 | 7.38E-05 | 2.64E-03 |
| AIM2 | 8.2 | 7.44E-05 | 2.65E-03 |
| TMEM165 | 1.6 | 7.45E-05 | 2.65E-03 |
| BCL3 | 3.0 | 7.53E-05 | 2.66E-03 |
| AOX1 | 7.1 | 7.76E-05 | 2.73E-03 |
| METAP2 | 1.4 | 8.05E-05 | 2.80E-03 |
| IL18RAP | 6.9 | 8.44E-05 | 2.90E-03 |
| ELOVL7 | 2.2 | 8.76E-05 | 2.99E-03 |
| ZHX2 | 1.6 | 9.20E-05 | 3.12E-03 |
| VAV1 | 3.9 | 9.29E-05 | 3.14E-03 |
| CSF1 | 2.2 | 9.37E-05 | 3.16E-03 |
| FCHSD2 | 1.7 | 9.48E-05 | 3.17E-03 |
| CYP24A1 | 11.9 | 9.48E-05 | 3.17E-03 |
| PIK3R5 | 3.8 | 9.67E-05 | 3.21E-03 |
| ABL2 | 1.5 | 9.84E-05 | 3.24E-03 |
| LOC284581 | 6.8 | 9.87E-05 | 3.24E-03 |
| CXCL14 | 20.8 | 1.03E-04 | 3.37E-03 |
| CTBS | 1.4 | 1.04E-04 | 3.41E-03 |
| NCF2 | 4.7 | 1.08E-04 | 3.50E-03 |
| TNFSF8 | 3.3 | 1.10E-04 | 3.51E-03 |
| SMURF1 | 1.5 | 1.13E-04 | 3.57E-03 |
| UBE2W | 1.5 | 1.13E-04 | 3.58E-03 |
| CFLAR | 1.6 | 1.15E-04 | 3.63E-03 |
| AKR1B15 | 8.9 | 1.16E-04 | 3.63E-03 |
| CDK13 | 1.2 | 1.18E-04 | 3.68E-03 |
| CD200R1 | 2.5 | 1.20E-04 | 3.74E-03 |
| TNFSF14 | 4.6 | 1.21E-04 | 3.75E-03 |
| CEBPD | 2.8 | 1.21E-04 | 3.75E-03 |
| ST3GAL1 | 2.4 | 1.21E-04 | 3.75E-03 |
| FAM89A | 2.1 | 1.21E-04 | 3.76E-03 |
| RIN2 | 2.2 | 1.27E-04 | 3.92E-03 |
| NFE2L2 | 1.6 | 1.30E-04 | 3.96E-03 |
| CPEB4 | 1.8 | 1.31E-04 | 3.96E-03 |
| SIK2 | 1.5 | 1.32E-04 | 3.98E-03 |
| STAT3 | 1.8 | 1.32E-04 | 3.98E-03 |
| SPI1 | 3.9 | 1.37E-04 | 4.08E-03 |
| IL21R | 4.7 | 1.41E-04 | 4.18E-03 |
| FGF11 | 1.9 | 1.48E-04 | 4.33E-03 |
| TTBK2 | 1.4 | 1.48E-04 | 4.33E-03 |
| ZBTB43 | 1.5 | 1.48E-04 | 4.33E-03 |
| PRKCB | 3.6 | 1.50E-04 | 4.37E-03 |
| LCN2 | 8.6 | 1.53E-04 | 4.44E-03 |
| CYBB | 2.8 | 1.53E-04 | 4.44E-03 |
| C10orf128 | 3.7 | 1.55E-04 | 4.48E-03 |
| LRMP | 2.5 | 1.56E-04 | 4.52E-03 |
| MAN1C1 | 2.0 | 1.57E-04 | 4.53E-03 |
| SLCO3A1 | 2.1 | 1.58E-04 | 4.54E-03 |
| THEMIS2 | 3.6 | 1.59E-04 | 4.57E-03 |
| PADI2 | 5.6 | 1.60E-04 | 4.58E-03 |
| KYNU | 3.9 | 1.65E-04 | 4.69E-03 |
| AP4S1 | 1.5 | 1.66E-04 | 4.70E-03 |
| CCBP2 | 2.9 | 1.67E-04 | 4.71E-03 |
| LOC100289230 | 1.6 | 1.74E-04 | 4.85E-03 |
| RASGRP4 | 4.2 | 1.75E-04 | 4.87E-03 |
| TEX15 | 10.2 | 1.75E-04 | 4.87E-03 |
| TGFBR2 | 1.8 | 1.78E-04 | 4.95E-03 |
| GNAI1 | 1.7 | 1.79E-04 | 4.95E-03 |
| GPR97 | 6.7 | 1.86E-04 | 5.12E-03 |
| SDS | 5.0 | 1.88E-04 | 5.14E-03 |
| FYB | 3.9 | 1.87E-04 | 5.14E-03 |
| CD226 | 2.3 | 1.95E-04 | 5.31E-03 |
| CPNE5 | 2.6 | 1.96E-04 | 5.34E-03 |
| MGAM | 7.3 | 1.99E-04 | 5.37E-03 |
| PTGER3 | 2.4 | 2.00E-04 | 5.38E-03 |
| PLEK | 3.6 | 2.00E-04 | 5.38E-03 |
| SERPINB1 | 2.5 | 2.00E-04 | 5.38E-03 |
| ITSN2 | 1.4 | 2.03E-04 | 5.43E-03 |
| RNF217 | 1.5 | 2.03E-04 | 5.43E-03 |
| PHC3 | 1.3 | 2.06E-04 | 5.48E-03 |
| FCN1 | 7.9 | 2.11E-04 | 5.58E-03 |
| KCNQ5 | 3.0 | 2.20E-04 | 5.79E-03 |
| RAPGEF5 | 1.7 | 2.22E-04 | 5.82E-03 |
| PLA2G4A | 2.2 | 2.26E-04 | 5.89E-03 |
| SLC26A8 | 2.7 | 2.26E-04 | 5.89E-03 |
| RAP1A | 1.7 | 2.29E-04 | 5.90E-03 |
| SNX27 | 1.3 | 2.28E-04 | 5.90E-03 |
| GPR65 | 3.8 | 2.32E-04 | 5.95E-03 |
| ALPK1 | 1.6 | 2.35E-04 | 6.03E-03 |
| HRH1 | 1.8 | 2.45E-04 | 6.24E-03 |
| CHI3L1 | 13.0 | 2.46E-04 | 6.25E-03 |
| CMIP | 1.6 | 2.50E-04 | 6.33E-03 |
| HABP2 | 8.8 | 2.51E-04 | 6.35E-03 |
| C14orf182 | 2.7 | 2.56E-04 | 6.45E-03 |
| FMNL1 | 3.4 | 2.56E-04 | 6.45E-03 |
| SIRPB1 | 5.4 | 2.59E-04 | 6.49E-03 |
| ELL2 | 3.1 | 2.59E-04 | 6.49E-03 |
| PTGES | 3.7 | 2.62E-04 | 6.56E-03 |
| ARHGAP25 | 2.8 | 2.63E-04 | 6.56E-03 |
| FILIP1 | 4.0 | 2.63E-04 | 6.56E-03 |
| IRF2 | 1.8 | 2.66E-04 | 6.60E-03 |
| SORCS1 | 4.5 | 2.69E-04 | 6.61E-03 |
| NFKB1 | 1.7 | 2.70E-04 | 6.63E-03 |
| MT2A | 3.2 | 2.75E-04 | 6.70E-03 |
| ASXL1 | 1.3 | 2.75E-04 | 6.70E-03 |
| LDLRAD3 | 1.7 | 2.77E-04 | 6.72E-03 |
| MXD1 | 3.1 | 2.78E-04 | 6.72E-03 |
| PLCL1 | 4.2 | 2.78E-04 | 6.72E-03 |
| KIF13A | 1.4 | 2.77E-04 | 6.72E-03 |
| FTX | 1.4 | 2.77E-04 | 6.72E-03 |
| FCGR2C | 4.6 | 2.79E-04 | 6.73E-03 |
| TFEC | 3.7 | 2.84E-04 | 6.82E-03 |
| TNFRSF10C | 6.4 | 2.85E-04 | 6.85E-03 |
| DCUN1D3 | 2.1 | 2.86E-04 | 6.86E-03 |
| PTAFR | 3.8 | 2.91E-04 | 6.92E-03 |
| SLC9A8 | 1.4 | 2.91E-04 | 6.92E-03 |
| MICU1 | 1.3 | 2.94E-04 | 6.95E-03 |
| LOC728558 | 2.0 | 2.94E-04 | 6.95E-03 |
| NCOA3 | 1.5 | 2.93E-04 | 6.95E-03 |
| ATG7 | 1.4 | 2.95E-04 | 6.95E-03 |
| IDO1 | 4.0 | 2.94E-04 | 6.95E-03 |
| DCP1A | 1.3 | 2.97E-04 | 6.96E-03 |
| GRB10 | 1.6 | 2.98E-04 | 6.98E-03 |
| DOCK4 | 2.1 | 3.08E-04 | 7.19E-03 |
| IL1A | 3.7 | 3.10E-04 | 7.21E-03 |
| ETV6 | 1.8 | 3.12E-04 | 7.22E-03 |
| SLC25A12 | 1.3 | 3.11E-04 | 7.22E-03 |
| EML4 | 1.5 | 3.14E-04 | 7.24E-03 |
| RASSF5 | 3.6 | 3.15E-04 | 7.24E-03 |
| IPMK | 1.6 | 3.14E-04 | 7.24E-03 |
| TSHZ2 | 1.9 | 3.18E-04 | 7.31E-03 |
| BATF | 10.7 | 3.20E-04 | 7.31E-03 |
| FAM20A | 3.1 | 3.21E-04 | 7.34E-03 |
| MAPK14 | 1.3 | 3.22E-04 | 7.34E-03 |
| IL2RA | 3.9 | 3.23E-04 | 7.35E-03 |
| STAT4 | 2.6 | 3.23E-04 | 7.35E-03 |
| RUFY3 | 1.4 | 3.24E-04 | 7.35E-03 |
| RNF19B | 1.8 | 3.29E-04 | 7.44E-03 |
| LIMK2 | 2.1 | 3.36E-04 | 7.57E-03 |
| KCNQ3 | 2.5 | 3.36E-04 | 7.57E-03 |
| DOCK10 | 2.5 | 3.39E-04 | 7.61E-03 |
| EMB | 3.1 | 3.39E-04 | 7.62E-03 |
| LAPTM5 | 3.2 | 3.41E-04 | 7.63E-03 |
| IFI16 | 2.2 | 3.46E-04 | 7.72E-03 |
| PCED1B-AS1 | 2.1 | 3.48E-04 | 7.72E-03 |
| PAK2 | 1.3 | 3.51E-04 | 7.77E-03 |
| KIAA0247 | 2.1 | 3.53E-04 | 7.81E-03 |
| CYP2C19 | 5.5 | 3.56E-04 | 7.83E-03 |
| CIR1 | 1.4 | 3.57E-04 | 7.84E-03 |
| CHIC2 | 1.5 | 3.59E-04 | 7.87E-03 |
| UPP1 | 2.2 | 3.63E-04 | 7.92E-03 |
| SRGN | 4.3 | 3.66E-04 | 7.96E-03 |
| RAP1B | 1.8 | 3.65E-04 | 7.96E-03 |
| IRF4 | 5.8 | 3.66E-04 | 7.96E-03 |
| NCMAP | 3.5 | 3.78E-04 | 8.17E-03 |
| CHST11 | 2.0 | 3.81E-04 | 8.22E-03 |
| SFMBT2 | 2.3 | 3.82E-04 | 8.23E-03 |
| LOC653513 | 1.8 | 3.86E-04 | 8.30E-03 |
| CLU | 3.0 | 3.89E-04 | 8.35E-03 |
| TYMP | 3.7 | 4.00E-04 | 8.53E-03 |
| C4orf19 | 1.9 | 4.01E-04 | 8.53E-03 |
| HMGN2P46 | 2.1 | 4.04E-04 | 8.57E-03 |
| CLEC7A | 4.3 | 4.08E-04 | 8.64E-03 |
| STX11 | 2.8 | 4.12E-04 | 8.69E-03 |
| KAT6A | 1.4 | 4.18E-04 | 8.80E-03 |
| C5AR1 | 3.2 | 4.28E-04 | 8.94E-03 |
| SLC9A7 | 1.6 | 4.29E-04 | 8.94E-03 |
| PROM1 | 3.3 | 4.31E-04 | 8.96E-03 |
| KLF3 | 1.5 | 4.38E-04 | 9.06E-03 |
| SP140L | 1.9 | 4.42E-04 | 9.11E-03 |
| TGFB2 | 3.2 | 4.46E-04 | 9.18E-03 |
| LCOR | 1.4 | 4.49E-04 | 9.23E-03 |
| EMR2 | 4.3 | 4.53E-04 | 9.25E-03 |
| OSCAR | 3.7 | 4.51E-04 | 9.25E-03 |
| ZBTB20 | 1.5 | 4.53E-04 | 9.25E-03 |
| VCAM1 | 5.0 | 4.55E-04 | 9.28E-03 |
| RAPH1 | 1.6 | 4.56E-04 | 9.28E-03 |
| IGSF21 | 3.8 | 4.62E-04 | 9.38E-03 |
| PTP4A2 | 1.4 | 4.67E-04 | 9.46E-03 |
| TRAF6 | 1.2 | 4.67E-04 | 9.46E-03 |
| LOC100271836 | 1.4 | 4.69E-04 | 9.46E-03 |
| AUH | 1.4 | 4.69E-04 | 9.46E-03 |
| FCER1G | 3.6 | 4.78E-04 | 9.57E-03 |
| FKBP5 | 3.6 | 4.78E-04 | 9.57E-03 |
| LOC283587 | 4.9 | 4.89E-04 | 9.73E-03 |
| SERPINB8 | 2.5 | 4.89E-04 | 9.73E-03 |
| CD86 | 3.4 | 4.89E-04 | 9.73E-03 |
| TNFRSF8 | 4.1 | 4.96E-04 | 9.75E-03 |
| REL | 1.9 | 4.95E-04 | 9.75E-03 |
| ICOS | 5.3 | 4.95E-04 | 9.75E-03 |
| SLC25A37 | 2.0 | 4.93E-04 | 9.75E-03 |
| C1S | 2.3 | 5.00E-04 | 9.79E-03 |
| FAM65B | 2.9 | 5.08E-04 | 9.92E-03 |
| TPK1 | 2.2 | 5.18E-04 | 1.01E-02 |
| FGF9 | 2.7 | 5.21E-04 | 1.01E-02 |
| CLEC12A | 4.6 | 5.23E-04 | 1.01E-02 |
| PAEP | 33.5 | 5.23E-04 | 1.01E-02 |
| MKLN1 | 1.4 | 5.27E-04 | 1.02E-02 |
| SLC2A14 | 3.2 | 5.32E-04 | 1.03E-02 |
| STK24 | 1.4 | 5.33E-04 | 1.03E-02 |
| ATXN7 | 1.4 | 5.34E-04 | 1.03E-02 |
| SPTBN1 | 1.6 | 5.38E-04 | 1.03E-02 |
| PLEKHM3 | 1.4 | 5.38E-04 | 1.03E-02 |
| CYB5R4 | 1.6 | 5.42E-04 | 1.03E-02 |
| CD3G | 3.7 | 5.48E-04 | 1.04E-02 |
| GPX3 | 7.8 | 5.48E-04 | 1.04E-02 |
| GDA | 3.3 | 5.50E-04 | 1.04E-02 |
| SMG7 | 1.2 | 5.53E-04 | 1.05E-02 |
| CREBRF | 1.7 | 5.57E-04 | 1.05E-02 |
| MBD2 | 1.4 | 5.63E-04 | 1.06E-02 |
| RALGAPA2 | 1.4 | 5.68E-04 | 1.07E-02 |
| SOS2 | 1.3 | 5.79E-04 | 1.08E-02 |
| TNFAIP2 | 2.8 | 5.77E-04 | 1.08E-02 |
| CCR7 | 3.8 | 5.79E-04 | 1.08E-02 |
| SIRPA | 2.4 | 5.78E-04 | 1.08E-02 |
| BCMO1 | 2.5 | 5.92E-04 | 1.10E-02 |
| WIPF1 | 2.1 | 6.00E-04 | 1.11E-02 |
| CD28 | 3.5 | 6.04E-04 | 1.11E-02 |
| C2 | 2.3 | 6.07E-04 | 1.11E-02 |
| COL19A1 | 3.5 | 6.14E-04 | 1.12E-02 |
| RTN1 | 1.6 | 6.31E-04 | 1.14E-02 |
| ZBP1 | 4.0 | 6.30E-04 | 1.14E-02 |
| C9orf84 | 1.5 | 6.31E-04 | 1.14E-02 |
| ARHGAP15 | 3.2 | 6.47E-04 | 1.16E-02 |
| CLIP1 | 1.5 | 6.50E-04 | 1.16E-02 |
| ELL | 1.8 | 6.52E-04 | 1.17E-02 |
| S100A9 | 10.9 | 6.54E-04 | 1.17E-02 |
| TNFRSF10A | 1.7 | 6.77E-04 | 1.20E-02 |
| PHF11 | 1.7 | 6.83E-04 | 1.20E-02 |
| RBBP8 | 1.8 | 6.86E-04 | 1.21E-02 |
| RNF149 | 1.9 | 6.94E-04 | 1.22E-02 |
| SLC9A9 | 1.7 | 6.98E-04 | 1.22E-02 |
| MAP1LC3B2 | 1.5 | 7.08E-04 | 1.23E-02 |
| RNF19A | 1.8 | 7.09E-04 | 1.23E-02 |
| ASXL2 | 1.3 | 7.16E-04 | 1.24E-02 |
| LCP1 | 2.7 | 7.19E-04 | 1.24E-02 |
| CCDC69 | 2.0 | 7.23E-04 | 1.25E-02 |
| TRAF1 | 2.5 | 7.24E-04 | 1.25E-02 |
| NCOA2 | 1.4 | 7.34E-04 | 1.26E-02 |
| MPP7 | 1.5 | 7.36E-04 | 1.27E-02 |
| MGEA5 | 1.3 | 7.40E-04 | 1.27E-02 |
| CYSLTR1 | 2.6 | 7.50E-04 | 1.28E-02 |
| CYTIP | 4.0 | 7.54E-04 | 1.29E-02 |
| GJB7 | 2.1 | 7.65E-04 | 1.30E-02 |
| SMPDL3A | 2.4 | 7.64E-04 | 1.30E-02 |
| MTMR10 | 1.5 | 7.70E-04 | 1.30E-02 |
| HERPUD2 | 1.3 | 7.82E-04 | 1.31E-02 |
| MLL5 | 1.4 | 7.94E-04 | 1.32E-02 |
| GNAQ | 1.5 | 7.94E-04 | 1.32E-02 |
| SIGLEC7 | 3.2 | 7.96E-04 | 1.33E-02 |
| SERPING1 | 3.1 | 8.02E-04 | 1.33E-02 |
| EPB41L3 | 2.0 | 8.02E-04 | 1.33E-02 |
| SOS1 | 1.2 | 8.02E-04 | 1.33E-02 |
| LTF | 6.9 | 8.00E-04 | 1.33E-02 |
| ABLIM3 | 3.5 | 8.02E-04 | 1.33E-02 |
| GIGYF2 | 1.2 | 8.17E-04 | 1.35E-02 |
| ARHGAP42 | 1.7 | 8.23E-04 | 1.36E-02 |
| LINC00472 | 1.6 | 8.26E-04 | 1.36E-02 |
| PYROXD1 | 1.4 | 8.35E-04 | 1.36E-02 |
| TLR10 | 4.0 | 8.34E-04 | 1.36E-02 |
| WDFY3 | 1.2 | 8.33E-04 | 1.36E-02 |
| ST8SIA4 | 2.6 | 8.35E-04 | 1.36E-02 |
| PAPPA | 1.9 | 8.35E-04 | 1.36E-02 |
| ANKRD12 | 1.4 | 8.45E-04 | 1.37E-02 |
| B3GNT2 | 1.7 | 8.47E-04 | 1.37E-02 |
| LEPR | 1.8 | 8.49E-04 | 1.38E-02 |
| DOCK2 | 3.0 | 8.50E-04 | 1.38E-02 |
| FAM49B | 1.9 | 8.52E-04 | 1.38E-02 |
| ADORA2A-AS1 | 2.7 | 8.55E-04 | 1.38E-02 |
| TP63 | 2.6 | 8.59E-04 | 1.38E-02 |
| GRAMD1B | 3.0 | 8.70E-04 | 1.39E-02 |
| HERC4 | 1.3 | 8.72E-04 | 1.39E-02 |
| HK3 | 4.1 | 8.92E-04 | 1.42E-02 |
| CD163L1 | 2.3 | 8.95E-04 | 1.42E-02 |
| PPP2R5E | 1.4 | 9.03E-04 | 1.43E-02 |
| RAB27A | 1.9 | 9.11E-04 | 1.44E-02 |
| ADAM19 | 2.8 | 9.13E-04 | 1.44E-02 |
| NLRP1 | 2.1 | 9.17E-04 | 1.44E-02 |
| BANK1 | 2.1 | 9.32E-04 | 1.46E-02 |
| LACTB | 1.7 | 9.39E-04 | 1.47E-02 |
| NACC2 | 1.5 | 9.41E-04 | 1.47E-02 |
| IL1RAP | 2.0 | 9.46E-04 | 1.47E-02 |
| CASP1 | 3.2 | 9.56E-04 | 1.48E-02 |
| ALOX5 | 3.7 | 9.60E-04 | 1.49E-02 |
| SPATA9 | 2.0 | 9.62E-04 | 1.49E-02 |
| GNE | 1.5 | 9.62E-04 | 1.49E-02 |
| SLC7A5 | 3.2 | 9.69E-04 | 1.50E-02 |
| TMEM45B | 3.1 | 9.75E-04 | 1.50E-02 |
| APOL3 | 2.8 | 9.87E-04 | 1.52E-02 |
| OSTF1 | 1.4 | 9.88E-04 | 1.52E-02 |
| ARL11 | 3.3 | 9.92E-04 | 1.52E-02 |
| MYOZ2 | 2.2 | 9.93E-04 | 1.52E-02 |
| SLC35F1 | 1.8 | 9.91E-04 | 1.52E-02 |
| DSE | 2.0 | 9.94E-04 | 1.52E-02 |
| SH2D2A | 3.3 | 1.00E-03 | 1.53E-02 |
| ZNF169 | 1.4 | 1.01E-03 | 1.54E-02 |
| UBR2 | 1.4 | 1.01E-03 | 1.54E-02 |
| LHFPL3 | 3.1 | 1.01E-03 | 1.54E-02 |
| CIITA | 2.7 | 1.02E-03 | 1.54E-02 |
| OPTN | 1.7 | 1.04E-03 | 1.57E-02 |
| CD53 | 3.2 | 1.05E-03 | 1.59E-02 |
| MARK3 | 1.3 | 1.06E-03 | 1.59E-02 |
| N4BP1 | 1.6 | 1.06E-03 | 1.60E-02 |
| PTGER2 | 2.8 | 1.07E-03 | 1.61E-02 |
| PPP2R1B | 1.3 | 1.08E-03 | 1.62E-02 |
| NCF1 | 6.0 | 1.09E-03 | 1.62E-02 |
| APOBR | 4.8 | 1.09E-03 | 1.62E-02 |
| GAB3 | 2.0 | 1.11E-03 | 1.64E-02 |
| LAX1 | 3.7 | 1.12E-03 | 1.65E-02 |
| SFT2D2 | 1.4 | 1.12E-03 | 1.66E-02 |
| GPRIN3 | 1.7 | 1.13E-03 | 1.66E-02 |
| AKAP10 | 1.2 | 1.14E-03 | 1.68E-02 |
| USP32 | 1.4 | 1.14E-03 | 1.68E-02 |
| TNRC6B | 1.3 | 1.14E-03 | 1.68E-02 |
| OXNAD1 | 1.4 | 1.14E-03 | 1.68E-02 |
| ITGAX | 3.3 | 1.15E-03 | 1.68E-02 |
| F5 | 4.3 | 1.16E-03 | 1.68E-02 |
| SIGLEC10 | 3.9 | 1.16E-03 | 1.68E-02 |
| CD40 | 2.9 | 1.16E-03 | 1.68E-02 |
| CXorf21 | 5.4 | 1.16E-03 | 1.69E-02 |
| EPHA4 | 2.3 | 1.16E-03 | 1.69E-02 |
| SMURF2 | 1.4 | 1.17E-03 | 1.69E-02 |
| CDK17 | 1.7 | 1.17E-03 | 1.69E-02 |
| ICAM2 | 1.8 | 1.18E-03 | 1.70E-02 |
| NPL | 1.9 | 1.19E-03 | 1.71E-02 |
| ARHGAP29 | 1.6 | 1.20E-03 | 1.72E-02 |
| SESTD1 | 1.6 | 1.20E-03 | 1.72E-02 |
| CCDC109B | 2.0 | 1.20E-03 | 1.72E-02 |
| WDFY4 | 3.3 | 1.21E-03 | 1.72E-02 |
| TNFRSF1A | 1.5 | 1.21E-03 | 1.72E-02 |
| ELK3 | 1.8 | 1.22E-03 | 1.72E-02 |
| MZB1 | 4.1 | 1.21E-03 | 1.72E-02 |
| CCDC129 | 3.8 | 1.21E-03 | 1.72E-02 |
| GAPT | 3.2 | 1.22E-03 | 1.72E-02 |
| ABI1 | 1.4 | 1.22E-03 | 1.73E-02 |
| TLK2 | 1.3 | 1.22E-03 | 1.73E-02 |
| LEPREL1 | 2.1 | 1.23E-03 | 1.73E-02 |
| PHF21A | 1.5 | 1.23E-03 | 1.73E-02 |
| FLT3 | 3.4 | 1.23E-03 | 1.73E-02 |
| MYO1F | 2.8 | 1.23E-03 | 1.73E-02 |
| LDHAL6A | 1.7 | 1.24E-03 | 1.74E-02 |
| SLFN12L | 3.2 | 1.25E-03 | 1.75E-02 |
| PARN | 1.2 | 1.25E-03 | 1.75E-02 |
| ARHGAP30 | 2.9 | 1.25E-03 | 1.75E-02 |
| IL16 | 2.0 | 1.25E-03 | 1.75E-02 |
| MKL1 | 1.6 | 1.26E-03 | 1.75E-02 |
| KREMEN1 | 1.4 | 1.27E-03 | 1.77E-02 |
| MTMR3 | 1.3 | 1.28E-03 | 1.77E-02 |
| PTPN22 | 2.7 | 1.29E-03 | 1.77E-02 |
| PLEKHO2 | 1.8 | 1.28E-03 | 1.77E-02 |
| FPR3 | 2.7 | 1.28E-03 | 1.77E-02 |
| XYLT1 | 1.8 | 1.29E-03 | 1.78E-02 |
| GPR176 | 2.7 | 1.30E-03 | 1.78E-02 |
| CCR5 | 3.0 | 1.33E-03 | 1.82E-02 |
| XDH | 4.1 | 1.33E-03 | 1.82E-02 |
| IL10RA | 2.9 | 1.33E-03 | 1.82E-02 |
| SBF2 | 1.5 | 1.34E-03 | 1.83E-02 |
| RGS18 | 3.4 | 1.35E-03 | 1.83E-02 |
| HK1 | 1.3 | 1.36E-03 | 1.85E-02 |
| SLC6A12 | 3.6 | 1.38E-03 | 1.87E-02 |
| SMAP2 | 1.8 | 1.39E-03 | 1.87E-02 |
| ZMYM5 | 1.2 | 1.38E-03 | 1.87E-02 |
| MYO5A | 1.6 | 1.38E-03 | 1.87E-02 |
| C3 | 5.2 | 1.39E-03 | 1.87E-02 |
| ZFP36 | 4.2 | 1.39E-03 | 1.87E-02 |
| MAP4K4 | 1.4 | 1.39E-03 | 1.87E-02 |
| NFE2L3 | 1.5 | 1.39E-03 | 1.87E-02 |
| SRGAP1 | 1.6 | 1.40E-03 | 1.87E-02 |
| ELK4 | 1.5 | 1.41E-03 | 1.89E-02 |
| KLHL6 | 2.5 | 1.41E-03 | 1.89E-02 |
| PPARGC1A | 3.3 | 1.41E-03 | 1.89E-02 |
| CAB39 | 1.5 | 1.43E-03 | 1.91E-02 |
| PCDH17 | 2.6 | 1.44E-03 | 1.91E-02 |
| ARHGAP24 | 2.4 | 1.44E-03 | 1.92E-02 |
| DENND4A | 1.6 | 1.45E-03 | 1.93E-02 |
| CD37 | 2.6 | 1.46E-03 | 1.94E-02 |
| SMG1P1 | 1.4 | 1.47E-03 | 1.94E-02 |
| LAMC1 | 1.6 | 1.47E-03 | 1.95E-02 |
| EMBP1 | 2.2 | 1.48E-03 | 1.95E-02 |
| IL2RG | 3.6 | 1.48E-03 | 1.95E-02 |
| SSH2 | 1.6 | 1.48E-03 | 1.95E-02 |
| TNS4 | 3.2 | 1.49E-03 | 1.95E-02 |
| POU6F1 | 1.5 | 1.49E-03 | 1.96E-02 |
| UVRAG | 1.5 | 1.50E-03 | 1.96E-02 |
| IL10RB | 1.5 | 1.50E-03 | 1.96E-02 |
| ATP11A | 1.5 | 1.51E-03 | 1.97E-02 |
| TNFRSF9 | 2.5 | 1.51E-03 | 1.97E-02 |
| ZNF438 | 1.5 | 1.53E-03 | 1.98E-02 |
| SLAMF7 | 4.0 | 1.53E-03 | 1.98E-02 |
| STRN3 | 1.5 | 1.53E-03 | 1.98E-02 |
| ARFGEF1 | 1.3 | 1.53E-03 | 1.98E-02 |
| MAML2 | 1.6 | 1.54E-03 | 2.00E-02 |
| RGS5 | 1.5 | 1.55E-03 | 2.00E-02 |
| ZNF267 | 2.0 | 1.55E-03 | 2.00E-02 |
| MYO1G | 3.9 | 1.55E-03 | 2.00E-02 |
| PILRA | 2.5 | 1.55E-03 | 2.00E-02 |
| PPP2R3C | 1.3 | 1.56E-03 | 2.01E-02 |
| PTK2B | 2.4 | 1.56E-03 | 2.01E-02 |
| COL4A4 | 1.9 | 1.58E-03 | 2.03E-02 |
| TGFA | 1.8 | 1.59E-03 | 2.04E-02 |
| ANKRD36BP2 | 2.0 | 1.59E-03 | 2.04E-02 |
| SIGLEC9 | 3.2 | 1.60E-03 | 2.05E-02 |
| VEZT | 1.3 | 1.61E-03 | 2.05E-02 |
| MLKL | 3.2 | 1.61E-03 | 2.05E-02 |
| RBM33 | 1.2 | 1.62E-03 | 2.06E-02 |
| LINC00674 | 1.3 | 1.63E-03 | 2.07E-02 |
| NEAT1 | 2.1 | 1.63E-03 | 2.07E-02 |
| ARHGAP31 | 1.5 | 1.63E-03 | 2.07E-02 |
| PTPRJ | 1.8 | 1.64E-03 | 2.07E-02 |
| RIMKLB | 3.0 | 1.64E-03 | 2.07E-02 |
| ARID4B | 1.4 | 1.64E-03 | 2.07E-02 |
| MSN | 1.5 | 1.65E-03 | 2.08E-02 |
| FAM219A | 1.4 | 1.65E-03 | 2.08E-02 |
| VNN1 | 4.3 | 1.66E-03 | 2.09E-02 |
| CSGALNACT2 | 1.5 | 1.68E-03 | 2.10E-02 |
| ARRB1 | 1.6 | 1.68E-03 | 2.11E-02 |
| C3orf55 | 3.7 | 1.68E-03 | 2.11E-02 |
| CLOCK | 1.2 | 1.68E-03 | 2.11E-02 |
| TMEM196 | 6.2 | 1.68E-03 | 2.11E-02 |
| MCU | 1.5 | 1.70E-03 | 2.13E-02 |
| KCNK1 | 2.2 | 1.71E-03 | 2.13E-02 |
| LPIN2 | 1.5 | 1.71E-03 | 2.13E-02 |
| PYHIN1 | 2.8 | 1.71E-03 | 2.13E-02 |
| SPDYA | 1.5 | 1.71E-03 | 2.13E-02 |
| HIF1A-AS2 | 2.6 | 1.72E-03 | 2.14E-02 |
| MCTP2 | 1.9 | 1.72E-03 | 2.14E-02 |
| CMKLR1 | 2.2 | 1.73E-03 | 2.14E-02 |
| SLC2A5 | 3.0 | 1.74E-03 | 2.16E-02 |
| SELL | 9.2 | 1.75E-03 | 2.16E-02 |
| ACVR1C | 1.9 | 1.76E-03 | 2.17E-02 |
| RASGRP3 | 1.8 | 1.79E-03 | 2.21E-02 |
| CREBL2 | 1.5 | 1.80E-03 | 2.22E-02 |
| POU2F1 | 1.3 | 1.82E-03 | 2.23E-02 |
| DGAT2 | 2.9 | 1.83E-03 | 2.24E-02 |
| MFHAS1 | 1.6 | 1.83E-03 | 2.24E-02 |
| GPR110 | 11.4 | 1.83E-03 | 2.24E-02 |
| CYP2C9 | 3.6 | 1.84E-03 | 2.24E-02 |
| PTPN2 | 1.4 | 1.84E-03 | 2.24E-02 |
| DESI1 | 1.4 | 1.84E-03 | 2.24E-02 |
| KIAA0430 | 1.3 | 1.85E-03 | 2.25E-02 |
| CD163 | 3.0 | 1.86E-03 | 2.25E-02 |
| LRRK1 | 1.4 | 1.88E-03 | 2.28E-02 |
| ZNF831 | 3.1 | 1.88E-03 | 2.28E-02 |
| LYST | 1.6 | 1.88E-03 | 2.28E-02 |
| NCF4 | 3.3 | 1.89E-03 | 2.28E-02 |
| LOC100506776 | 3.1 | 1.92E-03 | 2.32E-02 |
| CD84 | 2.7 | 1.93E-03 | 2.32E-02 |
| ERO1LB | 1.6 | 1.94E-03 | 2.32E-02 |
| BTK | 3.2 | 1.95E-03 | 2.34E-02 |
| C4orf32 | 1.9 | 1.96E-03 | 2.35E-02 |
| TRAF3 | 1.6 | 1.96E-03 | 2.35E-02 |
| CR1L | 2.4 | 1.98E-03 | 2.36E-02 |
| P2RY6 | 3.9 | 1.98E-03 | 2.36E-02 |
| SLC34A2 | 2.3 | 1.98E-03 | 2.36E-02 |
| TBCK | 1.2 | 1.98E-03 | 2.36E-02 |
| DOK3 | 3.2 | 1.99E-03 | 2.37E-02 |
| ASH1L | 1.2 | 1.99E-03 | 2.37E-02 |
| SH3KBP1 | 1.6 | 1.99E-03 | 2.37E-02 |
| CYTH4 | 3.4 | 2.00E-03 | 2.38E-02 |
| CARD8 | 1.4 | 2.01E-03 | 2.38E-02 |
| RAB21 | 1.5 | 2.02E-03 | 2.39E-02 |
| PLSCR1 | 2.2 | 2.03E-03 | 2.40E-02 |
| TLL2 | 2.5 | 2.04E-03 | 2.40E-02 |
| STAT5B | 1.5 | 2.03E-03 | 2.40E-02 |
| TLR8 | 5.2 | 2.03E-03 | 2.40E-02 |
| CMAHP | 2.4 | 2.04E-03 | 2.40E-02 |
| SCML4 | 3.6 | 2.04E-03 | 2.40E-02 |
| NFKBIA | 1.8 | 2.05E-03 | 2.41E-02 |
| ADAMTS1 | 2.6 | 2.08E-03 | 2.44E-02 |
| RC3H2 | 1.2 | 2.08E-03 | 2.44E-02 |
| RAB11FIP1 | 1.6 | 2.12E-03 | 2.46E-02 |
| RASGRP2 | 2.7 | 2.13E-03 | 2.47E-02 |
| SLC15A1 | 8.4 | 2.14E-03 | 2.48E-02 |
| INPP4A | 1.4 | 2.17E-03 | 2.52E-02 |
| UGCG | 2.1 | 2.18E-03 | 2.52E-02 |
| SNTB1 | 1.6 | 2.19E-03 | 2.52E-02 |
| GBP2 | 2.9 | 2.20E-03 | 2.53E-02 |
| PTPRE | 2.7 | 2.20E-03 | 2.53E-02 |
| CLECL1 | 3.5 | 2.20E-03 | 2.53E-02 |
| MED13 | 1.3 | 2.19E-03 | 2.53E-02 |
| EIF4E3 | 1.8 | 2.20E-03 | 2.53E-02 |
| HIVEP1 | 1.7 | 2.20E-03 | 2.53E-02 |
| HPSE | 2.9 | 2.21E-03 | 2.54E-02 |
| KIAA0513 | 1.8 | 2.23E-03 | 2.55E-02 |
| RUNX3 | 3.9 | 2.23E-03 | 2.55E-02 |
| ARSB | 1.8 | 2.24E-03 | 2.56E-02 |
| TRERF1 | 1.5 | 2.25E-03 | 2.56E-02 |
| FSD1L | 1.7 | 2.28E-03 | 2.58E-02 |
| GCH1 | 2.0 | 2.28E-03 | 2.58E-02 |
| BCAS3 | 1.5 | 2.32E-03 | 2.61E-02 |
| FAM126B | 1.3 | 2.32E-03 | 2.61E-02 |
| RUNX1 | 2.5 | 2.31E-03 | 2.61E-02 |
| STAU2 | 1.3 | 2.32E-03 | 2.61E-02 |
| CTSB | 2.3 | 2.33E-03 | 2.62E-02 |
| PIK3CB | 1.4 | 2.34E-03 | 2.63E-02 |
| S100A1 | 4.6 | 2.35E-03 | 2.63E-02 |
| LILRB4 | 3.3 | 2.36E-03 | 2.64E-02 |
| STK17B | 2.2 | 2.36E-03 | 2.65E-02 |
| TFPI | 2.1 | 2.37E-03 | 2.65E-02 |
| CHD2 | 1.2 | 2.38E-03 | 2.66E-02 |
| AASS | 1.7 | 2.39E-03 | 2.67E-02 |
| C4orf6 | 2.3 | 2.40E-03 | 2.67E-02 |
| FAM49A | 2.4 | 2.41E-03 | 2.68E-02 |
| TNRC6C | 1.4 | 2.42E-03 | 2.68E-02 |
| PRLR | 2.1 | 2.42E-03 | 2.68E-02 |
| SLCO4C1 | 3.6 | 2.42E-03 | 2.68E-02 |
| CCDC126 | 1.4 | 2.43E-03 | 2.68E-02 |
| TMEM71 | 2.8 | 2.43E-03 | 2.68E-02 |
| FCHO2 | 1.4 | 2.45E-03 | 2.70E-02 |
| ZEB2 | 1.6 | 2.47E-03 | 2.72E-02 |
| SPPL2A | 1.3 | 2.50E-03 | 2.74E-02 |
| ADAM23 | 2.1 | 2.51E-03 | 2.75E-02 |
| ENOX2 | 1.4 | 2.52E-03 | 2.75E-02 |
| PIAS1 | 1.4 | 2.52E-03 | 2.75E-02 |
| SBNO2 | 2.3 | 2.53E-03 | 2.76E-02 |
| PARP15 | 2.6 | 2.52E-03 | 2.76E-02 |
| LINC00152 | 2.7 | 2.54E-03 | 2.77E-02 |
| LOC100130231 | 2.8 | 2.59E-03 | 2.80E-02 |
| DPYD | 1.9 | 2.60E-03 | 2.81E-02 |
| CD6 | 3.1 | 2.60E-03 | 2.81E-02 |
| NUAK1 | 1.6 | 2.60E-03 | 2.81E-02 |
| TMOD1 | 2.4 | 2.61E-03 | 2.82E-02 |
| CPVL | 2.1 | 2.62E-03 | 2.83E-02 |
| MT1G | 5.9 | 2.63E-03 | 2.83E-02 |
| ATP6V1C1 | 1.3 | 2.63E-03 | 2.83E-02 |
| IFNAR1 | 1.4 | 2.65E-03 | 2.84E-02 |
| NEMF | 1.2 | 2.65E-03 | 2.84E-02 |
| TNFRSF1B | 2.1 | 2.66E-03 | 2.85E-02 |
| KIAA0922 | 1.6 | 2.66E-03 | 2.85E-02 |
| FAM105B | 1.2 | 2.67E-03 | 2.85E-02 |
| ZNF407 | 1.3 | 2.69E-03 | 2.86E-02 |
| SLC7A7 | 2.3 | 2.70E-03 | 2.87E-02 |
| CCRN4L | 1.7 | 2.71E-03 | 2.88E-02 |
| LTB | 3.5 | 2.71E-03 | 2.88E-02 |
| CFH | 2.5 | 2.73E-03 | 2.89E-02 |
| GPR114 | 2.7 | 2.74E-03 | 2.90E-02 |
| LRRFIP1 | 1.7 | 2.75E-03 | 2.91E-02 |
| SLAMF6 | 3.2 | 2.78E-03 | 2.94E-02 |
| PCNX | 1.5 | 2.79E-03 | 2.95E-02 |
| C11orf75 | 1.6 | 2.80E-03 | 2.96E-02 |
| ITK | 2.8 | 2.82E-03 | 2.97E-02 |
| BATF2 | 4.0 | 2.82E-03 | 2.97E-02 |
| EGLN1 | 1.4 | 2.83E-03 | 2.97E-02 |
| JHDM1D | 1.5 | 2.84E-03 | 2.98E-02 |
| GABRE | 2.7 | 2.84E-03 | 2.98E-02 |
| USP15 | 1.7 | 2.86E-03 | 2.99E-02 |
| SELP | 3.5 | 2.88E-03 | 3.00E-02 |
| EPB41L4A | 1.5 | 2.89E-03 | 3.01E-02 |
| KIF2A | 1.3 | 2.90E-03 | 3.01E-02 |
| MST4 | 1.6 | 2.90E-03 | 3.01E-02 |
| DAPK1 | 1.9 | 2.91E-03 | 3.01E-02 |
| NR4A3 | 4.1 | 2.90E-03 | 3.01E-02 |
| SPPL3 | 1.4 | 2.92E-03 | 3.02E-02 |
| MYO16 | 1.9 | 2.92E-03 | 3.02E-02 |
| CNTD1 | 1.6 | 2.92E-03 | 3.02E-02 |
| PRKAA1 | 1.4 | 2.93E-03 | 3.02E-02 |
| HS3ST3B1 | 4.0 | 2.93E-03 | 3.02E-02 |
| HNMT | 1.5 | 2.96E-03 | 3.04E-02 |
| GATA6 | 3.9 | 2.98E-03 | 3.05E-02 |
| ZC3H12D | 2.7 | 2.98E-03 | 3.06E-02 |
| TESPA1 | 3.0 | 2.98E-03 | 3.06E-02 |
| ASAP1 | 1.7 | 2.99E-03 | 3.06E-02 |
| GRIK2 | 1.7 | 3.00E-03 | 3.06E-02 |
| UIMC1 | 1.3 | 3.03E-03 | 3.09E-02 |
| TSPAN12 | 2.2 | 3.03E-03 | 3.09E-02 |
| SLC36A4 | 1.6 | 3.04E-03 | 3.09E-02 |
| FAM13A-AS1 | 1.4 | 3.04E-03 | 3.10E-02 |
| SQRDL | 2.0 | 3.06E-03 | 3.11E-02 |
| ATP8B4 | 2.1 | 3.07E-03 | 3.12E-02 |
| DNAJC6 | 2.5 | 3.09E-03 | 3.12E-02 |
| SLC15A3 | 2.4 | 3.09E-03 | 3.12E-02 |
| ATP2A3 | 2.8 | 3.10E-03 | 3.13E-02 |
| HAVCR2 | 2.8 | 3.11E-03 | 3.13E-02 |
| FPR1 | 7.8 | 3.11E-03 | 3.14E-02 |
| RHBDF2 | 2.5 | 3.13E-03 | 3.15E-02 |
| FBXW7 | 1.3 | 3.13E-03 | 3.15E-02 |
| KLHL5 | 1.6 | 3.15E-03 | 3.16E-02 |
| RAPGEF1 | 1.5 | 3.14E-03 | 3.16E-02 |
| GRB2 | 1.5 | 3.15E-03 | 3.16E-02 |
| LY9 | 3.1 | 3.18E-03 | 3.18E-02 |
| TBX19 | 1.6 | 3.18E-03 | 3.18E-02 |
| ADARB1 | 1.5 | 3.18E-03 | 3.18E-02 |
| ST5 | 1.7 | 3.25E-03 | 3.23E-02 |
| SIK3 | 1.6 | 3.28E-03 | 3.25E-02 |
| LY96 | 2.4 | 3.29E-03 | 3.25E-02 |
| HELZ | 1.3 | 3.30E-03 | 3.26E-02 |
| ADAMDEC1 | 3.4 | 3.32E-03 | 3.27E-02 |
| CASP4 | 1.8 | 3.34E-03 | 3.29E-02 |
| BCAR3 | 1.6 | 3.35E-03 | 3.29E-02 |
| S100A4 | 2.2 | 3.37E-03 | 3.30E-02 |
| SLC15A4 | 1.8 | 3.37E-03 | 3.30E-02 |
| RNF13 | 1.4 | 3.38E-03 | 3.30E-02 |
| FNIP1 | 1.4 | 3.37E-03 | 3.30E-02 |
| UBE2H | 1.4 | 3.38E-03 | 3.30E-02 |
| ACTR3 | 1.6 | 3.39E-03 | 3.31E-02 |
| WTAP | 1.4 | 3.39E-03 | 3.31E-02 |
| SH2B3 | 1.5 | 3.43E-03 | 3.34E-02 |
| IL8 | 10.0 | 3.46E-03 | 3.37E-02 |
| LINC00598 | 1.9 | 3.48E-03 | 3.38E-02 |
| LIPG | 2.2 | 3.49E-03 | 3.38E-02 |
| CYB5B | 1.5 | 3.54E-03 | 3.42E-02 |
| NSRP1 | 1.4 | 3.54E-03 | 3.42E-02 |
| PHF20 | 1.3 | 3.56E-03 | 3.44E-02 |
| BRAF | 1.4 | 3.56E-03 | 3.44E-02 |
| LAG3 | 3.7 | 3.57E-03 | 3.44E-02 |
| FBXO34 | 1.4 | 3.62E-03 | 3.48E-02 |
| SYNPO2 | 2.1 | 3.65E-03 | 3.50E-02 |
| PIK3CG | 2.2 | 3.66E-03 | 3.50E-02 |
| SVIL | 1.7 | 3.67E-03 | 3.51E-02 |
| DOCK11 | 2.1 | 3.68E-03 | 3.52E-02 |
| SGMS1 | 1.5 | 3.70E-03 | 3.52E-02 |
| ANTXR2 | 1.8 | 3.70E-03 | 3.52E-02 |
| TMTC2 | 1.3 | 3.71E-03 | 3.53E-02 |
| VSIG4 | 2.6 | 3.73E-03 | 3.54E-02 |
| AOAH | 2.5 | 3.75E-03 | 3.57E-02 |
| MMP25 | 3.0 | 3.77E-03 | 3.57E-02 |
| ADAM10 | 1.4 | 3.78E-03 | 3.58E-02 |
| ZNF708 | 1.2 | 3.78E-03 | 3.58E-02 |
| WDR26 | 1.3 | 3.79E-03 | 3.59E-02 |
| MPEG1 | 2.2 | 3.83E-03 | 3.61E-02 |
| FGD6 | 1.8 | 3.83E-03 | 3.61E-02 |
| TTC39B | 1.7 | 3.83E-03 | 3.61E-02 |
| RPS6KA3 | 1.6 | 3.83E-03 | 3.61E-02 |
| STX7 | 1.4 | 3.88E-03 | 3.64E-02 |
| MNDA | 5.8 | 3.90E-03 | 3.64E-02 |
| MYO7B | 2.4 | 3.90E-03 | 3.64E-02 |
| TFAP2C | 2.0 | 3.90E-03 | 3.64E-02 |
| CHSY3 | 1.8 | 3.90E-03 | 3.65E-02 |
| TPRG1 | 2.5 | 3.91E-03 | 3.65E-02 |
| FOXO1 | 2.2 | 3.92E-03 | 3.65E-02 |
| PTGER4 | 2.2 | 3.92E-03 | 3.65E-02 |
| NFKB2 | 2.0 | 3.93E-03 | 3.65E-02 |
| GMFG | 2.4 | 3.93E-03 | 3.65E-02 |
| LATS1 | 1.1 | 3.94E-03 | 3.66E-02 |
| CUL1 | 1.3 | 3.97E-03 | 3.68E-02 |
| NLGN4X | 2.1 | 3.98E-03 | 3.68E-02 |
| ZBTB1 | 1.2 | 3.99E-03 | 3.68E-02 |
| ENOX1 | 2.0 | 4.01E-03 | 3.69E-02 |
| TAPBP | 1.6 | 4.01E-03 | 3.69E-02 |
| ZNF277 | 1.3 | 4.01E-03 | 3.69E-02 |
| NFAM1 | 3.0 | 4.03E-03 | 3.71E-02 |
| TMEM108 | 2.1 | 4.04E-03 | 3.71E-02 |
| MRVI1-AS1 | 2.0 | 4.06E-03 | 3.73E-02 |
| PHACTR1 | 1.6 | 4.09E-03 | 3.75E-02 |
| THEMIS | 2.3 | 4.09E-03 | 3.75E-02 |
| MSR1 | 2.2 | 4.11E-03 | 3.76E-02 |
| ABCC8 | 6.0 | 4.14E-03 | 3.78E-02 |
| NTRK3 | 3.5 | 4.16E-03 | 3.79E-02 |
| CD80 | 6.1 | 4.16E-03 | 3.79E-02 |
| RALA | 1.3 | 4.17E-03 | 3.79E-02 |
| CFTR | 3.3 | 4.18E-03 | 3.80E-02 |
| UEVLD | 1.3 | 4.19E-03 | 3.80E-02 |
| SERPINA3 | 4.0 | 4.19E-03 | 3.80E-02 |
| HIVEP2 | 1.6 | 4.19E-03 | 3.80E-02 |
| SASH3 | 2.9 | 4.20E-03 | 3.80E-02 |
| USP25 | 1.5 | 4.21E-03 | 3.81E-02 |
| USH2A | 1.4 | 4.25E-03 | 3.83E-02 |
| CREBBP | 1.2 | 4.26E-03 | 3.83E-02 |
| DUOX2 | 3.8 | 4.27E-03 | 3.84E-02 |
| CD40LG | 2.9 | 4.28E-03 | 3.84E-02 |
| TRPC4 | 3.5 | 4.28E-03 | 3.84E-02 |
| KBTBD8 | 2.3 | 4.30E-03 | 3.85E-02 |
| IDO2 | 5.1 | 4.30E-03 | 3.86E-02 |
| MET | 1.9 | 4.31E-03 | 3.86E-02 |
| STARD13 | 1.8 | 4.34E-03 | 3.87E-02 |
| NFATC1 | 1.8 | 4.34E-03 | 3.88E-02 |
| LY86 | 2.1 | 4.35E-03 | 3.88E-02 |
| TRAPPC10 | 1.3 | 4.37E-03 | 3.89E-02 |
| FGD3 | 2.6 | 4.37E-03 | 3.89E-02 |
| PFKFB3 | 2.1 | 4.42E-03 | 3.92E-02 |
| RUNX2 | 2.2 | 4.42E-03 | 3.92E-02 |
| PHOSPHO1 | 3.2 | 4.44E-03 | 3.94E-02 |
| NCOA1 | 1.4 | 4.46E-03 | 3.95E-02 |
| CEBPB | 1.9 | 4.50E-03 | 3.98E-02 |
| DGKB | 4.0 | 4.52E-03 | 4.00E-02 |
| P2RY10 | 3.2 | 4.54E-03 | 4.01E-02 |
| ENTPD1 | 1.5 | 4.56E-03 | 4.01E-02 |
| ANKRD36BP1 | 1.5 | 4.62E-03 | 4.06E-02 |
| RSPO3 | 2.9 | 4.62E-03 | 4.06E-02 |
| LRCH3 | 1.4 | 4.64E-03 | 4.07E-02 |
| SRSF4 | 1.2 | 4.65E-03 | 4.07E-02 |
| CBL | 1.4 | 4.66E-03 | 4.08E-02 |
| SLC2A3 | 3.5 | 4.66E-03 | 4.08E-02 |
| EVC2 | 1.3 | 4.70E-03 | 4.11E-02 |
| PPP2R5A | 1.4 | 4.72E-03 | 4.12E-02 |
| EPS8 | 1.7 | 4.73E-03 | 4.12E-02 |
| TGM2 | 2.1 | 4.73E-03 | 4.12E-02 |
| TRAT1 | 3.2 | 4.73E-03 | 4.12E-02 |
| KDM6A | 1.4 | 4.76E-03 | 4.14E-02 |
| MOCOS | 1.7 | 4.78E-03 | 4.16E-02 |
| CEPT1 | 1.3 | 4.81E-03 | 4.17E-02 |
| TAOK1 | 1.2 | 4.81E-03 | 4.17E-02 |
| IRF1 | 2.5 | 4.82E-03 | 4.18E-02 |
| C1RL | 1.8 | 4.83E-03 | 4.18E-02 |
| KIAA2026 | 1.3 | 4.85E-03 | 4.20E-02 |
| RRN3P2 | 1.4 | 4.86E-03 | 4.20E-02 |
| USP31 | 1.6 | 4.87E-03 | 4.21E-02 |
| TET2 | 1.4 | 4.89E-03 | 4.22E-02 |
| BMP2K | 1.5 | 4.89E-03 | 4.22E-02 |
| LPGAT1 | 1.3 | 4.90E-03 | 4.22E-02 |
| ARHGDIB | 2.2 | 4.92E-03 | 4.23E-02 |
| TMED5 | 1.4 | 4.92E-03 | 4.23E-02 |
| TIGIT | 2.1 | 4.94E-03 | 4.24E-02 |
| PICALM | 1.6 | 4.94E-03 | 4.24E-02 |
| CD14 | 2.2 | 4.97E-03 | 4.25E-02 |
| SMAD1 | 1.2 | 4.98E-03 | 4.26E-02 |
| UPF2 | 1.2 | 4.99E-03 | 4.26E-02 |
| AIDA | 1.3 | 4.99E-03 | 4.26E-02 |
| RHD | 2.3 | 5.00E-03 | 4.27E-02 |
| DOCK9 | 1.2 | 5.02E-03 | 4.28E-02 |
| AFF3 | 1.5 | 5.03E-03 | 4.28E-02 |
| RAB5A | 1.3 | 5.03E-03 | 4.28E-02 |
| DUSP10 | 2.0 | 5.05E-03 | 4.29E-02 |
| UHRF1BP1L | 1.3 | 5.08E-03 | 4.30E-02 |
| CMTM2 | 2.7 | 5.07E-03 | 4.30E-02 |
| ZCCHC2 | 1.7 | 5.07E-03 | 4.30E-02 |
| JAK3 | 3.1 | 5.07E-03 | 4.30E-02 |
| MMAA | 1.4 | 5.08E-03 | 4.30E-02 |
| STOM | 2.0 | 5.07E-03 | 4.30E-02 |
| RAB9A | 1.4 | 5.07E-03 | 4.30E-02 |
| SLC46A3 | 1.9 | 5.10E-03 | 4.31E-02 |
| NCKAP1L | 2.5 | 5.12E-03 | 4.32E-02 |
| C7orf10 | 2.0 | 5.12E-03 | 4.32E-02 |
| MAP4K1 | 2.9 | 5.14E-03 | 4.33E-02 |
| C12orf75 | 1.8 | 5.18E-03 | 4.34E-02 |
| IRS2 | 2.3 | 5.17E-03 | 4.34E-02 |
| SLC4A5 | 1.4 | 5.17E-03 | 4.34E-02 |
| ITPR1 | 1.7 | 5.17E-03 | 4.34E-02 |
| RIPK1 | 1.4 | 5.18E-03 | 4.34E-02 |
| CCR6 | 2.6 | 5.17E-03 | 4.34E-02 |
| CLEC4A | 2.0 | 5.20E-03 | 4.35E-02 |
| EFHD2 | 1.7 | 5.22E-03 | 4.36E-02 |
| EHD1 | 1.6 | 5.24E-03 | 4.37E-02 |
| RASA3 | 1.6 | 5.24E-03 | 4.37E-02 |
| TNRC6A | 1.3 | 5.24E-03 | 4.37E-02 |
| DGKG | 2.2 | 5.26E-03 | 4.38E-02 |
| ATXN3 | 1.3 | 5.28E-03 | 4.39E-02 |
| MUC13 | 2.6 | 5.31E-03 | 4.42E-02 |
| SLPI | 3.0 | 5.35E-03 | 4.44E-02 |
| BIN2 | 2.9 | 5.36E-03 | 4.45E-02 |
| EVI2A | 2.1 | 5.42E-03 | 4.47E-02 |
| FAM63B | 1.4 | 5.43E-03 | 4.48E-02 |
| SLC12A6 | 1.4 | 5.50E-03 | 4.51E-02 |
| ARHGAP9 | 2.7 | 5.57E-03 | 4.54E-02 |
| SLC16A7 | 1.7 | 5.56E-03 | 4.54E-02 |
| NOD2 | 2.5 | 5.56E-03 | 4.54E-02 |
| TFEB | 1.9 | 5.56E-03 | 4.54E-02 |
| GPR155 | 1.8 | 5.60E-03 | 4.56E-02 |
| PSAT1 | 2.4 | 5.60E-03 | 4.56E-02 |
| RNF145 | 1.7 | 5.63E-03 | 4.58E-02 |
| LOC100505702 | 2.0 | 5.64E-03 | 4.59E-02 |
| HECA | 1.5 | 5.65E-03 | 4.59E-02 |
| GRAP2 | 2.8 | 5.67E-03 | 4.59E-02 |
| JAK1 | 1.3 | 5.69E-03 | 4.61E-02 |
| PLIN5 | 1.9 | 5.70E-03 | 4.61E-02 |
| PPARD | 1.3 | 5.69E-03 | 4.61E-02 |
| TRAM2 | 1.4 | 5.69E-03 | 4.61E-02 |
| FAM134B | 1.7 | 5.70E-03 | 4.61E-02 |
| TANK | 1.5 | 5.71E-03 | 4.61E-02 |
| GNPTAB | 1.4 | 5.72E-03 | 4.61E-02 |
| C16orf52 | 1.7 | 5.74E-03 | 4.62E-02 |
| SLC16A3 | 2.2 | 5.74E-03 | 4.62E-02 |
| KANSL1 | 1.3 | 5.75E-03 | 4.63E-02 |
| MICALCL | 2.2 | 5.77E-03 | 4.64E-02 |
| CD5 | 2.4 | 5.78E-03 | 4.64E-02 |
| USP8 | 1.2 | 5.79E-03 | 4.65E-02 |
| ARAP2 | 1.7 | 5.85E-03 | 4.69E-02 |
| ZBTB16 | 6.0 | 5.87E-03 | 4.70E-02 |
| HERC3 | 1.3 | 5.88E-03 | 4.70E-02 |
| TOX | 1.9 | 5.88E-03 | 4.70E-02 |
| CD3D | 3.2 | 5.92E-03 | 4.72E-02 |
| CCR2 | 2.8 | 5.92E-03 | 4.72E-02 |
| NFKBID | 1.7 | 5.93E-03 | 4.73E-02 |
| NRG1 | 3.2 | 5.95E-03 | 4.74E-02 |
| WWTR1 | 1.6 | 5.96E-03 | 4.74E-02 |
| FKBP15 | 1.3 | 5.98E-03 | 4.75E-02 |
| TASP1 | 1.2 | 6.00E-03 | 4.75E-02 |
| DHCR24 | 1.9 | 6.01E-03 | 4.76E-02 |
| SYT15 | 1.5 | 6.09E-03 | 4.80E-02 |
| EPS15 | 1.5 | 6.09E-03 | 4.81E-02 |
| C19orf38 | 2.8 | 6.09E-03 | 4.81E-02 |
| C6orf106 | 1.3 | 6.11E-03 | 4.81E-02 |
| UBAP1 | 1.3 | 6.12E-03 | 4.81E-02 |
| EP300 | 1.2 | 6.16E-03 | 4.85E-02 |
| CDA | 3.4 | 6.19E-03 | 4.86E-02 |
| NRF1 | 1.3 | 6.20E-03 | 4.87E-02 |
| ZNF655 | 1.2 | 6.21E-03 | 4.87E-02 |
| DYRK2 | 1.5 | 6.23E-03 | 4.88E-02 |
| SNTB2 | 1.4 | 6.23E-03 | 4.88E-02 |
| FBXO11 | 1.3 | 6.28E-03 | 4.91E-02 |
| PPP1R16B | 2.4 | 6.28E-03 | 4.91E-02 |
| PPAP2B | 2.2 | 6.31E-03 | 4.92E-02 |
| RC3H1 | 1.2 | 6.31E-03 | 4.92E-02 |
| KCMF1 | 1.3 | 6.31E-03 | 4.92E-02 |
| SLC35G2 | 1.4 | 6.30E-03 | 4.92E-02 |
| ARHGAP12 | 1.3 | 6.33E-03 | 4.92E-02 |
| CHST15 | 1.8 | 6.33E-03 | 4.92E-02 |
| PRKCA | 1.9 | 6.33E-03 | 4.92E-02 |
| TNFSF13B | 1.7 | 6.36E-03 | 4.94E-02 |
| TAB2 | 1.3 | 6.39E-03 | 4.96E-02 |
| ZHX1-C8ORF76 | 1.5 | 6.42E-03 | 4.97E-02 |
| SNX25 | 1.3 | 6.45E-03 | 5.00E-02 |
