## Supplemental Table 1-10 for "miRNA/mRNA analysis of increased TGF-β pathways drive epithelial-mesenchymal transition and regulatory T cell differentiation": Supp Table 4.docx

| **Supp Table 4. Downregulated mRNAs in Endo+ compared to Endo- women with FDR <0.05.**  **(sorted by FDR values)** | | | |
| --- | --- | --- | --- |
| **Gene symbol** | **Fold Change** | **P value** | **FDR** |
| MDK | 0.41 | 1.86E-12 | 9.63E-09 |
| KATNB1 | 0.62 | 3.02E-12 | 1.17E-08 |
| C11orf31 | 0.58 | 2.66E-10 | 4.57E-07 |
| HIST1H1A | 0.25 | 5.57E-10 | 7.84E-07 |
| APLP1 | 0.27 | 6.68E-10 | 7.91E-07 |
| C19orf48 | 0.35 | 7.66E-10 | 7.91E-07 |
| BCL7C | 0.59 | 2.53E-09 | 1.87E-06 |
| GSTZ1 | 0.50 | 1.26E-08 | 6.72E-06 |
| HIST1H4D | 0.36 | 2.07E-08 | 1.00E-05 |
| DNAJC22 | 0.53 | 2.31E-08 | 1.02E-05 |
| MYL6B | 0.53 | 2.29E-08 | 1.02E-05 |
| HIST1H3A | 0.29 | 2.94E-08 | 1.23E-05 |
| WDR18 | 0.59 | 3.60E-08 | 1.47E-05 |
| PYCR1 | 0.43 | 8.19E-08 | 2.95E-05 |
| FBN3 | 0.25 | 1.09E-07 | 3.54E-05 |
| ID3 | 0.53 | 1.10E-07 | 3.54E-05 |
| HIST1H2AI | 0.28 | 1.42E-07 | 4.11E-05 |
| TNPO2 | 0.78 | 1.43E-07 | 4.11E-05 |
| TRIM28 | 0.65 | 1.48E-07 | 4.14E-05 |
| STRA13 | 0.43 | 1.70E-07 | 4.53E-05 |
| RRP7B | 0.58 | 1.84E-07 | 4.84E-05 |
| DGCR6L | 0.62 | 2.58E-07 | 6.35E-05 |
| H2AFX | 0.30 | 2.94E-07 | 6.91E-05 |
| HINT2 | 0.51 | 2.91E-07 | 6.91E-05 |
| HIST2H3D | 0.27 | 3.04E-07 | 7.04E-05 |
| BOK | 0.55 | 3.23E-07 | 7.20E-05 |
| R3HCC1 | 0.63 | 3.38E-07 | 7.28E-05 |
| C16orf59 | 0.31 | 3.54E-07 | 7.40E-05 |
| NDUFB7 | 0.54 | 3.84E-07 | 7.72E-05 |
| TBL3 | 0.67 | 3.95E-07 | 7.75E-05 |
| HIST1H3F | 0.30 | 4.60E-07 | 8.90E-05 |
| SLC2A4RG | 0.47 | 4.70E-07 | 8.99E-05 |
| CDK4 | 0.54 | 4.86E-07 | 9.18E-05 |
| MTRNR2L8 | 0.11 | 5.41E-07 | 9.62E-05 |
| PSMG3 | 0.49 | 5.55E-07 | 9.77E-05 |
| HIST1H3D | 0.36 | 6.38E-07 | 1.09E-04 |
| PTRHD1 | 0.50 | 6.39E-07 | 1.09E-04 |
| GPAA1 | 0.65 | 6.70E-07 | 1.12E-04 |
| RNASEH2A | 0.38 | 7.56E-07 | 1.22E-04 |
| XKR5 | 0.41 | 8.26E-07 | 1.29E-04 |
| LAMTOR4 | 0.60 | 8.86E-07 | 1.36E-04 |
| KIF22 | 0.43 | 1.03E-06 | 1.55E-04 |
| LIG1 | 0.48 | 1.30E-06 | 1.85E-04 |
| PRDX2 | 0.59 | 1.39E-06 | 1.95E-04 |
| APRT | 0.50 | 1.42E-06 | 1.96E-04 |
| SAPCD2 | 0.26 | 1.42E-06 | 1.96E-04 |
| COMMD4 | 0.63 | 1.51E-06 | 2.02E-04 |
| HIST1H2AB | 0.28 | 1.64E-06 | 2.11E-04 |
| HIST1H2AE | 0.35 | 1.73E-06 | 2.17E-04 |
| CUTA | 0.57 | 1.82E-06 | 2.27E-04 |
| LRRC45 | 0.46 | 2.08E-06 | 2.50E-04 |
| LRP3 | 0.58 | 2.23E-06 | 2.66E-04 |
| POLD1 | 0.55 | 2.29E-06 | 2.70E-04 |
| SLC9A2 | 0.31 | 2.30E-06 | 2.70E-04 |
| HMG20B | 0.54 | 2.53E-06 | 2.92E-04 |
| NDUFB11 | 0.61 | 2.61E-06 | 2.97E-04 |
| GALNT12 | 0.37 | 2.85E-06 | 3.18E-04 |
| CD320 | 0.47 | 3.09E-06 | 3.34E-04 |
| ATP6V0E2 | 0.35 | 3.16E-06 | 3.38E-04 |
| LRRC14 | 0.65 | 3.33E-06 | 3.50E-04 |
| SIVA1 | 0.56 | 3.30E-06 | 3.50E-04 |
| GJB6 | 0.07 | 3.73E-06 | 3.81E-04 |
| PLOD1 | 0.70 | 3.77E-06 | 3.81E-04 |
| ENDOG | 0.26 | 4.12E-06 | 4.09E-04 |
| HIST1H3H | 0.37 | 4.22E-06 | 4.14E-04 |
| MTRNR2L4 | 0.16 | 4.57E-06 | 4.42E-04 |
| BANF1 | 0.60 | 4.68E-06 | 4.50E-04 |
| ARHGEF39 | 0.28 | 4.94E-06 | 4.67E-04 |
| SEMA3E | 0.14 | 5.11E-06 | 4.73E-04 |
| GTF2H4 | 0.69 | 5.36E-06 | 4.74E-04 |
| KAZALD1 | 0.30 | 5.25E-06 | 4.74E-04 |
| UBL7 | 0.66 | 5.61E-06 | 4.88E-04 |
| HIST1H4B | 0.44 | 5.66E-06 | 4.90E-04 |
| HIST1H4A | 0.29 | 5.73E-06 | 4.92E-04 |
| LINC00261 | 0.22 | 5.75E-06 | 4.92E-04 |
| HIST1H2BN | 0.35 | 5.85E-06 | 4.97E-04 |
| HIST1H1E | 0.50 | 6.16E-06 | 5.19E-04 |
| HIST1H2BB | 0.20 | 6.29E-06 | 5.21E-04 |
| TMEM106C | 0.44 | 6.63E-06 | 5.46E-04 |
| METTL1 | 0.66 | 6.73E-06 | 5.51E-04 |
| YDJC | 0.56 | 7.50E-06 | 5.93E-04 |
| TIMM13 | 0.59 | 7.55E-06 | 5.94E-04 |
| NARFL | 0.67 | 8.51E-06 | 6.59E-04 |
| HIST1H3C | 0.30 | 9.63E-06 | 7.38E-04 |
| BRAT1 | 0.75 | 1.02E-05 | 7.60E-04 |
| HIST1H1B | 0.29 | 1.02E-05 | 7.60E-04 |
| HIST1H3G | 0.26 | 1.02E-05 | 7.60E-04 |
| LMF2 | 0.73 | 1.12E-05 | 8.21E-04 |
| MCRS1 | 0.71 | 1.18E-05 | 8.40E-04 |
| HDDC2 | 0.71 | 1.19E-05 | 8.49E-04 |
| HIST1H4F | 0.46 | 1.22E-05 | 8.51E-04 |
| CDCA3 | 0.32 | 1.24E-05 | 8.53E-04 |
| FPGT-TNNI3K | 0.00 | 1.26E-05 | 8.61E-04 |
| GRB14 | 0.25 | 1.30E-05 | 8.72E-04 |
| HMBS | 0.61 | 1.36E-05 | 8.95E-04 |
| PHGDH | 0.38 | 1.38E-05 | 8.99E-04 |
| GLB1L2 | 0.48 | 1.43E-05 | 9.26E-04 |
| STK25 | 0.69 | 1.45E-05 | 9.29E-04 |
| GPS1 | 0.66 | 1.48E-05 | 9.38E-04 |
| HIST1H3B | 0.28 | 1.65E-05 | 1.03E-03 |
| ARHGEF19 | 0.43 | 1.71E-05 | 1.05E-03 |
| PGP | 0.57 | 1.79E-05 | 1.08E-03 |
| ERGIC3 | 0.70 | 1.86E-05 | 1.12E-03 |
| HIST1H2AG | 0.34 | 1.87E-05 | 1.12E-03 |
| CAD | 0.71 | 1.91E-05 | 1.14E-03 |
| LMNB2 | 0.45 | 1.91E-05 | 1.14E-03 |
| EMID1 | 0.42 | 1.97E-05 | 1.15E-03 |
| TIMELESS | 0.51 | 1.98E-05 | 1.15E-03 |
| TP53I3 | 0.68 | 1.99E-05 | 1.15E-03 |
| CDT1 | 0.36 | 2.02E-05 | 1.15E-03 |
| MSX1 | 0.39 | 2.05E-05 | 1.17E-03 |
| TELO2 | 0.66 | 2.06E-05 | 1.17E-03 |
| HIST4H4 | 0.55 | 2.22E-05 | 1.23E-03 |
| SEZ6L2 | 0.61 | 2.26E-05 | 1.24E-03 |
| HAUS5 | 0.59 | 2.34E-05 | 1.28E-03 |
| C9orf142 | 0.57 | 2.37E-05 | 1.28E-03 |
| CDC20 | 0.31 | 2.37E-05 | 1.28E-03 |
| GEMIN4 | 0.75 | 2.37E-05 | 1.28E-03 |
| DHFR | 0.16 | 2.43E-05 | 1.30E-03 |
| HIST1H2AJ | 0.32 | 2.47E-05 | 1.32E-03 |
| PSRC1 | 0.34 | 2.48E-05 | 1.32E-03 |
| FAM64A | 0.21 | 2.55E-05 | 1.34E-03 |
| HMGB3 | 0.47 | 2.56E-05 | 1.35E-03 |
| DTYMK | 0.59 | 2.67E-05 | 1.37E-03 |
| HIST1H2BD | 0.44 | 2.69E-05 | 1.38E-03 |
| CHTF18 | 0.40 | 2.76E-05 | 1.39E-03 |
| CLEC11A | 0.53 | 2.88E-05 | 1.40E-03 |
| EDAR | 0.31 | 2.88E-05 | 1.40E-03 |
| MRPL11 | 0.61 | 2.87E-05 | 1.40E-03 |
| NDUFA11 | 0.61 | 2.88E-05 | 1.40E-03 |
| SLC25A10 | 0.47 | 3.02E-05 | 1.46E-03 |
| UQCRC1 | 0.66 | 3.03E-05 | 1.46E-03 |
| XRCC1 | 0.68 | 3.05E-05 | 1.46E-03 |
| MARCKSL1 | 0.55 | 3.23E-05 | 1.53E-03 |
| TROAP | 0.26 | 3.31E-05 | 1.56E-03 |
| C9orf114 | 0.60 | 3.39E-05 | 1.57E-03 |
| HIST1H2BF | 0.29 | 3.38E-05 | 1.57E-03 |
| FANCG | 0.56 | 3.42E-05 | 1.57E-03 |
| BOD1 | 0.73 | 3.45E-05 | 1.58E-03 |
| MMP26 | 0.08 | 3.46E-05 | 1.58E-03 |
| ASF1B | 0.34 | 3.57E-05 | 1.62E-03 |
| EDC4 | 0.73 | 3.61E-05 | 1.64E-03 |
| MRAP2 | 0.24 | 3.83E-05 | 1.72E-03 |
| NME1 | 0.45 | 3.87E-05 | 1.73E-03 |
| HIST1H2BE | 0.38 | 3.94E-05 | 1.75E-03 |
| PCDH10 | 0.18 | 3.95E-05 | 1.75E-03 |
| DAK | 0.56 | 4.19E-05 | 1.83E-03 |
| JTB | 0.56 | 4.29E-05 | 1.87E-03 |
| HIST1H2BG | 0.32 | 4.36E-05 | 1.89E-03 |
| SORCS2 | 0.32 | 4.43E-05 | 1.91E-03 |
| ATP5G2 | 0.72 | 4.55E-05 | 1.94E-03 |
| POLR2G | 0.70 | 4.60E-05 | 1.94E-03 |
| THYN1 | 0.65 | 4.63E-05 | 1.94E-03 |
| CITED4 | 0.34 | 4.79E-05 | 1.97E-03 |
| NHP2 | 0.63 | 4.82E-05 | 1.98E-03 |
| HIST1H1D | 0.43 | 4.87E-05 | 1.99E-03 |
| RXRG | 0.24 | 4.89E-05 | 1.99E-03 |
| TSEN34 | 0.64 | 4.89E-05 | 1.99E-03 |
| CCDC22 | 0.65 | 5.03E-05 | 2.02E-03 |
| HIST1H2BI | 0.38 | 5.05E-05 | 2.03E-03 |
| DUS3L | 0.68 | 5.33E-05 | 2.10E-03 |
| AP2S1 | 0.60 | 5.35E-05 | 2.10E-03 |
| TMED3 | 0.59 | 5.75E-05 | 2.21E-03 |
| OGFOD2 | 0.64 | 6.01E-05 | 2.29E-03 |
| C17orf89 | 0.59 | 6.18E-05 | 2.32E-03 |
| DBN1 | 0.62 | 6.26E-05 | 2.34E-03 |
| NME4 | 0.47 | 6.28E-05 | 2.34E-03 |
| PAQR4 | 0.33 | 6.43E-05 | 2.39E-03 |
| SCD5 | 0.51 | 6.47E-05 | 2.40E-03 |
| NUDT1 | 0.53 | 6.53E-05 | 2.42E-03 |
| SOGA2 | 0.25 | 6.57E-05 | 2.43E-03 |
| ARL6IP4 | 0.64 | 6.79E-05 | 2.49E-03 |
| SPC24 | 0.36 | 6.78E-05 | 2.49E-03 |
| PKN3 | 0.49 | 7.03E-05 | 2.56E-03 |
| DDX51 | 0.71 | 7.09E-05 | 2.57E-03 |
| RPL36 | 0.51 | 7.28E-05 | 2.63E-03 |
| FAM83D | 0.34 | 7.37E-05 | 2.64E-03 |
| WFS1 | 0.64 | 7.43E-05 | 2.65E-03 |
| ISOC2 | 0.56 | 7.49E-05 | 2.65E-03 |
| AURKB | 0.34 | 7.54E-05 | 2.66E-03 |
| WDR83OS | 0.64 | 7.77E-05 | 2.73E-03 |
| DVL2 | 0.72 | 7.94E-05 | 2.77E-03 |
| MFSD3 | 0.63 | 7.92E-05 | 2.77E-03 |
| C17orf53 | 0.49 | 8.02E-05 | 2.80E-03 |
| B4GALT2 | 0.65 | 8.23E-05 | 2.85E-03 |
| ZWINT | 0.32 | 8.24E-05 | 2.85E-03 |
| C1orf174 | 0.74 | 8.34E-05 | 2.88E-03 |
| SPDEF | 0.20 | 8.39E-05 | 2.89E-03 |
| ECI1 | 0.49 | 8.52E-05 | 2.93E-03 |
| CCNB1 | 0.33 | 8.58E-05 | 2.94E-03 |
| RASL11B | 0.50 | 8.77E-05 | 2.99E-03 |
| THAP7 | 0.61 | 8.83E-05 | 3.01E-03 |
| FAM69B | 0.49 | 8.93E-05 | 3.03E-03 |
| RAC3 | 0.35 | 9.37E-05 | 3.16E-03 |
| THOP1 | 0.61 | 9.42E-05 | 3.17E-03 |
| RBM42 | 0.71 | 9.47E-05 | 3.17E-03 |
| BIRC5 | 0.29 | 9.56E-05 | 3.19E-03 |
| HEY2 | 0.34 | 9.71E-05 | 3.22E-03 |
| SLC25A1 | 0.45 | 9.73E-05 | 3.22E-03 |
| FAAH | 0.59 | 9.85E-05 | 3.24E-03 |
| IHH | 0.22 | 1.04E-04 | 3.40E-03 |
| CREB3L4 | 0.40 | 1.05E-04 | 3.41E-03 |
| RPS15 | 0.60 | 1.05E-04 | 3.41E-03 |
| CDK16 | 0.73 | 1.05E-04 | 3.41E-03 |
| POC1A | 0.40 | 1.08E-04 | 3.50E-03 |
| CUL7 | 0.70 | 1.09E-04 | 3.50E-03 |
| DHRS11 | 0.52 | 1.09E-04 | 3.50E-03 |
| HIST1H2BH | 0.37 | 1.10E-04 | 3.51E-03 |
| HIST1H4C | 0.47 | 1.10E-04 | 3.51E-03 |
| LINGO1 | 0.24 | 1.10E-04 | 3.51E-03 |
| CENPM | 0.41 | 1.13E-04 | 3.57E-03 |
| GGT6 | 0.35 | 1.12E-04 | 3.57E-03 |
| GTF2H2C | 0.58 | 1.12E-04 | 3.57E-03 |
| NDUFS8 | 0.61 | 1.15E-04 | 3.63E-03 |
| MSX2 | 0.35 | 1.16E-04 | 3.64E-03 |
| HIST1H2AK | 0.43 | 1.21E-04 | 3.75E-03 |
| MRPL55 | 0.59 | 1.23E-04 | 3.80E-03 |
| PPP2R1A | 0.76 | 1.24E-04 | 3.81E-03 |
| CTSL2 | 0.22 | 1.30E-04 | 3.96E-03 |
| G6PC3 | 0.63 | 1.30E-04 | 3.96E-03 |
| INTS1 | 0.71 | 1.30E-04 | 3.96E-03 |
| NRCAM | 0.43 | 1.30E-04 | 3.96E-03 |
| THOC6 | 0.62 | 1.30E-04 | 3.96E-03 |
| AP4M1 | 0.71 | 1.31E-04 | 3.97E-03 |
| SNRPA | 0.65 | 1.31E-04 | 3.97E-03 |
| PMM1 | 0.62 | 1.33E-04 | 3.99E-03 |
| HFM1 | 0.20 | 1.36E-04 | 4.08E-03 |
| ELP6 | 0.70 | 1.39E-04 | 4.16E-03 |
| TRAPPC2L | 0.65 | 1.41E-04 | 4.18E-03 |
| APEX1 | 0.62 | 1.43E-04 | 4.24E-03 |
| HEY1 | 0.28 | 1.43E-04 | 4.24E-03 |
| IQGAP3 | 0.34 | 1.44E-04 | 4.24E-03 |
| DDX41 | 0.70 | 1.45E-04 | 4.27E-03 |
| NEK2 | 0.29 | 1.45E-04 | 4.27E-03 |
| GINS2 | 0.45 | 1.46E-04 | 4.28E-03 |
| SLC37A4 | 0.56 | 1.47E-04 | 4.31E-03 |
| ARMCX6 | 0.52 | 1.60E-04 | 4.58E-03 |
| KIF20A | 0.27 | 1.60E-04 | 4.58E-03 |
| MCM2 | 0.44 | 1.65E-04 | 4.69E-03 |
| RECQL4 | 0.40 | 1.65E-04 | 4.69E-03 |
| RUVBL2 | 0.62 | 1.65E-04 | 4.69E-03 |
| TMEM98 | 0.62 | 1.67E-04 | 4.71E-03 |
| GRHPR | 0.72 | 1.69E-04 | 4.74E-03 |
| KRTCAP3 | 0.51 | 1.68E-04 | 4.74E-03 |
| PKMYT1 | 0.31 | 1.69E-04 | 4.74E-03 |
| VTCN1 | 0.34 | 1.70E-04 | 4.77E-03 |
| PIF1 | 0.40 | 1.77E-04 | 4.92E-03 |
| KIFC1 | 0.35 | 1.82E-04 | 5.03E-03 |
| FIBP | 0.63 | 1.83E-04 | 5.05E-03 |
| PUF60 | 0.71 | 1.83E-04 | 5.05E-03 |
| NLGN3 | 0.47 | 1.88E-04 | 5.14E-03 |
| GUSB | 0.42 | 1.91E-04 | 5.23E-03 |
| HIST1H2BC | 0.37 | 1.96E-04 | 5.33E-03 |
| CCNA2 | 0.34 | 1.98E-04 | 5.37E-03 |
| RAB34 | 0.64 | 1.98E-04 | 5.37E-03 |
| PLEKHJ1 | 0.59 | 2.00E-04 | 5.38E-03 |
| C16orf13 | 0.61 | 2.01E-04 | 5.38E-03 |
| ARHGAP33 | 0.52 | 2.06E-04 | 5.48E-03 |
| TPGS2 | 0.74 | 2.06E-04 | 5.48E-03 |
| NPM3 | 0.62 | 2.10E-04 | 5.56E-03 |
| OPN3 | 0.43 | 2.13E-04 | 5.62E-03 |
| NR2C2AP | 0.58 | 2.15E-04 | 5.67E-03 |
| ATP5D | 0.55 | 2.16E-04 | 5.68E-03 |
| CDO1 | 0.55 | 2.24E-04 | 5.87E-03 |
| NT5DC2 | 0.48 | 2.24E-04 | 5.87E-03 |
| CDCA4 | 0.56 | 2.26E-04 | 5.89E-03 |
| FAM159A | 0.42 | 2.27E-04 | 5.90E-03 |
| POP5 | 0.73 | 2.27E-04 | 5.90E-03 |
| FAM5B | 0.23 | 2.28E-04 | 5.90E-03 |
| PGAP3 | 0.66 | 2.29E-04 | 5.91E-03 |
| SOX17 | 0.42 | 2.31E-04 | 5.94E-03 |
| COX8A | 0.63 | 2.36E-04 | 6.03E-03 |
| POLRMT | 0.75 | 2.36E-04 | 6.03E-03 |
| MRPL51 | 0.64 | 2.43E-04 | 6.20E-03 |
| SURF1 | 0.68 | 2.44E-04 | 6.21E-03 |
| MAGEF1 | 0.68 | 2.51E-04 | 6.35E-03 |
| MMAB | 0.55 | 2.56E-04 | 6.45E-03 |
| NDUFV1 | 0.73 | 2.64E-04 | 6.57E-03 |
| WFDC1 | 0.32 | 2.64E-04 | 6.57E-03 |
| UXT | 0.71 | 2.66E-04 | 6.60E-03 |
| CKAP4 | 0.65 | 2.68E-04 | 6.62E-03 |
| HIST1H1C | 0.49 | 2.68E-04 | 6.62E-03 |
| LEPREL4 | 0.65 | 2.69E-04 | 6.62E-03 |
| MZT2B | 0.51 | 2.67E-04 | 6.62E-03 |
| MAD2L2 | 0.61 | 2.73E-04 | 6.69E-03 |
| TRAIP | 0.46 | 2.73E-04 | 6.69E-03 |
| PIAS3 | 0.60 | 2.75E-04 | 6.71E-03 |
| WDR90 | 0.54 | 2.78E-04 | 6.72E-03 |
| PCBD1 | 0.64 | 2.86E-04 | 6.86E-03 |
| PRPF19 | 0.70 | 2.89E-04 | 6.91E-03 |
| PTOV1 | 0.70 | 2.89E-04 | 6.91E-03 |
| CHGA | 0.35 | 2.94E-04 | 6.95E-03 |
| EFCAB11 | 0.58 | 2.97E-04 | 6.96E-03 |
| HDAC6 | 0.80 | 2.96E-04 | 6.96E-03 |
| MRPL23 | 0.65 | 2.97E-04 | 6.96E-03 |
| GHDC | 0.69 | 3.08E-04 | 7.20E-03 |
| KDELR3 | 0.64 | 3.12E-04 | 7.22E-03 |
| PTTG1 | 0.36 | 3.12E-04 | 7.22E-03 |
| UBE2C | 0.33 | 3.12E-04 | 7.22E-03 |
| NABP2 | 0.66 | 3.16E-04 | 7.25E-03 |
| PRC1 | 0.34 | 3.19E-04 | 7.31E-03 |
| ACE | 0.32 | 3.24E-04 | 7.35E-03 |
| PRSS8 | 0.50 | 3.24E-04 | 7.35E-03 |
| YIPF2 | 0.65 | 3.33E-04 | 7.52E-03 |
| CDCA5 | 0.35 | 3.41E-04 | 7.63E-03 |
| NPAS3 | 0.31 | 3.41E-04 | 7.63E-03 |
| DLX5 | 0.53 | 3.43E-04 | 7.65E-03 |
| MCM3 | 0.56 | 3.47E-04 | 7.72E-03 |
| MLF2 | 0.75 | 3.51E-04 | 7.77E-03 |
| HIST1H4H | 0.49 | 3.53E-04 | 7.81E-03 |
| HIST1H2AM | 0.43 | 3.55E-04 | 7.83E-03 |
| HIST2H2BF | 0.50 | 3.55E-04 | 7.83E-03 |
| HJURP | 0.35 | 3.60E-04 | 7.88E-03 |
| MAN1B1 | 0.72 | 3.61E-04 | 7.89E-03 |
| JMJD8 | 0.66 | 3.67E-04 | 7.98E-03 |
| MSH2 | 0.67 | 3.70E-04 | 8.03E-03 |
| PFAS | 0.71 | 3.77E-04 | 8.17E-03 |
| GNAS | 0.78 | 3.81E-04 | 8.23E-03 |
| PEG10 | 0.34 | 3.87E-04 | 8.32E-03 |
| MZT2A | 0.53 | 3.97E-04 | 8.50E-03 |
| NMB | 0.58 | 3.98E-04 | 8.52E-03 |
| NPDC1 | 0.46 | 4.01E-04 | 8.53E-03 |
| TADA3 | 0.76 | 4.00E-04 | 8.53E-03 |
| EARS2 | 0.77 | 4.03E-04 | 8.56E-03 |
| UBTF | 0.76 | 4.04E-04 | 8.57E-03 |
| STMN1 | 0.50 | 4.12E-04 | 8.69E-03 |
| DDX49 | 0.68 | 4.13E-04 | 8.70E-03 |
| CDCA2 | 0.36 | 4.23E-04 | 8.88E-03 |
| KIF2C | 0.41 | 4.23E-04 | 8.88E-03 |
| ULK3 | 0.62 | 4.24E-04 | 8.88E-03 |
| ZNF530 | 0.71 | 4.26E-04 | 8.91E-03 |
| ECE2 | 0.57 | 4.28E-04 | 8.94E-03 |
| CENPA | 0.32 | 4.30E-04 | 8.95E-03 |
| DUS1L | 0.69 | 4.34E-04 | 9.01E-03 |
| HID1 | 0.50 | 4.35E-04 | 9.03E-03 |
| KDELR1 | 0.71 | 4.39E-04 | 9.06E-03 |
| RAB1B | 0.70 | 4.39E-04 | 9.06E-03 |
| RRP9 | 0.68 | 4.47E-04 | 9.19E-03 |
| RPS21 | 0.61 | 4.53E-04 | 9.25E-03 |
| ARFIP2 | 0.66 | 4.58E-04 | 9.32E-03 |
| C21orf58 | 0.47 | 4.68E-04 | 9.46E-03 |
| HTRA2 | 0.65 | 4.69E-04 | 9.46E-03 |
| ARF5 | 0.65 | 4.71E-04 | 9.48E-03 |
| HOXA11-AS | 0.61 | 4.71E-04 | 9.48E-03 |
| GID8 | 0.81 | 4.73E-04 | 9.49E-03 |
| MXD3 | 0.42 | 4.75E-04 | 9.53E-03 |
| CENPF | 0.34 | 4.88E-04 | 9.73E-03 |
| SCAND1 | 0.56 | 4.88E-04 | 9.73E-03 |
| CRIP2 | 0.61 | 4.95E-04 | 9.75E-03 |
| HIST1H3E | 0.52 | 4.94E-04 | 9.75E-03 |
| PRR7 | 0.46 | 4.94E-04 | 9.75E-03 |
| SCARF2 | 0.60 | 4.92E-04 | 9.75E-03 |
| SLC35B2 | 0.62 | 4.96E-04 | 9.75E-03 |
| CTXN1 | 0.44 | 4.99E-04 | 9.79E-03 |
| COPE | 0.64 | 5.01E-04 | 9.82E-03 |
| ACTR1B | 0.75 | 5.08E-04 | 9.92E-03 |
| NDUFA13 | 0.63 | 5.09E-04 | 9.92E-03 |
| LOC643623 | 0.27 | 5.10E-04 | 9.94E-03 |
| GAMT | 0.54 | 5.32E-04 | 1.03E-02 |
| LEPREL2 | 0.64 | 5.33E-04 | 1.03E-02 |
| GSS | 0.68 | 5.38E-04 | 1.03E-02 |
| HIST1H3J | 0.37 | 5.36E-04 | 1.03E-02 |
| FOXM1 | 0.41 | 5.41E-04 | 1.03E-02 |
| NUSAP1 | 0.37 | 5.40E-04 | 1.03E-02 |
| WDR6 | 0.75 | 5.43E-04 | 1.03E-02 |
| POLE | 0.61 | 5.47E-04 | 1.04E-02 |
| NCAPD2 | 0.58 | 5.53E-04 | 1.05E-02 |
| PAK7 | 0.40 | 5.59E-04 | 1.05E-02 |
| ALKBH7 | 0.55 | 5.61E-04 | 1.06E-02 |
| SLITRK5 | 0.22 | 5.63E-04 | 1.06E-02 |
| HMGN2 | 0.63 | 5.67E-04 | 1.06E-02 |
| ESPL1 | 0.38 | 5.72E-04 | 1.07E-02 |
| NUBP2 | 0.67 | 5.74E-04 | 1.07E-02 |
| EGFEM1P | 0.25 | 5.77E-04 | 1.08E-02 |
| TIMM10 | 0.59 | 5.88E-04 | 1.09E-02 |
| MIR3687 | 0.36 | 5.90E-04 | 1.09E-02 |
| RMRP | 0.52 | 5.92E-04 | 1.10E-02 |
| LOC148709 | 0.31 | 5.96E-04 | 1.10E-02 |
| DALRD3 | 0.67 | 6.01E-04 | 1.11E-02 |
| OST4 | 0.61 | 6.02E-04 | 1.11E-02 |
| SPAG5 | 0.41 | 6.07E-04 | 1.11E-02 |
| TUFM | 0.66 | 6.15E-04 | 1.13E-02 |
| KIAA0101 | 0.32 | 6.17E-04 | 1.13E-02 |
| WDR54 | 0.63 | 6.18E-04 | 1.13E-02 |
| FAHD2A | 0.75 | 6.23E-04 | 1.14E-02 |
| CDK1 | 0.35 | 6.30E-04 | 1.14E-02 |
| KLHL31 | 0.32 | 6.32E-04 | 1.14E-02 |
| ORC6 | 0.40 | 6.29E-04 | 1.14E-02 |
| TMEM147 | 0.68 | 6.28E-04 | 1.14E-02 |
| LOC284801 | 0.47 | 6.38E-04 | 1.15E-02 |
| LSM2 | 0.68 | 6.39E-04 | 1.15E-02 |
| ZNF428 | 0.69 | 6.38E-04 | 1.15E-02 |
| SULT1A1 | 0.44 | 6.44E-04 | 1.16E-02 |
| UROS | 0.73 | 6.47E-04 | 1.16E-02 |
| C14orf80 | 0.52 | 6.49E-04 | 1.16E-02 |
| WDR77 | 0.43 | 6.50E-04 | 1.16E-02 |
| PSMB6 | 0.67 | 6.65E-04 | 1.19E-02 |
| METTL17 | 0.73 | 6.66E-04 | 1.19E-02 |
| STOML2 | 0.68 | 6.70E-04 | 1.19E-02 |
| AURKAIP1 | 0.61 | 6.71E-04 | 1.19E-02 |
| APEH | 0.73 | 6.75E-04 | 1.20E-02 |
| CBX2 | 0.44 | 6.75E-04 | 1.20E-02 |
| LSM7 | 0.59 | 6.77E-04 | 1.20E-02 |
| PGLS | 0.62 | 6.76E-04 | 1.20E-02 |
| TMEM9 | 0.71 | 6.79E-04 | 1.20E-02 |
| NMRAL1 | 0.70 | 6.80E-04 | 1.20E-02 |
| TICRR | 0.41 | 6.89E-04 | 1.21E-02 |
| EPN3 | 0.47 | 6.90E-04 | 1.21E-02 |
| PRRT3 | 0.61 | 6.92E-04 | 1.21E-02 |
| CCNF | 0.51 | 6.99E-04 | 1.22E-02 |
| SIAH3 | 0.35 | 6.99E-04 | 1.22E-02 |
| TACC3 | 0.47 | 7.06E-04 | 1.23E-02 |
| C20orf96 | 0.63 | 7.09E-04 | 1.23E-02 |
| PIK3R2 | 0.65 | 7.15E-04 | 1.24E-02 |
| SEMA3G | 0.31 | 7.18E-04 | 1.24E-02 |
| CPSF4 | 0.66 | 7.26E-04 | 1.25E-02 |
| ECHS1 | 0.68 | 7.39E-04 | 1.27E-02 |
| MCM7 | 0.56 | 7.40E-04 | 1.27E-02 |
| C19orf57 | 0.54 | 7.44E-04 | 1.27E-02 |
| UBE2T | 0.38 | 7.43E-04 | 1.27E-02 |
| PAK6 | 0.50 | 7.52E-04 | 1.28E-02 |
| KLK6 | 0.31 | 7.55E-04 | 1.29E-02 |
| HMGB2 | 0.39 | 7.57E-04 | 1.29E-02 |
| SLIRP | 0.58 | 7.57E-04 | 1.29E-02 |
| HIST1H4L | 0.43 | 7.60E-04 | 1.29E-02 |
| COPS6 | 0.73 | 7.68E-04 | 1.30E-02 |
| FAM169A | 0.46 | 7.68E-04 | 1.30E-02 |
| TMEM178A | 0.43 | 7.72E-04 | 1.30E-02 |
| STAP2 | 0.58 | 7.75E-04 | 1.31E-02 |
| DEPDC1 | 0.27 | 7.78E-04 | 1.31E-02 |
| RANBP1 | 0.63 | 7.78E-04 | 1.31E-02 |
| MRPS34 | 0.58 | 7.84E-04 | 1.32E-02 |
| MAGED2 | 0.67 | 7.87E-04 | 1.32E-02 |
| RPUSD1 | 0.65 | 7.88E-04 | 1.32E-02 |
| EME1 | 0.44 | 7.91E-04 | 1.32E-02 |
| RNF187 | 0.75 | 7.96E-04 | 1.33E-02 |
| GTSE1 | 0.40 | 8.04E-04 | 1.33E-02 |
| CCDC24 | 0.54 | 8.21E-04 | 1.35E-02 |
| HMGN1 | 0.70 | 8.20E-04 | 1.35E-02 |
| KCNN1 | 0.43 | 8.26E-04 | 1.36E-02 |
| MTRNR2L1 | 0.37 | 8.27E-04 | 1.36E-02 |
| MARVELD3 | 0.52 | 8.30E-04 | 1.36E-02 |
| FAM195B | 0.71 | 8.41E-04 | 1.37E-02 |
| EIF2B4 | 0.81 | 8.43E-04 | 1.37E-02 |
| EMC10 | 0.68 | 8.45E-04 | 1.37E-02 |
| FAM136A | 0.70 | 8.50E-04 | 1.38E-02 |
| NLE1 | 0.70 | 8.52E-04 | 1.38E-02 |
| TSSC4 | 0.67 | 8.54E-04 | 1.38E-02 |
| APOA1BP | 0.62 | 8.55E-04 | 1.38E-02 |
| LOC400680 | 0.51 | 8.57E-04 | 1.38E-02 |
| CASKIN2 | 0.76 | 8.63E-04 | 1.38E-02 |
| MRPL12 | 0.61 | 8.66E-04 | 1.39E-02 |
| TOP2A | 0.32 | 8.70E-04 | 1.39E-02 |
| TMPO-AS1 | 0.46 | 8.71E-04 | 1.39E-02 |
| DNAJC9 | 0.63 | 8.82E-04 | 1.41E-02 |
| PSMD3 | 0.74 | 8.86E-04 | 1.41E-02 |
| PTPRN2 | 0.50 | 8.85E-04 | 1.41E-02 |
| HIST1H4E | 0.55 | 9.03E-04 | 1.43E-02 |
| LINC00617 | 0.39 | 9.03E-04 | 1.43E-02 |
| TMSB15A | 0.38 | 9.02E-04 | 1.43E-02 |
| NGFRAP1 | 0.65 | 9.06E-04 | 1.43E-02 |
| PRMT1 | 0.70 | 9.06E-04 | 1.43E-02 |
| MAD2L1 | 0.34 | 9.27E-04 | 1.46E-02 |
| PIP5KL1 | 0.50 | 9.28E-04 | 1.46E-02 |
| PRR15L | 0.39 | 9.29E-04 | 1.46E-02 |
| MLF1IP | 0.36 | 9.32E-04 | 1.46E-02 |
| VANGL1 | 0.70 | 9.32E-04 | 1.46E-02 |
| CDH24 | 0.55 | 9.37E-04 | 1.46E-02 |
| DPH2 | 0.73 | 9.49E-04 | 1.48E-02 |
| RPL18A | 0.62 | 9.50E-04 | 1.48E-02 |
| SLC27A5 | 0.51 | 9.52E-04 | 1.48E-02 |
| TIMM17B | 0.69 | 9.54E-04 | 1.48E-02 |
| KLHDC9 | 0.50 | 9.70E-04 | 1.50E-02 |
| FKBP10 | 0.69 | 9.72E-04 | 1.50E-02 |
| RNF26 | 0.70 | 1.01E-03 | 1.54E-02 |
| LINC00665 | 0.58 | 1.01E-03 | 1.54E-02 |
| TRPT1 | 0.65 | 1.01E-03 | 1.54E-02 |
| CCNO | 0.32 | 1.02E-03 | 1.55E-02 |
| RPS10 | 0.60 | 1.04E-03 | 1.57E-02 |
| KMO | 0.24 | 1.06E-03 | 1.60E-02 |
| DHPS | 0.76 | 1.06E-03 | 1.60E-02 |
| AZI1 | 0.64 | 1.08E-03 | 1.62E-02 |
| C16orf58 | 0.78 | 1.08E-03 | 1.62E-02 |
| FBXW9 | 0.63 | 1.08E-03 | 1.62E-02 |
| TONSL | 0.48 | 1.08E-03 | 1.62E-02 |
| WBSCR22 | 0.73 | 1.08E-03 | 1.62E-02 |
| PEX10 | 0.64 | 1.08E-03 | 1.62E-02 |
| TP53I13 | 0.70 | 1.09E-03 | 1.62E-02 |
| KLHDC3 | 0.69 | 1.09E-03 | 1.62E-02 |
| FSCN1 | 0.62 | 1.09E-03 | 1.63E-02 |
| AKR7A2 | 0.72 | 1.10E-03 | 1.63E-02 |
| CDH10 | 0.16 | 1.10E-03 | 1.63E-02 |
| NDUFA2 | 0.56 | 1.11E-03 | 1.64E-02 |
| USP5 | 0.72 | 1.12E-03 | 1.65E-02 |
| GREM2 | 0.25 | 1.13E-03 | 1.67E-02 |
| APH1A | 0.72 | 1.14E-03 | 1.68E-02 |
| DNAJC10 | 0.51 | 1.14E-03 | 1.68E-02 |
| RMI2 | 0.50 | 1.14E-03 | 1.68E-02 |
| MYCBP | 0.54 | 1.15E-03 | 1.68E-02 |
| RPS19 | 0.69 | 1.15E-03 | 1.68E-02 |
| PBK | 0.29 | 1.15E-03 | 1.68E-02 |
| YEATS4 | 0.63 | 1.15E-03 | 1.68E-02 |
| TMEM107 | 0.41 | 1.16E-03 | 1.68E-02 |
| FNDC5 | 0.45 | 1.17E-03 | 1.69E-02 |
| GGH | 0.38 | 1.17E-03 | 1.69E-02 |
| MRPS12 | 0.61 | 1.17E-03 | 1.69E-02 |
| HIST1H2BO | 0.32 | 1.18E-03 | 1.70E-02 |
| CDCA8 | 0.40 | 1.19E-03 | 1.71E-02 |
| CHID1 | 0.72 | 1.19E-03 | 1.71E-02 |
| RPP25 | 0.50 | 1.19E-03 | 1.71E-02 |
| KIAA1324 | 0.32 | 1.20E-03 | 1.72E-02 |
| C1orf35 | 0.69 | 1.20E-03 | 1.72E-02 |
| IPO4 | 0.70 | 1.21E-03 | 1.72E-02 |
| MND1 | 0.45 | 1.21E-03 | 1.72E-02 |
| DNLZ | 0.62 | 1.23E-03 | 1.73E-02 |
| ERAL1 | 0.77 | 1.23E-03 | 1.73E-02 |
| MRPL28 | 0.70 | 1.23E-03 | 1.73E-02 |
| CDCA7 | 0.38 | 1.23E-03 | 1.73E-02 |
| LAGE3 | 0.65 | 1.25E-03 | 1.75E-02 |
| RAB11B | 0.68 | 1.25E-03 | 1.75E-02 |
| TCF19 | 0.48 | 1.25E-03 | 1.75E-02 |
| HIST1H2BJ | 0.37 | 1.26E-03 | 1.76E-02 |
| C19orf24 | 0.60 | 1.27E-03 | 1.76E-02 |
| BUB1B | 0.37 | 1.27E-03 | 1.77E-02 |
| ALYREF | 0.64 | 1.28E-03 | 1.77E-02 |
| GGTA1P | 0.34 | 1.28E-03 | 1.77E-02 |
| INCENP | 0.57 | 1.28E-03 | 1.77E-02 |
| SLC39A7 | 0.68 | 1.29E-03 | 1.78E-02 |
| DUSP15 | 0.48 | 1.29E-03 | 1.78E-02 |
| RCOR2 | 0.51 | 1.29E-03 | 1.78E-02 |
| DDOST | 0.68 | 1.30E-03 | 1.78E-02 |
| TTK | 0.36 | 1.30E-03 | 1.78E-02 |
| CRABP2 | 0.48 | 1.32E-03 | 1.80E-02 |
| NCKIPSD | 0.74 | 1.34E-03 | 1.82E-02 |
| SNRPB | 0.69 | 1.34E-03 | 1.83E-02 |
| BAI1 | 0.24 | 1.34E-03 | 1.83E-02 |
| EXO1 | 0.34 | 1.35E-03 | 1.83E-02 |
| TUBA1B | 0.54 | 1.36E-03 | 1.84E-02 |
| CYC1 | 0.66 | 1.37E-03 | 1.86E-02 |
| TK1 | 0.44 | 1.38E-03 | 1.86E-02 |
| DUT | 0.63 | 1.39E-03 | 1.87E-02 |
| WDR34 | 0.57 | 1.39E-03 | 1.87E-02 |
| EXOC7 | 0.74 | 1.41E-03 | 1.89E-02 |
| HNRNPL | 0.76 | 1.42E-03 | 1.90E-02 |
| NDUFC2 | 0.57 | 1.43E-03 | 1.91E-02 |
| ADPRHL1 | 0.66 | 1.45E-03 | 1.92E-02 |
| GINS4 | 0.49 | 1.45E-03 | 1.93E-02 |
| ATP8B3 | 0.46 | 1.45E-03 | 1.93E-02 |
| AUNIP | 0.48 | 1.46E-03 | 1.94E-02 |
| PCNA | 0.50 | 1.46E-03 | 1.94E-02 |
| PIGT | 0.73 | 1.47E-03 | 1.94E-02 |
| HAGH | 0.68 | 1.48E-03 | 1.95E-02 |
| HCFC1R1 | 0.51 | 1.48E-03 | 1.95E-02 |
| MRPS26 | 0.65 | 1.49E-03 | 1.95E-02 |
| SRP14 | 0.68 | 1.49E-03 | 1.95E-02 |
| FBL | 0.67 | 1.50E-03 | 1.96E-02 |
| PCSK1N | 0.40 | 1.50E-03 | 1.96E-02 |
| CUEDC2 | 0.72 | 1.51E-03 | 1.97E-02 |
| YIF1A | 0.68 | 1.51E-03 | 1.97E-02 |
| HIST1H3I | 0.49 | 1.52E-03 | 1.98E-02 |
| TPX2 | 0.39 | 1.54E-03 | 1.99E-02 |
| UBXN6 | 0.71 | 1.56E-03 | 2.01E-02 |
| TMEM97 | 0.50 | 1.57E-03 | 2.01E-02 |
| MYBL2 | 0.37 | 1.58E-03 | 2.03E-02 |
| GLI1 | 0.52 | 1.59E-03 | 2.03E-02 |
| PKP2 | 0.56 | 1.59E-03 | 2.03E-02 |
| TRIP6 | 0.73 | 1.60E-03 | 2.04E-02 |
| MACROD1 | 0.42 | 1.60E-03 | 2.05E-02 |
| DCXR | 0.57 | 1.61E-03 | 2.05E-02 |
| TRIM9 | 0.65 | 1.62E-03 | 2.06E-02 |
| GJA4 | 0.42 | 1.63E-03 | 2.07E-02 |
| ANAPC15 | 0.57 | 1.65E-03 | 2.08E-02 |
| MTFP1 | 0.54 | 1.66E-03 | 2.09E-02 |
| E2F1 | 0.40 | 1.67E-03 | 2.10E-02 |
| HOXA10 | 0.61 | 1.72E-03 | 2.14E-02 |
| TOR3A | 0.75 | 1.72E-03 | 2.14E-02 |
| ZNF511 | 0.67 | 1.72E-03 | 2.14E-02 |
| AKR1A1 | 0.74 | 1.75E-03 | 2.16E-02 |
| PLK1 | 0.45 | 1.75E-03 | 2.16E-02 |
| RPS9 | 0.68 | 1.79E-03 | 2.20E-02 |
| SORD | 0.58 | 1.79E-03 | 2.20E-02 |
| KRT8 | 0.50 | 1.80E-03 | 2.22E-02 |
| THOC3 | 0.64 | 1.80E-03 | 2.22E-02 |
| DSN1 | 0.58 | 1.81E-03 | 2.22E-02 |
| PPAP2C | 0.59 | 1.81E-03 | 2.22E-02 |
| NECAB3 | 0.59 | 1.82E-03 | 2.23E-02 |
| AIF1L | 0.46 | 1.83E-03 | 2.24E-02 |
| PTCH2 | 0.45 | 1.84E-03 | 2.24E-02 |
| CPM | 0.34 | 1.85E-03 | 2.25E-02 |
| SLC29A1 | 0.68 | 1.85E-03 | 2.25E-02 |
| RFNG | 0.67 | 1.89E-03 | 2.28E-02 |
| CNIH2 | 0.47 | 1.89E-03 | 2.29E-02 |
| OBSL1 | 0.73 | 1.90E-03 | 2.29E-02 |
| CKS2 | 0.43 | 1.91E-03 | 2.30E-02 |
| LZTS1 | 0.55 | 1.91E-03 | 2.30E-02 |
| SMAD9 | 0.47 | 1.91E-03 | 2.30E-02 |
| NENF | 0.63 | 1.93E-03 | 2.32E-02 |
| PUSL1 | 0.65 | 1.96E-03 | 2.35E-02 |
| SAMM50 | 0.72 | 1.97E-03 | 2.36E-02 |
| SF3B5 | 0.69 | 1.97E-03 | 2.36E-02 |
| SNAPIN | 0.59 | 1.98E-03 | 2.36E-02 |
| TMEM160 | 0.48 | 1.98E-03 | 2.36E-02 |
| KIAA1967 | 0.78 | 2.02E-03 | 2.39E-02 |
| TARBP2 | 0.67 | 2.03E-03 | 2.40E-02 |
| TBCCD1 | 0.84 | 2.04E-03 | 2.40E-02 |
| SARS2 | 0.70 | 2.05E-03 | 2.41E-02 |
| PRR15 | 0.45 | 2.05E-03 | 2.41E-02 |
| ANAPC11 | 0.68 | 2.07E-03 | 2.43E-02 |
| MRPL17 | 0.74 | 2.08E-03 | 2.44E-02 |
| OTUB1 | 0.79 | 2.08E-03 | 2.44E-02 |
| TM7SF2 | 0.53 | 2.08E-03 | 2.44E-02 |
| RFC5 | 0.72 | 2.09E-03 | 2.44E-02 |
| ALDH3B2 | 0.34 | 2.09E-03 | 2.44E-02 |
| CDC6 | 0.38 | 2.10E-03 | 2.45E-02 |
| DLGAP5 | 0.37 | 2.10E-03 | 2.45E-02 |
| MCM4 | 0.46 | 2.10E-03 | 2.45E-02 |
| B3GALT6 | 0.69 | 2.11E-03 | 2.45E-02 |
| COL1A1 | 0.55 | 2.11E-03 | 2.45E-02 |
| PXMP2 | 0.58 | 2.12E-03 | 2.47E-02 |
| CENPH | 0.53 | 2.13E-03 | 2.47E-02 |
| SHMT1 | 0.60 | 2.18E-03 | 2.52E-02 |
| KIF15 | 0.39 | 2.19E-03 | 2.52E-02 |
| EMC4 | 0.64 | 2.20E-03 | 2.53E-02 |
| OLFM1 | 0.37 | 2.21E-03 | 2.53E-02 |
| PRADC1 | 0.59 | 2.22E-03 | 2.54E-02 |
| COL1A2 | 0.53 | 2.24E-03 | 2.55E-02 |
| IFT20 | 0.68 | 2.24E-03 | 2.55E-02 |
| PTPRF | 0.64 | 2.25E-03 | 2.56E-02 |
| SEMA3D | 0.26 | 2.25E-03 | 2.56E-02 |
| NUDT8 | 0.48 | 2.26E-03 | 2.57E-02 |
| KIF18B | 0.40 | 2.27E-03 | 2.58E-02 |
| PCDH19 | 0.50 | 2.28E-03 | 2.58E-02 |
| SYNDIG1 | 0.38 | 2.28E-03 | 2.58E-02 |
| GCN1L1 | 0.81 | 2.29E-03 | 2.60E-02 |
| VWA5B2 | 0.38 | 2.30E-03 | 2.60E-02 |
| AEN | 0.53 | 2.31E-03 | 2.61E-02 |
| KIAA1467 | 0.44 | 2.31E-03 | 2.61E-02 |
| NCAPH2 | 0.67 | 2.31E-03 | 2.61E-02 |
| SEC14L1 | 0.42 | 2.33E-03 | 2.62E-02 |
| IQCC | 0.65 | 2.34E-03 | 2.62E-02 |
| RRP7A | 0.67 | 2.35E-03 | 2.63E-02 |
| NCAPH | 0.42 | 2.35E-03 | 2.63E-02 |
| ZNF205 | 0.71 | 2.38E-03 | 2.66E-02 |
| CENPN | 0.50 | 2.40E-03 | 2.67E-02 |
| E4F1 | 0.77 | 2.40E-03 | 2.67E-02 |
| CDK20 | 0.67 | 2.41E-03 | 2.68E-02 |
| RPS2 | 0.68 | 2.42E-03 | 2.68E-02 |
| CCDC74A | 0.58 | 2.43E-03 | 2.68E-02 |
| FKBP4 | 0.68 | 2.42E-03 | 2.68E-02 |
| HIST1H2AC | 0.51 | 2.42E-03 | 2.68E-02 |
| POC1B | 0.59 | 2.43E-03 | 2.68E-02 |
| TTLL7 | 0.63 | 2.43E-03 | 2.68E-02 |
| NME3 | 0.55 | 2.45E-03 | 2.70E-02 |
| SCAMP5 | 0.48 | 2.45E-03 | 2.70E-02 |
| RPL35 | 0.55 | 2.47E-03 | 2.72E-02 |
| PGRMC1 | 0.61 | 2.49E-03 | 2.74E-02 |
| MRPL52 | 0.78 | 2.50E-03 | 2.74E-02 |
| TRIP13 | 0.47 | 2.51E-03 | 2.75E-02 |
| LSMD1 | 0.61 | 2.54E-03 | 2.77E-02 |
| MEA1 | 0.63 | 2.54E-03 | 2.77E-02 |
| SKA3 | 0.40 | 2.54E-03 | 2.77E-02 |
| TYRO3 | 0.67 | 2.54E-03 | 2.77E-02 |
| UCK1 | 0.71 | 2.54E-03 | 2.77E-02 |
| ACADL | 0.44 | 2.57E-03 | 2.79E-02 |
| FOXRED2 | 0.66 | 2.57E-03 | 2.79E-02 |
| FAM127A | 0.73 | 2.58E-03 | 2.80E-02 |
| H2AFZ | 0.48 | 2.58E-03 | 2.80E-02 |
| RPL41 | 0.70 | 2.59E-03 | 2.81E-02 |
| ATP5G3 | 0.59 | 2.63E-03 | 2.84E-02 |
| FBXO5 | 0.51 | 2.65E-03 | 2.84E-02 |
| NIPSNAP1 | 0.73 | 2.65E-03 | 2.84E-02 |
| PDE6B | 0.63 | 2.64E-03 | 2.84E-02 |
| MNF1 | 0.69 | 2.65E-03 | 2.85E-02 |
| GCLC | 0.49 | 2.66E-03 | 2.85E-02 |
| RNASEH2C | 0.63 | 2.66E-03 | 2.85E-02 |
| ADAMTS6 | 0.38 | 2.68E-03 | 2.86E-02 |
| ERCC1 | 0.75 | 2.68E-03 | 2.86E-02 |
| SSRP1 | 0.73 | 2.68E-03 | 2.86E-02 |
| INTS5 | 0.74 | 2.72E-03 | 2.89E-02 |
| CLSPN | 0.41 | 2.72E-03 | 2.89E-02 |
| BCS1L | 0.71 | 2.73E-03 | 2.89E-02 |
| ECI2 | 0.43 | 2.73E-03 | 2.90E-02 |
| ILVBL | 0.67 | 2.73E-03 | 2.90E-02 |
| OSGEP | 0.78 | 2.76E-03 | 2.92E-02 |
| MIS18A | 0.60 | 2.82E-03 | 2.97E-02 |
| SMCR7 | 0.65 | 2.82E-03 | 2.97E-02 |
| CIT | 0.47 | 2.84E-03 | 2.98E-02 |
| ELP5 | 0.78 | 2.84E-03 | 2.98E-02 |
| FEN1 | 0.50 | 2.84E-03 | 2.98E-02 |
| NDUFS3 | 0.73 | 2.85E-03 | 2.98E-02 |
| SLC9A4 | 0.33 | 2.85E-03 | 2.98E-02 |
| SCRN2 | 0.64 | 2.85E-03 | 2.98E-02 |
| SKA1 | 0.38 | 2.86E-03 | 2.99E-02 |
| NCLN | 0.75 | 2.87E-03 | 3.00E-02 |
| PHPT1 | 0.67 | 2.88E-03 | 3.00E-02 |
| SGOL1 | 0.46 | 2.89E-03 | 3.01E-02 |
| LIN7B | 0.56 | 2.90E-03 | 3.01E-02 |
| SMYD5 | 0.79 | 2.90E-03 | 3.01E-02 |
| DDAH2 | 0.65 | 2.93E-03 | 3.02E-02 |
| PSMB5 | 0.71 | 2.93E-03 | 3.02E-02 |
| TERC | 0.46 | 2.93E-03 | 3.02E-02 |
| DNA2 | 0.55 | 2.95E-03 | 3.03E-02 |
| ESCO2 | 0.35 | 2.95E-03 | 3.03E-02 |
| KIF4A | 0.41 | 2.94E-03 | 3.03E-02 |
| TRIAP1 | 0.65 | 2.95E-03 | 3.03E-02 |
| PARP1 | 0.75 | 2.95E-03 | 3.04E-02 |
| LLGL1 | 0.69 | 2.99E-03 | 3.06E-02 |
| PRR11 | 0.52 | 3.01E-03 | 3.07E-02 |
| TRAPPC6A | 0.54 | 3.01E-03 | 3.08E-02 |
| IRF2BP1 | 0.71 | 3.02E-03 | 3.09E-02 |
| TEC | 0.50 | 3.04E-03 | 3.10E-02 |
| C4orf48 | 0.57 | 3.06E-03 | 3.11E-02 |
| CYP2J2 | 0.28 | 3.09E-03 | 3.12E-02 |
| FAM58A | 0.70 | 3.08E-03 | 3.12E-02 |
| MPI | 0.74 | 3.09E-03 | 3.12E-02 |
| RTKN2 | 0.47 | 3.08E-03 | 3.12E-02 |
| HIST1H2BM | 0.41 | 3.09E-03 | 3.12E-02 |
| CDH4 | 0.31 | 3.09E-03 | 3.13E-02 |
| FANCA | 0.49 | 3.10E-03 | 3.13E-02 |
| PCYOX1L | 0.62 | 3.14E-03 | 3.15E-02 |
| LOC440335 | 0.47 | 3.14E-03 | 3.16E-02 |
| SEPHS1 | 0.78 | 3.15E-03 | 3.16E-02 |
| MAZ | 0.64 | 3.18E-03 | 3.18E-02 |
| SERPINA4 | 0.23 | 3.19E-03 | 3.19E-02 |
| ANLN | 0.38 | 3.22E-03 | 3.21E-02 |
| POP7 | 0.65 | 3.22E-03 | 3.21E-02 |
| CDKN3 | 0.43 | 3.22E-03 | 3.21E-02 |
| KIAA2013 | 0.76 | 3.22E-03 | 3.21E-02 |
| PDCD11 | 0.84 | 3.23E-03 | 3.22E-02 |
| RNF215 | 0.72 | 3.25E-03 | 3.23E-02 |
| C21orf88 | 0.57 | 3.26E-03 | 3.24E-02 |
| SAC3D1 | 0.41 | 3.26E-03 | 3.24E-02 |
| NKAIN3 | 0.44 | 3.28E-03 | 3.25E-02 |
| ASNA1 | 0.68 | 3.28E-03 | 3.26E-02 |
| HIST2H2BE | 0.48 | 3.29E-03 | 3.26E-02 |
| ADRM1 | 0.74 | 3.30E-03 | 3.26E-02 |
| CKAP2 | 0.49 | 3.31E-03 | 3.27E-02 |
| AGPAT2 | 0.61 | 3.32E-03 | 3.27E-02 |
| CEP55 | 0.41 | 3.32E-03 | 3.27E-02 |
| PLK4 | 0.43 | 3.32E-03 | 3.27E-02 |
| PMF1 | 0.61 | 3.35E-03 | 3.29E-02 |
| TRIM45 | 0.63 | 3.35E-03 | 3.29E-02 |
| REC8 | 0.53 | 3.36E-03 | 3.30E-02 |
| FAM71E1 | 0.47 | 3.37E-03 | 3.30E-02 |
| OXLD1 | 0.59 | 3.40E-03 | 3.32E-02 |
| PTPMT1 | 0.68 | 3.42E-03 | 3.34E-02 |
| DACT2 | 0.40 | 3.43E-03 | 3.34E-02 |
| SLC7A4 | 0.30 | 3.43E-03 | 3.34E-02 |
| SLITRK6 | 0.24 | 3.43E-03 | 3.34E-02 |
| CHAF1A | 0.54 | 3.46E-03 | 3.37E-02 |
| C1orf233 | 0.47 | 3.47E-03 | 3.37E-02 |
| FGFR4 | 0.47 | 3.47E-03 | 3.37E-02 |
| PNKP | 0.71 | 3.48E-03 | 3.38E-02 |
| ATP5I | 0.63 | 3.51E-03 | 3.40E-02 |
| GSTP1 | 0.65 | 3.53E-03 | 3.42E-02 |
| SIGMAR1 | 0.69 | 3.53E-03 | 3.42E-02 |
| SLC25A15 | 0.63 | 3.53E-03 | 3.42E-02 |
| CDC45 | 0.41 | 3.59E-03 | 3.46E-02 |
| GTF3C4 | 0.78 | 3.60E-03 | 3.46E-02 |
| NOP56 | 0.78 | 3.63E-03 | 3.49E-02 |
| POLD2 | 0.68 | 3.63E-03 | 3.49E-02 |
| NAT8L | 0.38 | 3.64E-03 | 3.49E-02 |
| NPRL2 | 0.73 | 3.64E-03 | 3.50E-02 |
| C9orf123 | 0.67 | 3.65E-03 | 3.50E-02 |
| GMPPB | 0.67 | 3.65E-03 | 3.50E-02 |
| REEP4 | 0.61 | 3.66E-03 | 3.50E-02 |
| ZDHHC4 | 0.78 | 3.68E-03 | 3.52E-02 |
| NDUFA8 | 0.69 | 3.69E-03 | 3.52E-02 |
| FBXW12 | 0.29 | 3.70E-03 | 3.52E-02 |
| LAMA1 | 0.43 | 3.70E-03 | 3.52E-02 |
| NOL6 | 0.77 | 3.73E-03 | 3.54E-02 |
| TFPT | 0.63 | 3.77E-03 | 3.58E-02 |
| KHSRP | 0.80 | 3.78E-03 | 3.58E-02 |
| FANCB | 0.51 | 3.80E-03 | 3.60E-02 |
| EFCAB4A | 0.45 | 3.83E-03 | 3.61E-02 |
| MRPL38 | 0.75 | 3.82E-03 | 3.61E-02 |
| PHB2 | 0.65 | 3.82E-03 | 3.61E-02 |
| WDR62 | 0.48 | 3.83E-03 | 3.61E-02 |
| ZNF578 | 0.65 | 3.86E-03 | 3.63E-02 |
| SSNA1 | 0.62 | 3.87E-03 | 3.64E-02 |
| ZNF282 | 0.69 | 3.87E-03 | 3.64E-02 |
| CREB3L1 | 0.56 | 3.88E-03 | 3.64E-02 |
| DOLK | 0.72 | 3.87E-03 | 3.64E-02 |
| HRC | 0.31 | 3.88E-03 | 3.64E-02 |
| SLC25A5 | 0.62 | 3.88E-03 | 3.64E-02 |
| UBAC1 | 0.80 | 3.88E-03 | 3.64E-02 |
| FBXL15 | 0.62 | 3.91E-03 | 3.65E-02 |
| PATZ1 | 0.74 | 3.91E-03 | 3.65E-02 |
| PARP10 | 0.73 | 3.92E-03 | 3.65E-02 |
| SNRNP25 | 0.67 | 3.94E-03 | 3.66E-02 |
| ALDH1B1 | 0.54 | 3.95E-03 | 3.67E-02 |
| TMEM219 | 0.71 | 3.96E-03 | 3.67E-02 |
| COQ5 | 0.79 | 3.96E-03 | 3.67E-02 |
| CD276 | 0.77 | 3.97E-03 | 3.68E-02 |
| E2F7 | 0.39 | 3.98E-03 | 3.68E-02 |
| SLC25A39 | 0.68 | 3.98E-03 | 3.68E-02 |
| SPC25 | 0.38 | 3.99E-03 | 3.68E-02 |
| IGSF8 | 0.68 | 4.01E-03 | 3.69E-02 |
| ADAMTS18 | 0.34 | 4.03E-03 | 3.71E-02 |
| PARK7 | 0.72 | 4.03E-03 | 3.71E-02 |
| CENPE | 0.38 | 4.05E-03 | 3.71E-02 |
| MBD3 | 0.66 | 4.07E-03 | 3.73E-02 |
| C8orf33 | 0.72 | 4.13E-03 | 3.77E-02 |
| NOL12 | 0.76 | 4.13E-03 | 3.77E-02 |
| TP53 | 0.75 | 4.13E-03 | 3.77E-02 |
| CCDC106 | 0.57 | 4.15E-03 | 3.78E-02 |
| RACGAP1 | 0.49 | 4.14E-03 | 3.78E-02 |
| RIC8A | 0.75 | 4.14E-03 | 3.78E-02 |
| RRP1 | 0.80 | 4.14E-03 | 3.78E-02 |
| TXN2 | 0.76 | 4.17E-03 | 3.79E-02 |
| ATP5O | 0.72 | 4.20E-03 | 3.80E-02 |
| ATRAID | 0.65 | 4.19E-03 | 3.80E-02 |
| HDGFRP2 | 0.71 | 4.20E-03 | 3.80E-02 |
| LPPR3 | 0.32 | 4.22E-03 | 3.81E-02 |
| MFSD10 | 0.77 | 4.23E-03 | 3.82E-02 |
| DCTN3 | 0.70 | 4.23E-03 | 3.82E-02 |
| C19orf53 | 0.61 | 4.25E-03 | 3.83E-02 |
| POLR2E | 0.72 | 4.26E-03 | 3.83E-02 |
| NTPCR | 0.73 | 4.26E-03 | 3.84E-02 |
| RCCD1 | 0.70 | 4.27E-03 | 3.84E-02 |
| PMVK | 0.66 | 4.29E-03 | 3.84E-02 |
| UBXN11 | 0.66 | 4.31E-03 | 3.86E-02 |
| PANK1 | 0.64 | 4.34E-03 | 3.87E-02 |
| UHRF1 | 0.49 | 4.33E-03 | 3.87E-02 |
| DPYSL4 | 0.56 | 4.38E-03 | 3.91E-02 |
| CHST14 | 0.73 | 4.40E-03 | 3.91E-02 |
| LOC202781 | 0.63 | 4.40E-03 | 3.91E-02 |
| SLC15A2 | 0.28 | 4.41E-03 | 3.91E-02 |
| SLC29A2 | 0.55 | 4.40E-03 | 3.91E-02 |
| VKORC1 | 0.68 | 4.41E-03 | 3.91E-02 |
| C3orf67 | 0.58 | 4.47E-03 | 3.96E-02 |
| EIF4EBP1 | 0.63 | 4.48E-03 | 3.96E-02 |
| RHBDD3 | 0.59 | 4.53E-03 | 4.00E-02 |
| CKAP2L | 0.43 | 4.54E-03 | 4.01E-02 |
| COX4I1 | 0.69 | 4.54E-03 | 4.01E-02 |
| KIF24 | 0.55 | 4.54E-03 | 4.01E-02 |
| NME2 | 0.63 | 4.56E-03 | 4.01E-02 |
| MKI67 | 0.42 | 4.59E-03 | 4.04E-02 |
| VARS2 | 0.72 | 4.61E-03 | 4.05E-02 |
| UQCR10 | 0.69 | 4.62E-03 | 4.06E-02 |
| PODXL2 | 0.47 | 4.63E-03 | 4.06E-02 |
| FASTK | 0.71 | 4.63E-03 | 4.06E-02 |
| HIST2H2AB | 0.48 | 4.70E-03 | 4.11E-02 |
| EXOSC5 | 0.64 | 4.72E-03 | 4.12E-02 |
| CDK5 | 0.63 | 4.73E-03 | 4.12E-02 |
| PAFAH1B3 | 0.57 | 4.73E-03 | 4.12E-02 |
| PYCRL | 0.61 | 4.78E-03 | 4.16E-02 |
| DPP3 | 0.70 | 4.78E-03 | 4.16E-02 |
| POLA2 | 0.64 | 4.79E-03 | 4.16E-02 |
| POLR1C | 0.74 | 4.81E-03 | 4.17E-02 |
| DANCR | 0.50 | 4.85E-03 | 4.20E-02 |
| RRM2 | 0.37 | 4.88E-03 | 4.21E-02 |
| MCAT | 0.71 | 4.88E-03 | 4.21E-02 |
| NDC80 | 0.37 | 4.91E-03 | 4.23E-02 |
| RAD51AP1 | 0.43 | 4.92E-03 | 4.23E-02 |
| DDX12P | 0.52 | 4.92E-03 | 4.23E-02 |
| LEPRE1 | 0.69 | 4.93E-03 | 4.23E-02 |
| LINC00669 | 0.34 | 4.93E-03 | 4.23E-02 |
| RPPH1 | 0.56 | 4.95E-03 | 4.24E-02 |
| C19orf70 | 0.63 | 4.96E-03 | 4.25E-02 |
| UBE2S | 0.59 | 4.99E-03 | 4.26E-02 |
| GPC1 | 0.66 | 4.99E-03 | 4.26E-02 |
| CEP85 | 0.65 | 5.01E-03 | 4.27E-02 |
| RPS19BP1 | 0.72 | 5.01E-03 | 4.27E-02 |
| NPR2 | 0.70 | 5.04E-03 | 4.28E-02 |
| TIMM8A | 0.74 | 5.05E-03 | 4.29E-02 |
| FKBP9 | 0.80 | 5.08E-03 | 4.30E-02 |
| H1FX | 0.67 | 5.08E-03 | 4.30E-02 |
| PIDD | 0.66 | 5.11E-03 | 4.32E-02 |
| MCM10 | 0.42 | 5.12E-03 | 4.32E-02 |
| ZNF589 | 0.54 | 5.15E-03 | 4.34E-02 |
| HAGHL | 0.55 | 5.18E-03 | 4.34E-02 |
| NAT14 | 0.67 | 5.18E-03 | 4.34E-02 |
| SLC43A1 | 0.56 | 5.18E-03 | 4.34E-02 |
| TRAPPC5 | 0.62 | 5.17E-03 | 4.34E-02 |
| C8orf40 | 0.69 | 5.21E-03 | 4.35E-02 |
| SLC13A3 | 0.64 | 5.21E-03 | 4.35E-02 |
| TUBG1 | 0.69 | 5.22E-03 | 4.36E-02 |
| DHCR7 | 0.61 | 5.23E-03 | 4.37E-02 |
| WEE1 | 0.59 | 5.29E-03 | 4.40E-02 |
| TTI2 | 0.78 | 5.30E-03 | 4.41E-02 |
| NTHL1 | 0.69 | 5.31E-03 | 4.41E-02 |
| CHPF | 0.66 | 5.36E-03 | 4.45E-02 |
| GNPDA1 | 0.76 | 5.36E-03 | 4.45E-02 |
| LOC93622 | 0.70 | 5.37E-03 | 4.45E-02 |
| GGCT | 0.57 | 5.38E-03 | 4.45E-02 |
| RSG1 | 0.63 | 5.38E-03 | 4.45E-02 |
| POLR2C | 0.74 | 5.40E-03 | 4.46E-02 |
| MLST8 | 0.66 | 5.42E-03 | 4.47E-02 |
| NTN3 | 0.53 | 5.42E-03 | 4.47E-02 |
| RBBP7 | 0.68 | 5.42E-03 | 4.47E-02 |
| ZNHIT2 | 0.59 | 5.42E-03 | 4.47E-02 |
| FBF1 | 0.63 | 5.44E-03 | 4.48E-02 |
| ISG15 | 0.57 | 5.44E-03 | 4.48E-02 |
| SGOL2 | 0.49 | 5.44E-03 | 4.48E-02 |
| TESK1 | 0.67 | 5.43E-03 | 4.48E-02 |
| ARHGAP11A | 0.45 | 5.46E-03 | 4.48E-02 |
| PDZD11 | 0.72 | 5.46E-03 | 4.48E-02 |
| SMO | 0.72 | 5.45E-03 | 4.48E-02 |
| TUBB | 0.67 | 5.46E-03 | 4.49E-02 |
| GORASP1 | 0.75 | 5.49E-03 | 4.50E-02 |
| STUB1 | 0.64 | 5.49E-03 | 4.50E-02 |
| ISYNA1 | 0.57 | 5.49E-03 | 4.50E-02 |
| U2AF2 | 0.79 | 5.51E-03 | 4.51E-02 |
| LSM4 | 0.43 | 5.52E-03 | 4.52E-02 |
| E2F8 | 0.43 | 5.54E-03 | 4.53E-02 |
| C11orf48 | 0.72 | 5.57E-03 | 4.54E-02 |
| KLHL35 | 0.55 | 5.57E-03 | 4.54E-02 |
| HAUS4 | 0.75 | 5.60E-03 | 4.56E-02 |
| CKS1B | 0.58 | 5.64E-03 | 4.59E-02 |
| COX6C | 0.69 | 5.65E-03 | 4.59E-02 |
| NDUFA4 | 0.65 | 5.65E-03 | 4.59E-02 |
| RPLP2 | 0.76 | 5.67E-03 | 4.59E-02 |
| LOC81691 | 0.51 | 5.70E-03 | 4.61E-02 |
| CSRP2BP | 0.76 | 5.73E-03 | 4.62E-02 |
| PTGES2 | 0.71 | 5.74E-03 | 4.62E-02 |
| RPS5 | 0.64 | 5.74E-03 | 4.62E-02 |
| DPP10 | 0.28 | 5.76E-03 | 4.64E-02 |
| SLC25A11 | 0.73 | 5.77E-03 | 4.64E-02 |
| CCNB2 | 0.54 | 5.80E-03 | 4.65E-02 |
| GINS1 | 0.50 | 5.83E-03 | 4.67E-02 |
| OGDHL | 0.49 | 5.84E-03 | 4.68E-02 |
| UCHL1 | 0.42 | 5.85E-03 | 4.68E-02 |
| MCM5 | 0.64 | 5.88E-03 | 4.70E-02 |
| ENKD1 | 0.69 | 5.89E-03 | 4.71E-02 |
| TOR2A | 0.67 | 5.90E-03 | 4.71E-02 |
| SDCCAG3 | 0.81 | 5.94E-03 | 4.74E-02 |
| CHCHD1 | 0.71 | 5.96E-03 | 4.74E-02 |
| FTSJ1 | 0.76 | 5.96E-03 | 4.74E-02 |
| MEST | 0.47 | 5.96E-03 | 4.74E-02 |
| RFC2 | 0.73 | 5.96E-03 | 4.74E-02 |
| SLC39A3 | 0.73 | 5.97E-03 | 4.74E-02 |
| SH3RF2 | 0.46 | 5.98E-03 | 4.75E-02 |
| HIST1H2AL | 0.43 | 6.01E-03 | 4.76E-02 |
| CDC25C | 0.41 | 6.01E-03 | 4.76E-02 |
| ATP5F1 | 0.68 | 6.02E-03 | 4.76E-02 |
| GFER | 0.70 | 6.02E-03 | 4.76E-02 |
| FANCD2 | 0.50 | 6.04E-03 | 4.77E-02 |
| KIF11 | 0.43 | 6.06E-03 | 4.78E-02 |
| TSR3 | 0.67 | 6.10E-03 | 4.81E-02 |
| TMEM208 | 0.70 | 6.11E-03 | 4.81E-02 |
| LDHB | 0.66 | 6.17E-03 | 4.85E-02 |
| CELSR3 | 0.58 | 6.22E-03 | 4.88E-02 |
| IGSF9 | 0.49 | 6.23E-03 | 4.88E-02 |
| ENTPD2 | 0.44 | 6.25E-03 | 4.89E-02 |
| ROGDI | 0.54 | 6.28E-03 | 4.91E-02 |
| C9orf16 | 0.64 | 6.31E-03 | 4.92E-02 |
| CA11 | 0.49 | 6.33E-03 | 4.92E-02 |
| CCDC85B | 0.73 | 6.32E-03 | 4.92E-02 |
| MRPL4 | 0.73 | 6.34E-03 | 4.93E-02 |
| GLRX5 | 0.69 | 6.36E-03 | 4.94E-02 |
| SPATA20 | 0.72 | 6.42E-03 | 4.97E-02 |
| YARS2 | 0.78 | 6.41E-03 | 4.97E-02 |
| PNMAL1 | 0.57 | 6.44E-03 | 4.99E-02 |
