## Supplemental Table 1-10 for "miRNA/mRNA analysis of increased TGF-β pathways drive epithelial-mesenchymal transition and regulatory T cell differentiation": Supp Table 5.docx

| **Supp Table 5. Ingenuity canonical pathways enriched by significantly upregulated mRNAs in Endo+ compared to Endo- women with FDR<0.05 (Top 100)** | | | |
| --- | --- | --- | --- |
| **Ingenuity Canonical Pathways** | **FDR** | **Ratio** | **Molecules** |
| IL-10 Signaling | 2.51E-15 | 0.38 | CCR1,CCR5,CD14,FCGR2A,FCGR2C,IL10RA,IL10RB,IL18RAP,IL1A,IL1B,IL1R1,IL1R2,IL1RAP,IL1RN,IL4R,JAK1,MAP4K4,MAPK14,NFKB1,NFKB2,NFKBIA,NFKBID,REL,SOCS3,STAT3,TNF,TRAF6 |
| Role of Macrophages, Fibroblasts and Endothelial Cells in Rheumatoid Arthritis | 2.51E-15 | 0.18 | C5AR1,CCL2,CEBPB,CEBPD,CREBBP,CSF1,CXCL8,GNAQ,ICAM1,IL15,IL16,IL18R1,IL18RAP,IL1A,IL1B,IL1R1,IL1R2,IL1RAP,IL1RN,IL7,IRAK2,IRAK3,LTB,MAPK14,NFAT5,NFATC1,NFKB1,NFKBIA,NFKBID,PIK3CB,PIK3CG,PIK3R5,PLCL1,PPP3CC,PRKCA,PRKCB,PRKCH,RALA,RAP1A,RAP1B,RIPK1,SOCS3,STAT3,TCF7L2,TLR10,TLR2,TLR4,TLR8,TNF,TNFRSF1A,TNFRSF1B,TNFSF13B,TRAF1,TRAF3,TRAF6,VCAM1 |
| Th1 and Th2 Activation Pathway | 1.26E-14 | 0.23 | ACVR1C,CCR1,CCR5,CD28,CD3D,CD3G,CD40,CD40LG,CD80,CD86,GRB2,HAVCR2,ICAM1,ICOS,IFNAR1,IL10RA,IL10RB,IL12RB2,IL18R1,IL2RA,IL2RG,IL4R,IRF1,JAK1,JAK3,NFATC1,NFKB1,PIK3CB,PIK3CG,PIK3R5,PSEN1,RUNX3,SOCS3,SPI1,STAT3,STAT4,STAT5B,TGFBR2,VAV1 |
| PPARα/RXRα Activation | 2.51E-14 | 0.22 | ACVR1C,BCL3,CLOCK,CREBBP,CYP2C18,CYP2C19,CYP2C9,EP300,GK,GNA15,GNAQ,GRB2,IL18RAP,IL1B,IL1R1,IL1R2,IL1RAP,MAP4K4,MAPK14,NCOA3,NFKB1,NFKB2,NFKBIA,NFKBID,PLCL1,PPARGC1A,PRKAA1,PRKCA,PRKCB,RALA,RAP1A,RAP1B,REL,SMAD3,SOS1,SOS2,STAT5B,TGFB2,TGFBR2,TRAF6 |
| IL-6 Signaling | 5.01E-14 | 0.26 | CD14,CEBPB,CXCL8,GRB2,IL18RAP,IL1A,IL1B,IL1R1,IL1R2,IL1RAP,IL1RN,MAP4K4,MAPK14,NFKB1,NFKB2,NFKBIA,NFKBID,PIK3CB,PIK3CG,PIK3R5,RALA,RAP1A,RAP1B,REL,SOCS3,SOS1,SOS2,STAT3,TNF,TNFAIP6,TNFRSF1A,TNFRSF1B,TRAF6 |
| Acute Phase Response Signaling | 1.26E-13 | 0.21 | C1S,C2,C3,C4BPA,CEBPB,CFB,CP,GRB2,HP,IL1A,IL1B,IL1R1,IL1RAP,IL1RN,MAP3K5,MAPK14,NFKB1,NFKB2,NFKBIA,NFKBID,OSMR,PIK3CB,PIK3CG,RALA,RAP1A,RAP1B,REL,RIPK1,SERPINA3,SERPING1,SOCS3,SOD2,SOS1,SOS2,STAT3,TNF,TNFRSF1A,TNFRSF1B,TRAF6 |
| Natural Killer Cell Signaling | 5.01E-13 | 0.20 | CD226,CD48,FCER1G,FCGR2A,FYN,GRB2,IL12RB2,IL15,IL18R1,IL18RAP,JAK3,LCP2,LILRB1,LIMK2,MAP3K5,MAP3K8,MAPK14,NCK1,NFAT5,NFATC1,NFKB1,NFKB2,PAK2,PIK3CB,PIK3CG,PIK3R5,PTK2B,RALA,RAP1A,RAP1B,RASSF5,REL,SIGLEC7,SOS1,SOS2,STAT4,TRAF6,VAV1,WIPF1 |
| Granulocyte Adhesion and Diapedesis | 2.51E-12 | 0.21 | C5AR1,CCL2,CCL3,CCL4,CCR1,CCR2,CCR5,CCR6,CCR7,CSF3R,CXCL1,CXCL13,CXCL14,CXCL8,FPR1,FPR3,GNAI1,HRH1,ICAM1,ICAM2,IL18RAP,IL1A,IL1B,IL1R1,IL1R2,IL1RAP,IL1RN,ITGAM,MMP25,MSN,SELL,SELP,TNF,TNFRSF1A,TNFRSF1B,VCAM1 |
| TREM1 Signaling | 5.01E-12 | 0.32 | CASP1,CCL2,CCL3,CD40,CD86,CIITA,CXCL8,GRB2,ICAM1,IL1B,ITGAX,NFKB1,NFKB2,NLRP3,NOD2,REL,STAT3,STAT5B,TLR10,TLR2,TLR4,TLR8,TNF |
| PI3K/AKT Signaling | 5.01E-12 | 0.19 | CSF2RB,FOXO1,GAB2,GRB2,IFNAR1,IL10RA,IL10RB,IL12RB2,IL18R1,IL18RAP,IL1R1,IL1R2,IL21R,IL2RA,IL2RG,IL4R,ITGAM,ITGAX,JAK1,JAK3,MAP3K5,MAP3K8,NFKB1,NFKB2,NFKBIA,NFKBID,PIK3CB,PIK3CG,PPP2R1B,PPP2R5A,PPP2R5E,PTGS2,RALA,RAP1A,RAP1B,REL,SOS1,SOS2 |
| Production of Nitric Oxide and Reactive Oxygen Species in Macrophages | 5.01E-12 | 0.20 | CLU,CREBBP,CYBB,FNBP1,IRF1,JAK1,JAK3,MAP3K5,MAP3K8,MAPK14,NCF1,NCF2,NCF4,NFKB1,NFKB2,NFKBIA,NFKBID,PIK3CB,PIK3CG,PIK3R5,PPP2R1B,PPP2R5A,PPP2R5E,PRKCA,PRKCB,PRKCH,RAP1A,RAP1B,REL,RHOH,SIRPA,SPI1,TLR2,TLR4,TNF,TNFRSF1A,TNFRSF1B |
| Regulation Of The Epithelial Mesenchymal Transition By Growth Factors Pathway | 7.94E-12 | 0.19 | BRAF,CD40LG,DOCK10,FGF11,FGF9,FOXO1,GRB2,JAK1,JAK3,LATS1,LTB,MAPK14,MET,NFKB1,NFKB2,PIK3CB,PIK3CG,PIK3R5,RALA,RAP1A,RAP1B,REL,SMAD3,SMURF1,SOS1,SOS2,STAT3,TGFB2,TGFBR2,TNF,TNFRSF1A,TNFRSF1B,TNFSF13B,TNFSF14,TNFSF8,WWTR1,ZEB2 |
| Role of Pattern Recognition Receptors in Recognition of Bacteria and Viruses | 3.98E-11 | 0.21 | C3,C3AR1,C5AR1,CASP1,CD40LG,CLEC7A,CXCL8,IL15,IL1A,IL1B,LTB,NFKB1,NFKB2,NLRP3,NOD2,PIK3CB,PIK3CG,PIK3R5,PRKCA,PRKCB,PRKCH,REL,TGFB2,TLR2,TLR4,TLR8,TNF,TNFSF13B,TNFSF14,TNFSF8,TRAF6 |
| Phagosome Formation | 1.00E-10 | 0.11 | ACTR3,ADGRE1,ADGRE2,ADGRE3,ADGRF1,ADGRG3,ADGRG5,APBB1IP,C3,C3AR1,C5AR1,CCR1,CCR2,CCR5,CCR6,CCR7,CCRL2,CD14,CLEC4E,CLEC7A,CLIP1,CMKLR1,CR1,CYSLTR1,ELMO1,FCAR,FCER1G,FCGR2A,FCGR2C,FGR,FPR1,FPR3,FYN,GAB2,GPR155,GPR176,GPR65,GRB2,HCK,HRH1,ITGAM,ITGAX,ITPR1,LIMK2,LYN,MARCO,MSR1,P2RY10,P2RY6,PAK2,PIK3CB,PIK3CG,PIK3R5,PLA2G4A,PRKCA,PRKCB,PRKCH,PTAFR,PTGER2,PTGER3,PTGER4,PTK2B,RALA,RAP1A,RAP1B,RAPGEF1,RXFP1,SOS1,SOS2,TLR10,TLR2,TLR4,TLR8,VAV1,WIPF1 |
| Toll-like Receptor Signaling | 1.20E-10 | 0.29 | CD14,IL1A,IL1B,IL1RN,IRAK2,IRAK3,LY96,MAP4K4,MAPK14,NFKB1,NFKB2,NFKBIA,REL,TAB2,TLR10,TLR2,TLR4,TLR8,TNF,TNFAIP3,TRAF1,TRAF6 |
| CD40 Signaling | 2.95E-10 | 0.30 | CD40,CD40LG,ICAM1,JAK3,MAPK14,NFKB1,NFKB2,NFKBIA,NFKBID,PIK3CB,PIK3CG,PIK3R5,PTGS2,REL,STAT3,TANK,TNFAIP3,TRAF1,TRAF3,TRAF6 |
| HMGB1 Signaling | 3.24E-10 | 0.20 | CCL2,CD40LG,CXCL8,FNBP1,ICAM1,IL15,IL1A,IL1B,IL1R1,KAT6A,LTB,MAPK14,NFKB1,NFKB2,PIK3CB,PIK3CG,PIK3R5,RALA,RAP1A,RAP1B,REL,RHOH,TGFB2,TLR4,TNF,TNFRSF1A,TNFRSF1B,TNFSF13B,TNFSF14,TNFSF8,VCAM1 |
| IL-9 Signaling | 3.24E-10 | 0.43 | BCL3,IL2RG,IRS2,JAK1,JAK3,NFKB1,NFKB2,PIK3CB,PIK3CG,PIK3R5,REL,SOCS3,STAT3,STAT5B,TNF |
| Role of JAK1 and JAK3 in Cytokine Signaling | 3.55E-10 | 0.30 | GRB2,IL15,IL21R,IL2RA,IL2RG,IL4R,IL7,IRS2,JAK1,JAK3,PIK3CB,PIK3CG,PIK3R5,PTK2B,RALA,RAP1A,RAP1B,SOCS3,STAT3,STAT5B |
| JAK/STAT Signaling | 3.63E-10 | 0.27 | CEBPB,GNAQ,GRB2,JAK1,JAK3,NFKB1,NFKB2,PIAS1,PIK3CB,PIK3CG,PIK3R5,PTPN1,RALA,RAP1A,RAP1B,REL,SOCS3,SOS1,SOS2,STAT3,STAT4,STAT5B |
| fMLP Signaling in Neutrophils | 1.17E-09 | 0.21 | ACTR3,CYBB,FPR1,FPR3,GNA15,GNAI1,GNAQ,ITPR1,NCF1,NCF2,NFAT5,NFATC1,NFKB1,NFKB2,NFKBIA,NFKBID,PIK3CB,PIK3CG,PIK3R5,PPP3CC,PRKCA,PRKCB,PRKCH,RALA,RAP1A,RAP1B,REL |
| Glucocorticoid Receptor Signaling | 1.86E-09 | 0.12 | CCL2,CCL3,CD163,CD3D,CD3G,CREBBP,CSF2RB,CXCL8,EP300,FKBP5,GRB2,HP,ICAM1,IFNAR1,IL10RA,IL10RB,IL12RB2,IL18R1,IL18RAP,IL1A,IL1B,IL1R1,IL1R2,IL1RN,IL21R,IL2RA,IL2RG,IL4R,JAK1,JAK3,MAPK14,NCOA1,NCOA2,NCOA3,NFAT5,NFATC1,NFKB1,NFKB2,NFKBIA,NFKBID,PIK3CB,PIK3CG,PIK3R5,PLA2G4A,PPP3CC,PRKAA1,PTGS2,RALA,RAP1A,RAP1B,SLPI,SMAD3,SOS1,SOS2,STAT3,STAT5B,TGFB2,TGFBR2,TLR2,TNF,TRAF6,VCAM1 |
| GP6 Signaling Pathway | 1.95E-09 | 0.21 | ADAM10,APBB1IP,BTK,COL19A1,COL4A4,FCER1G,FYB1,FYN,GRAP2,GRB2,ITK,ITPR1,LAMB3,LAMC1,LCP2,LYN,NCK1,PIK3CB,PIK3CG,PIK3R5,PRKCA,PRKCB,PRKCH,RAP1B,RASGRP2,VAV1 |
| Pyroptosis Signaling Pathway | 2.04E-09 | 0.24 | AIM2,CASP1,CASP4,GBP2,IL1A,IL1B,IL1R1,IRF2,MAPK14,NFKB1,NFKB2,NLRP1,NLRP3,PTGER4,TLR10,TLR2,TLR4,TLR8,TNF,TNFRSF1A,TNFRSF1B,TRAF3 |
| IL-12 Signaling and Production in Macrophages | 2.04E-09 | 0.20 | CD40,CD40LG,CEBPB,CLU,EP300,IL12RB2,IRF1,MAP3K8,MAPK14,NCOA1,NFKB1,NFKB2,NFKBIA,PIK3CB,PIK3CG,PIK3R5,PRKCA,PRKCB,PRKCH,REL,SPI1,STAT4,TGFB2,TLR2,TLR4,TNF,TRAF6 |
| iNOS Signaling | 2.04E-09 | 0.35 | CD14,CREBBP,IRAK2,IRAK3,IRF1,JAK1,JAK3,LY96,MAPK14,NFKB1,NFKB2,NFKBIA,NFKBID,REL,TLR4,TRAF6 |
| IL-23 Signaling Pathway | 2.04E-09 | 0.35 | HIF1A,IL12RB2,IL1B,NFKB1,NFKB2,NFKBIA,PIK3CB,PIK3CG,PIK3R5,REL,RORA,RUNX1,SOCS3,STAT3,STAT4,TNF |
| IL-8 Signaling | 2.63E-09 | 0.17 | BRAF,CXCL1,CXCL8,CYBB,FNBP1,GNA15,GNAI1,GNAQ,ICAM1,IRAK2,IRAK3,ITGAM,ITGAX,LIMK2,MAP4K4,NCF1,NCF2,NFKB1,NFKBIA,PAK2,PIK3CB,PIK3CG,PIK3R5,PRKCA,PRKCB,PRKCH,PTGS2,PTK2B,RALA,RAP1A,RAP1B,RHOH,TRAF6,VCAM1 |
| ID1 Signaling Pathway | 3.98E-09 | 0.17 | ABI1,ACVR1C,BCL3,BLK,BRAF,CREBBP,EP300,FGR,FYN,HCK,HIF1A,LYN,MAPK14,NFKB1,NFKB2,PIK3CB,PIK3CG,PIK3R5,PLAC8,PTGER4,RALA,RAP1A,RAP1B,S100A9,SMAD1,SMAD3,STAT3,TGFB2,TGFBR2,TGM2,TNF,TNFRSF1A,TNFRSF1B |
| Prolactin Signaling | 4.68E-09 | 0.24 | CEBPB,CREBBP,EP300,FYN,GRB2,IRF1,PIK3CB,PIK3CG,PIK3R5,PRKCA,PRKCB,PRKCH,PRLR,RALA,RAP1A,RAP1B,SOCS3,SOS1,SOS2,STAT3,STAT5B |
| IL-7 Signaling Pathway | 9.55E-09 | 0.26 | BCL6,FOXO1,FYN,GRB2,IL2RG,IL7,JAK1,JAK3,LYN,MAPK14,MET,NFATC1,PAX5,PIK3CB,PIK3CG,PIK3R5,SOS1,SOS2,STAT5B |
| STAT3 Pathway | 1.02E-08 | 0.19 | CSF2RB,IFNAR1,IL10RA,IL10RB,IL12RB2,IL18R1,IL18RAP,IL1A,IL1B,IL1R1,IL1R2,IL21R,IL2RA,IL2RG,IL4R,MAPK14,NTRK3,PTPN2,RALA,RAP1A,RAP1B,SOCS3,STAT3,TGFA,TGFB2,TGFBR2 |
| G-Protein Coupled Receptor Signaling | 1.74E-08 | 0.10 | ADGRE1,ADGRE2,ADGRE3,ADGRF1,ADGRG3,ADGRG5,ARRB1,BRAF,C3AR1,C5AR1,CCR1,CCR2,CCR5,CCR6,CCR7,CCRL2,CMKLR1,CNGB1,CREBBP,CYSLTR1,FOXO1,FPR1,FPR3,FYN,GNA15,GNAI1,GNAQ,GPR155,GPR176,GPR65,GRB2,HRH1,KCNQ3,LATS1,MAP3K5,MAP3K8,MAPK14,MRTFA,NFAT5,NFATC1,NFKB1,NFKB2,NFKBIA,NFKBID,P2RY10,P2RY6,PAK2,PIK3CB,PIK3CG,PIK3R5,PPP3CC,PRKCA,PRKCB,PTAFR,PTGER2,PTGER3,PTGER4,PTK2B,RALA,RAP1A,RAP1B,RAPGEF1,REL,RGS18,RXFP1,SMPDL3A,SOS1,SOS2,STAT3,WWTR1 |
| Autophagy | 2.04E-08 | 0.16 | ATG7,CREBBP,DAPK1,DRAM1,FOXO1,HIF1A,IRS2,MAP1LC3B2,NFKB1,NFKB2,NOD2,OPTN,PIK3CB,PIK3CG,PIK3R5,PPP2R1B,PPP2R5A,PPP2R5E,PPP3CC,PRKAA1,RALA,RIPK1,SLC7A5,TFEB,TGFA,TGFB2,TLR4,TNF,TNFRSF1A,TNFRSF1B,TRAF3,TRAF6,UVRAG |
| Lymphotoxin β Receptor Signaling | 2.24E-08 | 0.30 | CREBBP,CXCL1,EP300,LTB,NFKB1,NFKB2,NFKBIA,NFKBID,PIK3CB,PIK3CG,PIK3R5,TNFSF14,TRAF1,TRAF3,TRAF6,VCAM1 |
| GM-CSF Signaling | 2.82E-08 | 0.26 | BCL2A1,CSF2RB,GRB2,HCK,LYN,PIK3CB,PIK3CG,PIK3R5,PPP3CC,PRKCB,RALA,RAP1A,RAP1B,RUNX1,SOS1,SOS2,STAT3,STAT5B |
| TGF-β Signaling | 3.31E-08 | 0.22 | ACVR1C,CREBBP,EP300,GRB2,MAP4K1,MAPK14,RALA,RAP1A,RAP1B,RNF111,RUNX2,RUNX3,SMAD1,SMAD3,SMURF1,SMURF2,SOS1,SOS2,TGFB2,TGFBR2,TRAF6 |
| IL-3 Signaling | 3.31E-08 | 0.24 | CSF2RB,FOXO1,GAB2,GRB2,JAK1,PIK3CB,PIK3CG,PIK3R5,PPP3CC,PRKCA,PRKCB,PRKCH,RALA,RAP1A,RAP1B,RAPGEF1,SOS1,STAT3,STAT5B |
| MSP-RON Signaling In Macrophages Pathway | 4.27E-08 | 0.20 | CIITA,CREBBP,GAB2,GRB2,ITGAM,NFKB1,NFKB2,NFKBIZ,PIK3CB,PIK3CG,PIK3R5,PTGS2,RALA,RAP1A,RAP1B,REL,SBNO2,SOCS3,SOS1,SOS2,STAT3,TLR4,TNF |
| Complement System | 4.27E-08 | 0.36 | C1S,C2,C3,C3AR1,C4BPA,C5AR1,CD55,CFB,CFH,CR1,ITGAM,ITGAX,SERPING1 |
| Inflammasome pathway | 5.75E-08 | 0.50 | AIM2,CASP1,CTSB,IL1B,NFKB1,NFKB2,NLRP1,NLRP3,NOD2,TLR4 |
| p38 MAPK Signaling | 8.13E-08 | 0.19 | CREBBP,DUSP10,IL18RAP,IL1A,IL1B,IL1R1,IL1R2,IL1RAP,IL1RN,IRAK2,IRAK3,MAP3K5,MAP4K1,MAPK14,PLA2G4A,RPS6KA3,TAB2,TGFB2,TGFBR2,TNF,TNFRSF1A,TNFRSF1B,TRAF6 |
| Fcγ Receptor-mediated Phagocytosis in Macrophages and Monocytes | 9.33E-08 | 0.22 | ACTR3,CBL,DGKB,FCGR2A,FGR,FYB1,FYN,GAB2,HCK,LCP2,LYN,MYO5A,NCF1,NCK1,PIK3CG,PRKCA,PRKCB,PRKCH,PTK2B,VAV1 |
| VDR/RXR Activation | 1.23E-07 | 0.23 | CD14,CEBPB,CYP24A1,EP300,FOXO1,MXD1,NCOA1,NCOA2,NCOA3,PPARD,PRKCA,PRKCB,PRKCH,RUNX2,SERPINB1,SPP1,TGFB2,THBD |
| NF-κB Signaling | 1.38E-07 | 0.12 | BRAF,CD3D,CD3G,CD40,CD40LG,CREBBP,EP300,FCER1G,IL1A,IL1B,IL1R1,IL1R2,IL1RN,IRAK3,MAP3K8,MAP4K4,NFKB1,NFKB2,NFKBIA,NFKBID,NTRK3,PIK3CB,PIK3CG,PIK3R5,PRKCB,RALA,RAP1A,RAP1B,RIPK1,TAB2,TANK,TGFA,TGFBR2,TLR10,TLR2,TLR4,TLR8,TNF,TNFAIP3,TNFRSF1A,TNFRSF1B,TNFSF13B,TRAF3,TRAF6 |
| IL-17 Signaling | 2.09E-07 | 0.16 | CCL2,CD40LG,CEBPB,CXCL1,CXCL8,DEFB1,IL15,IL1A,IL1B,JAK1,LCN2,LTB,MAPK14,NFKB1,PIK3CB,PIK3CG,PIK3R5,PTGS2,RALA,RAP1A,RAP1B,TAB2,TGFB2,TNF,TNFSF13B,TNFSF14,TNFSF8,TRAF6 |
| FAK Signaling | 2.09E-07 | 0.09 | ACTR3,ACVR1C,ADGRE1,ADGRE2,ADGRE3,ADGRF1,ADGRG3,ADGRG5,ASAP1,BCAR3,C3AR1,C5AR1,CCR1,CCR2,CCR5,CCR6,CCR7,CCRL2,CD3D,CD3G,CMKLR1,CSF2RB,CYSLTR1,ELK3,FCER1G,FPR1,FPR3,FYN,GPR155,GPR176,GPR65,GRB2,HRH1,IFNAR1,IL10RA,IL10RB,IL12RB2,IL18R1,IL18RAP,IL1R1,IL1R2,IL21R,IL2RA,IL2RG,IL4R,ITGAM,ITGAX,MAPK14,MET,NCK1,NCOA3,NFKB1,NFKB2,P2RY10,P2RY6,PAK2,PIK3CB,PIK3CG,PIK3R5,PTAFR,PTGER2,PTGER3,PTGER4,PTPN12,RALA,RAP1A,RAP1B,RXFP1,SH2D2A,SOCS3,SOS1,SOS2,TCF7L2,TGFB2,TGFBR2,WIPF1 |
| Fc Epsilon RI Signaling | 2.51E-07 | 0.19 | BTK,FCER1G,FYN,GRAP2,GRB2,LCP2,LYN,MAPK14,PIK3CB,PIK3CG,PIK3R5,PLA2G4A,PRKCA,PRKCB,PRKCH,RALA,RAP1A,RAP1B,SOS1,SOS2,TNF,VAV1 |
| p70S6K Signaling | 2.88E-07 | 0.18 | BTK,GNAI1,GNAQ,GRB2,IL2RG,IL4R,JAK1,LYN,PIK3CB,PIK3CG,PIK3R5,PLCL1,PPP2R1B,PPP2R5A,PPP2R5E,PRKCA,PRKCB,PRKCH,RALA,RAP1A,RAP1B,SOS1,SOS2 |
| Oncostatin M Signaling | 3.98E-07 | 0.30 | CHI3L1,EPAS1,GRB2,JAK1,JAK3,MT2A,OSMR,RALA,RAP1A,RAP1B,SOS1,STAT3,STAT5B |
| Role of PKR in Interferon Induction and Antiviral Response | 4.27E-07 | 0.17 | CASP1,IFNAR1,IL1B,IRF1,JAK1,MAPK14,MARCO,METAP2,MSR1,NFKB1,NFKB2,NFKBIA,NFKBID,NLRP1,NLRP3,REL,STAT3,TAB2,TLR4,TNF,TNFRSF1A,TRAF3,TRAF6 |
| Angiopoietin Signaling | 5.25E-07 | 0.22 | FOXO1,GRB2,NCK1,NFKB1,NFKB2,NFKBIA,NFKBID,PAK2,PIK3CB,PIK3CG,PIK3R5,RALA,RAP1A,RAP1B,REL,SOS1,STAT5B |
| LPS-stimulated MAPK Signaling | 5.25E-07 | 0.21 | CD14,MAP3K5,MAPK14,NFKB1,NFKB2,NFKBIA,NFKBID,PIK3CB,PIK3CG,PIK3R5,PRKCA,PRKCB,PRKCH,RALA,RAP1A,RAP1B,REL,TLR4 |
| Leukocyte Extravasation Signaling | 6.03E-07 | 0.15 | ARHGAP12,ARHGAP9,BTK,CYBB,GNAI1,ICAM1,ITGAM,ITK,MAPK14,MMP25,MSN,NCF1,NCF2,NCF4,PIK3CB,PIK3CG,PIK3R5,PRKCA,PRKCB,PRKCH,PTK2B,RAP1A,RAP1B,RASSF5,RHOH,VAV1,VCAM1,WIPF1 |
| TNFR2 Signaling | 6.17E-07 | 0.36 | BIRC3,NFKB1,NFKB2,NFKBIA,NFKBID,REL,TANK,TNF,TNFAIP3,TNFRSF1B,TRAF1 |
| IL-17A Signaling in Fibroblasts | 7.08E-07 | 0.32 | CCL2,CEBPB,CEBPD,LCN2,MAPK14,NFKB1,NFKB2,NFKBIA,NFKBID,NFKBIZ,REL,TRAF6 |
| IL-2 Signaling | 7.59E-07 | 0.25 | GRB2,IL2RA,IL2RG,JAK1,JAK3,PIK3CB,PIK3CG,PIK3R5,PTK2B,RALA,RAP1A,RAP1B,SOS1,SOS2,STAT5B |
| HIF1α Signaling | 8.32E-07 | 0.14 | BRAF,CREBBP,CYBB,EGLN1,EP300,HIF1A,HK1,HK3,MET,MMP25,NCF1,NCF2,NCOA1,PIK3CB,PIK3CG,PIK3R5,PPP3CC,PRKCA,PRKCB,PRKCH,RALA,RAP1A,RAP1B,SLC2A14,SLC2A3,SLC2A5,STAT3,TGFA,TGFB2 |
| Agranulocyte Adhesion and Diapedesis | 1.29E-06 | 0.14 | C5AR1,CCL2,CCL3,CCL4,CCR1,CCR2,CCR5,CCR6,CCR7,CXCL1,CXCL13,CXCL14,CXCL8,GNAI1,HRH1,ICAM1,ICAM2,IL1A,IL1B,IL1R1,IL1RN,MMP25,MSN,SELL,SELP,TNF,TNFRSF1A,VCAM1 |
| LXR/RXR Activation | 1.91E-06 | 0.17 | C3,CCL2,CD14,CLU,IL18RAP,IL1A,IL1B,IL1R1,IL1R2,IL1RAP,IL1RN,LY96,MSR1,NFKB1,NFKB2,PTGS2,REL,TLR4,TNF,TNFRSF1A,TNFRSF1B |
| ERBB Signaling | 1.91E-06 | 0.19 | FOXO1,GRB2,MAPK14,NCK1,NRG1,PAK2,PIK3CB,PIK3CG,PIK3R5,PRKCA,PRKCB,PRKCH,RALA,RAP1A,RAP1B,SOS1,SOS2,TGFA |
| B Cell Activating Factor Signaling | 2.75E-06 | 0.28 | MAPK14,NFAT5,NFATC1,NFKB1,NFKB2,NFKBIA,NFKBID,REL,TNFSF13B,TRAF1,TRAF3,TRAF6 |
| Regulation of the Epithelial-Mesenchymal Transition Pathway | 2.75E-06 | 0.14 | BRAF,FGF11,FGF9,GRB2,HIF1A,JAK1,JAK3,MET,NFKB1,NFKB2,PIK3CB,PIK3CG,PIK3R5,PSEN1,RALA,RAP1A,RAP1B,REL,SMAD3,SMURF1,SOS1,SOS2,STAT3,TCF7L2,TGFB2,TGFBR2,ZEB2 |
| CLEAR Signaling Pathway | 3.16E-06 | 0.12 | ATP6V1C1,CREBBP,CTSB,EP300,FNIP1,HIF1A,ITPR1,MAPK14,NTRK3,PPARGC1A,PPP2R1B,PPP2R5A,PPP2R5E,PPP3CC,PRKAA1,PRKCA,PRKCB,PRKCH,RALA,RAP1A,RAP1B,RRAGD,TFEB,TGFA,TGFB2,TGFBR2,TLR10,TLR2,TLR4,TLR8,TNF,TNFRSF1A,TNFRSF1B,UVRAG |
| NF-κB Activation by Viruses | 3.47E-06 | 0.21 | CCR5,NFKB1,NFKB2,NFKBIA,NFKBID,PIK3CB,PIK3CG,PIK3R5,PRKCA,PRKCB,PRKCH,RALA,RAP1A,RAP1B,REL,RIPK1 |
| HGF Signaling | 5.01E-06 | 0.16 | ELK3,GRB2,ITGAM,ITGAX,MAP3K5,MAP3K8,MET,PIK3CB,PIK3CG,PIK3R5,PRKCA,PRKCB,PRKCH,PTGS2,RALA,RAP1A,RAP1B,RAPGEF1,SOS1,SOS2,STAT3 |
| Activation of IRF by Cytosolic Pattern Recognition Receptors | 6.03E-06 | 0.22 | CD40,CREBBP,IFNAR1,NFKB1,NFKB2,NFKBIA,NFKBID,REL,RIPK1,TANK,TNF,TRAF3,TRAF6,ZBP1 |
| Crosstalk between Dendritic Cells and Natural Killer Cells | 6.03E-06 | 0.19 | CCR7,CD226,CD28,CD40,CD40LG,CD80,CD86,CSF2RB,IL15,IL2RG,LTB,NFKB1,NFKB2,REL,TLR4,TNF,TNFRSF1B |
| Role of MAPK Signaling in Inhibiting the Pathogenesis of Influenza | 8.51E-06 | 0.20 | BRAF,CCL2,CXCL8,IL1B,MAP3K5,MAPK14,NFKB1,NFKB2,NFKBIA,NFKBID,PLA2G4A,PTGS2,RPS6KA3,TLR4,TNF |
| Death Receptor Signaling | 9.33E-06 | 0.18 | ARHGDIB,BIRC3,CFLAR,MAP3K5,MAP4K4,NFKB1,NFKB2,NFKBIA,NFKBID,PARP15,REL,RIPK1,TANK,TNF,TNFRSF10A,TNFRSF1A,TNFRSF1B |
| Xenobiotic Metabolism Signaling | 1.02E-05 | 0.12 | CHST11,CHST15,CREBBP,CYP2C19,CYP2C9,EP300,HS3ST3B1,IL1A,IL1B,IL4I1,MAP3K5,MAP3K8,MAPK14,NCOA1,NFE2L2,NFKB1,NFKB2,PIK3CB,PIK3CG,PIK3R5,PPARGC1A,PPP2R1B,PPP2R5A,PPP2R5E,PRKCA,PRKCB,PRKCH,RALA,RAP1A,RAP1B,REL,TNF |
| B Cell Receptor Signaling | 1.15E-05 | 0.11 | APBB1IP,BCL2A1,BCL6,BTK,CREBBP,DAPP1,FCGR2A,FCGR2C,FOXO1,GAB2,GRB2,LYN,MAP3K5,MAP3K8,MAPK14,NFAT5,NFATC1,NFKB1,NFKB2,NFKBIA,NFKBID,PAX5,PIK3AP1,PIK3CB,PIK3CG,PIK3R5,POU2F2,PPP3CC,PRKCB,PTK2B,PTPRC,RALA,RAP1A,RAP1B,RASSF5,REL,SOS1,SOS2,VAV1 |
| TNFR1 Signaling | 1.32E-05 | 0.24 | BIRC3,NFKB1,NFKB2,NFKBIA,NFKBID,PAK2,REL,RIPK1,TANK,TNF,TNFAIP3,TNFRSF1A |
| Apelin Endothelial Signaling Pathway | 1.35E-05 | 0.15 | CCL2,GNA15,GNAI1,GNAQ,HIF1A,ICAM1,NFKB1,NFKB2,PIK3CB,PIK3CG,PIK3R5,PRKAA1,PRKCA,PRKCB,PRKCH,RALA,RAP1A,RAP1B,REL,SMAD3,VCAM1 |
| April Mediated Signaling | 1.35E-05 | 0.26 | MAPK14,NFAT5,NFATC1,NFKB1,NFKB2,NFKBIA,NFKBID,REL,TRAF1,TRAF3,TRAF6 |
| Gαq Signaling | 1.62E-05 | 0.14 | BTK,FNBP1,GNA15,GNAI1,GNAQ,HRH1,ITPR1,NFATC1,NFKB1,NFKB2,NFKBIA,NFKBID,PIK3CB,PIK3CG,PIK3R5,PPP3CC,PRKCA,PRKCB,PRKCH,PTK2B,REL,RGS18,RHOH |
| IL-15 Production | 1.70E-05 | 0.16 | ABL2,BLK,BTK,EPHA4,FGR,FYN,HCK,IL15,IRF1,ITK,JAK1,JAK3,LYN,MET,NFKB1,NFKB2,NTRK3,PTK2B,REL |
| MIF-mediated Glucocorticoid Regulation | 2.04E-05 | 0.28 | CD14,LY96,NFKB1,NFKB2,NFKBIA,NFKBID,PLA2G4A,PTGS2,REL,TLR4 |
| Role of Hypercytokinemia/hyperchemokinemia in the Pathogenesis of Influenza | 2.04E-05 | 0.19 | CASP1,CCL2,CCL3,CCL4,CXCL8,IFNAR1,IL1A,IL1B,IL1RN,JAK1,NFKB1,NFKB2,NLRP3,TLR4,TNF |
| IL-4 Signaling | 2.04E-05 | 0.18 | GRB2,IL2RG,IL4R,IRF4,JAK1,JAK3,NFAT5,NFATC1,PIK3CB,PIK3CG,PIK3R5,RALA,RAP1A,RAP1B,SOS1,SOS2 |
| Role of RIG1-like Receptors in Antiviral Innate Immunity | 2.09E-05 | 0.25 | CREBBP,EP300,NFKB1,NFKB2,NFKBIA,NFKBID,REL,RIPK1,TANK,TRAF3,TRAF6 |
| RAC Signaling | 3.16E-05 | 0.15 | ACTR3,ANK1,CYBB,ELK4,ITGAM,ITGAX,LIMK2,NCF1,NCF2,NFKB1,NFKB2,PAK2,PIK3CB,PIK3CG,PIK3R5,PTK2B,RALA,RAP1A,RAP1B,REL |
| IL-13 Signaling Pathway | 3.31E-05 | 0.16 | BLK,CHI3L1,DEFB1,FGR,FOSL2,FYN,HCK,IL4R,JAK1,LYN,MAPK14,PIK3CB,PIK3CG,PIK3R5,SOCS3,STAT3,TGFB2,TRAF6 |
| PEDF Signaling | 3.63E-05 | 0.18 | CFLAR,MAPK14,NFKB1,NFKB2,NFKBIA,NFKBID,PIK3CB,PIK3CG,PIK3R5,RALA,RAP1A,RAP1B,REL,SOD2,TCF7L2 |
| Ephrin Receptor Signaling | 4.47E-05 | 0.12 | ABI1,ACTR3,ADAM10,CREBBP,EPHA4,FYN,GNA15,GNAI1,GNAQ,GRB2,ITGAM,ITGAX,LIMK2,MAP4K4,NCK1,PAK2,PIK3CG,RALA,RAP1A,RAP1B,RAPGEF1,SOS1,SOS2,STAT3,WIPF1 |
| T Cell Receptor Signaling | 4.47E-05 | 0.10 | CBL,CD28,CD3D,CD3G,CD80,CD86,FCER1G,FYB1,FYN,GRAP2,GRB2,ICAM1,ICOS,ITK,LCP2,MAPK14,NCK1,NFAT5,NFATC1,NFKB1,NFKB2,NFKBIA,PIK3CB,PIK3CG,PIK3R5,PPP3CC,PTK2B,PTPN22,PTPRC,RALA,RAP1A,RAP1B,REL,SOS1,SOS2,TCF7L2,TNF,VAV1 |
| IL-1 Signaling | 4.47E-05 | 0.17 | GNA15,GNAI1,GNAQ,IL1A,IL1R1,IL1RAP,IRAK2,IRAK3,MAPK14,NFKB1,NFKB2,NFKBIA,NFKBID,REL,TAB2,TRAF6 |
| FGF Signaling | 4.68E-05 | 0.17 | CREBBP,FGF11,FGF9,GRB2,ITPR1,MAP3K5,MAPK14,MET,PIK3CB,PIK3CG,PIK3R5,PRKCA,SOS1,SOS2,STAT3 |
| PDGF Signaling | 4.68E-05 | 0.17 | ABL2,GRB2,JAK1,JAK3,PIK3CB,PIK3CG,PIK3R5,PRKCA,PRKCB,RALA,RAP1A,RAP1B,SOS1,SOS2,STAT3 |
| Macropinocytosis Signaling | 4.68E-05 | 0.18 | ABI1,CD14,CSF1,MET,PIK3CB,PIK3CG,PIK3R5,PRKCA,PRKCB,PRKCH,RAB5A,RALA,RAP1A,RAP1B |
| ERK/MAPK Signaling | 4.68E-05 | 0.12 | BRAF,CREBBP,ELK3,FYN,GRB2,ITGAM,ITGAX,NFATC1,PAK2,PIK3CB,PIK3CG,PIK3R5,PLA2G4A,PPP2R1B,PPP2R5A,PPP2R5E,PRKCA,PRKCB,PTK2B,RALA,RAP1A,RAP1B,RAPGEF1,SOS1,SOS2,STAT3 |
| Role of JAK2 in Hormone-like Cytokine Signaling | 4.79E-05 | 0.28 | IRS2,JAK1,PRLR,PTPN1,SH2B3,SIRPA,SOCS3,STAT3,STAT5B |
| Estrogen Receptor Signaling | 7.24E-05 | 0.10 | CREBBP,EP300,FOXO1,GNA15,GNAI1,GNAQ,GRB2,HIF1A,JAK1,JAK3,LEPR,LIMK2,MCU,MED13,MED13L,MMP25,NCOA1,NCOA2,NCOA3,NFKB1,NFKB2,NRF1,PIK3CB,PIK3CG,PIK3R5,PLCL1,PPARGC1A,PRKAA1,PRKCA,PRKCB,PRKCH,RALA,RAP1A,RAP1B,REL,RUNX2,SOD2,SOS1,SOS2 |
| Ceramide Signaling | 7.76E-05 | 0.17 | NFKB1,NFKB2,PIK3CB,PIK3CG,PIK3R5,PPP2R1B,PPP2R5A,PPP2R5E,RALA,RAP1A,RAP1B,REL,TNF,TNFRSF1A,TNFRSF1B |
| 4-1BB Signaling in T Lymphocytes | 7.94E-05 | 0.27 | MAP3K5,MAPK14,NFKB1,NFKB2,NFKBIA,NFKBID,REL,TNFRSF9,TRAF1 |
| TEC Kinase Signaling | 8.51E-05 | 0.10 | BLK,BTK,CD3D,CD3G,FCER1G,FGR,FNBP1,FYN,GNA15,GNAI1,GNAQ,HCK,ITGAM,ITGAX,ITK,JAK1,JAK3,LYN,NFKB1,NFKB2,PAK2,PIK3CB,PIK3CG,PIK3R5,PRKCA,PRKCB,PRKCH,PTK2B,REL,RHOH,STAT3,STAT4,STAT5B,TLR4,TNF,TNFRSF10A,VAV1 |
| Opioid Signaling Pathway | 8.71E-05 | 0.11 | ARRB1,BLK,BRAF,CREBBP,EP300,FGR,FYN,GNA15,GNAI1,GNAQ,HCK,ITPR1,LYN,NFKB1,NFKB2,NFKBIA,PIK3CG,PPP3CC,PRKCA,PRKCB,PRKCH,RALA,RAP1A,RAP1B,RGS18,RGS5,RPS6KA3,SOS1,SOS2,TCF7L2 |
| Airway Pathology in Chronic Obstructive Pulmonary Disease | 8.71E-05 | 0.15 | CCL2,CD40LG,CXCL1,CXCL8,FGF11,FGF9,IL15,IL1A,IL1B,LCN2,LTB,PAEP,TGFB2,TNF,TNFSF13B,TNFSF14,TNFSF8 |
| Apoptosis Signaling | 8.71E-05 | 0.16 | BCL2A1,BIRC3,MAP3K5,MAP4K4,NFKB1,NFKB2,NFKBIA,NFKBID,PRKCA,RALA,RAP1A,RAP1B,REL,TNF,TNFRSF1A,TNFRSF1B |
| Endothelin-1 Signaling | 9.33E-05 | 0.12 | BRAF,CASP1,CASP4,GNA15,GNAI1,GNAQ,GRB2,ITPR1,MAPK14,PIK3CB,PIK3CG,PIK3R5,PLA2G4A,PLCL1,PRKCA,PRKCB,PRKCH,PTGER2,PTGS2,RALA,RAP1A,RAP1B,SOS1 |
