## Supplemental Table 1-10 for "miRNA/mRNA analysis of increased TGF-β pathways drive epithelial-mesenchymal transition and regulatory T cell differentiation": Supp Table 6.docx

**Supp Table 6. Ingenuity canonical pathways enriched by significantly downregulated mRNAs in Endo+ compared to Endo- with FDR<0.05.**

| **Ingenuity Canonical Pathways** | **FDR** | **Ratio** | **Molecules** |
| --- | --- | --- | --- |
| Kinetochore Metaphase Signaling Pathway | 1.00E-19 | 0.34 | ANAPC11,AURKB,BIRC5,BOD1,BUB1B,CCNB1,CDC20,CDCA8,CDK1,CENPA,CENPE,CENPH,CENPN,CENPU,DSN1,ESPL1,H2AC21,H2AX,H2AZ1,INCENP,KIF2C,MAD2L1,MAD2L2,MXD3,NDC80,NEK2,PLK1,PMF1/PMF1-BGLAP,PTTG1,REC8,SKA1, SKA3,SPC24,SPC25,TTK,ZWINT |
| Cell Cycle Control of Chromosomal Replication | 1.58E-12 | 0.38 | CDC45,CDC6,CDK1,CDK16,CDK20,CDK4,CDK5,CDT1,DNA2,LIG1,MCM2,MCM3,MCM4,MCM5,MCM7,ORC6,PCNA,POLA2,POLD1,POLE,TOP2A |
| Granzyme A Signaling | 2.45E-10 | 0.30 | APEX1,H1-1,H1-10,H1-2,H1-3,H1-4,H1-5, HMGB2, LMNB2,NDUFA11,NDUFA13,NDUFA2,NDUFA4,NDUFA8,NDUFB11,NDUFB7,NDUFS3,NDUFS8,NDUFV1,NME1,PARP1 |
| Oxidative Phosphorylation | 1.86E-08 | 0.22 | ATP5F1D,ATP5MC2,ATP5MC3,ATP5ME,ATP5PB,ATP5PO,COX4I1,COX6C,COX8A,CYC1,NDUFA11,NDUFA13,NDUFA2,NDUFA4,NDUFA8,NDUFB11,NDUFB7,NDUFS3,NDUFS8,NDUFV1,SURF1,UQCR10,UQCRC1 |
| DNA Methylation and Transcriptional Repression Signaling | 2.40E-07 | 0.20 | CDK1,CDK16,CDK20,CDK4,CDK5,E2F1,E2F7,E2F8,GLI1,H4C1,H4C13,H4C16,H4C2,H4C3,H4C4,H4C6,H4C8,MBD3,RBBP7,TCF19,TP53,UHRF1 |
| Mitochondrial Dysfunction | 2.88E-07 | 0.16 | APH1A,ATP5F1D,ATP5MC2,ATP5MC3,ATP5ME,ATP5PB,ATP5PO,COX4I1,COX6C,COX8A,CYC1,HTRA2,NDUFA11,NDUFA13,NDUFA2,NDUFA4,NDUFA8,NDUFB11,NDUFB7,NDUFS3,NDUFS8,NDUFV1,PARK7,SURF1,TXN2,UQCR10,UQCRC1 |
| Sirtuin Signaling Pathway | 2.51E-06 | 0.12 | ACADL,APEX1,ATP5F1D,ATP5PB,CYC1,E2F1,H1-1,H1-10,H1-2,H1-3,H1-4,H1-5, H3C3, LDHB, NDUFA11, NDUFA13, NDUFA2,NDUFA4,NDUFA8,NDUFB11,NDUFB7,NDUFS3,NDUFS8,NDUFV1,PARP1,POLR1C,RRP9,SLC25A5,TIMM10,TIMM13,TIMM17B,TIMM8A,TP53,TRIM28,TUBA1B |
| NAD Signaling Pathway | 4.68E-06 | 0.16 | ACADL,H1-1,H1-10,H1-2,H1-3,H1-4,H1-5,H2BC10,H2BC15,H2BC17,H2BC5,H2BC8,LDHB,PARP1,PARP10,PIK3R2,POLR2C,POLR2E,POLR2G,POLRMT,SLC29A1,SLC29A2,TP53 |
| Role of CHK Proteins in Cell Cycle Checkpoint Control | 2.40E-04 | 0.21 | CDC25C,CDK1,CLSPN,E2F1,E2F7,E2F8,PCNA,PLK1,PPP2R1A,RFC2,RFC5,TP53 |
| Pyrimidine Deoxyribonucleotides De Novo Biosynthesis I | 1.35E-03 | 0.30 | DTYMK,DUT,NME1,NME2,NME3,NME4,RRM2 |
| Ribonucleotide Reductase Signaling Pathway | 1.82E-03 | 0.12 | BIRC5,CCNF,CDK1,CDK4,CREB3L1,CREB3L4,E2F1,E2F7,E2F8,EIF4EBP1,FOXM1,H3C3,MLST8,MYBL2,PARP1,PARP10,PIK3R2,RRM2,TP53,WEE1 |
| Role of BRCA1 in DNA Damage Response | 5.50E-03 | 0.15 | E2F1,E2F7,E2F8,FANCA,FANCB,FANCD2,FANCG,MSH2,PLK1,RFC2,RFC5,TP53 |
| Salvage Pathways of Pyrimidine Ribonucleotides | 7.94E-03 | 0.14 | CDK1,CDK4,CDK5,NEK2,NME1,NME2,NME3,NME4,PAK5,PLK1,PYCR3,TTK,UCK1 |
| Cyclins and Cell Cycle Regulation | 7.94E-03 | 0.14 | CCNA2,CCNB1,CCNB2,CDK1,CDK4,E2F1,E2F7,E2F8,HDAC6,PPP2R1A,TP53,WEE1 |
| Pyrimidine Ribonucleotides De Novo Biosynthesis | 2.82E-02 | 0.18 | CAD,ENTPD2,NME1,NME2,NME3,NME4,NTPCR |
| ATM Signaling | 3.09E-02 | 0.12 | BRAT1,CCNB1,CCNB2,CDC25C,CDK1,CREB3L1,CREB3L4,FANCD2,H2AX,PPP2R1A,TP53,TRIM28 |
| GDP-mannose Biosynthesis | 3.24E-02 | 0.50 | GMPPB,MPI,PMM1 |
| γ-glutamyl Cycle | 3.98E-02 | 0.31 | GCLC,GGCT,GGT6,GSS |
