## Supplemental Table 1-10 for "miRNA/mRNA analysis of increased TGF-β pathways drive epithelial-mesenchymal transition and regulatory T cell differentiation": Supp Table 7.docx

| **Supp Table 7. Significantly upregulated miRNAs in Endo+ compared to Endo- biopsies with FDR<0.05 (Sorted by descending fold changes)** | | | |
| --- | --- | --- | --- |
| **miRNA symbol** | **Fold Change** | **P value** | **FDR** |
| miR-7150 | 9.2 | 7.69E-09 | 6.48E-07 |
| miR-6752-5p | 8.5 | 2.44E-09 | 4.65E-07 |
| miR-4530 | 6.1 | 3.68E-09 | 4.65E-07 |
| miR-3197 | 4.1 | 5.23E-05 | 7.35E-04 |
| miR-6516-5p | 3.9 | 2.18E-07 | 7.88E-06 |
| miR-3687 | 3.8 | 3.93E-06 | 9.94E-05 |
| miR-198 | 3.5 | 1.34E-04 | 1.41E-03 |
| miR-6088 | 3.4 | 6.54E-04 | 4.04E-03 |
| miR-1291 | 3.3 | 3.83E-04 | 2.77E-03 |
| miR-4417 | 3.3 | 8.01E-07 | 2.25E-05 |
| miR-664b-5p | 3.2 | 3.08E-04 | 2.57E-03 |
| miR-664a-5p | 3.2 | 1.26E-04 | 1.39E-03 |
| miR-1244 | 3.2 | 1.28E-05 | 2.39E-04 |
| miR-6756-5p | 3.0 | 3.37E-04 | 2.58E-03 |
| miR-664b-3p | 2.8 | 1.06E-03 | 5.71E-03 |
| miR-149-3p | 2.7 | 6.32E-03 | 2.35E-02 |
| miR-3663-5p | 2.7 | 1.00E-03 | 5.52E-03 |
| miR-4444 | 2.6 | 1.10E-05 | 2.33E-04 |
| miR-6085 | 2.4 | 5.84E-03 | 2.24E-02 |
| let-7b-5p | 2.3 | 5.66E-05 | 7.54E-04 |
| miR-574-5p | 2.3 | 5.38E-03 | 2.09E-02 |
| miR-6803-5p | 2.3 | 3.21E-04 | 2.57E-03 |
| miR-4496 | 2.2 | 1.05E-02 | 3.47E-02 |
| miR-3654 | 2.2 | 5.88E-04 | 3.72E-03 |
| miR-6124 | 2.2 | 1.66E-02 | 4.77E-02 |
| miR-6845-5p | 2.1 | 3.67E-03 | 1.55E-02 |
| miR-4271 | 2.1 | 6.17E-03 | 2.33E-02 |
| miR-6131 | 2.1 | 1.34E-02 | 4.14E-02 |
| miR-146a-5p | 2.1 | 1.70E-02 | 4.85E-02 |
| let-7c-5p | 2.1 | 1.52E-04 | 1.54E-03 |
| miR-2861 | 2.1 | 3.25E-04 | 2.57E-03 |
| miR-6727-5p | 2.0 | 7.02E-04 | 4.13E-03 |
| miR-663b | 2.0 | 9.45E-04 | 5.31E-03 |
| miR-5739 | 1.9 | 1.60E-02 | 4.71E-02 |
| miR-320b | 1.9 | 1.75E-05 | 2.95E-04 |
| miR-320a | 1.9 | 2.59E-05 | 3.85E-04 |
| miR-4429 | 1.9 | 1.32E-05 | 2.39E-04 |
| miR-320c | 1.9 | 1.94E-05 | 3.07E-04 |
| miR-6165 | 1.9 | 1.56E-02 | 4.63E-02 |
| miR-1237-5p | 1.9 | 8.10E-03 | 2.85E-02 |
| miR-320d | 1.8 | 8.90E-05 | 1.13E-03 |
| miR-3940-5p | 1.8 | 1.12E-02 | 3.59E-02 |
| miR-664a-3p | 1.8 | 5.41E-04 | 3.53E-03 |
| miR-8069 | 1.7 | 1.37E-02 | 4.18E-02 |
| miR-30a-3p | 1.7 | 8.51E-03 | 2.95E-02 |
| miR-197-3p | 1.7 | 1.24E-02 | 3.92E-02 |
| miR-320e | 1.7 | 1.72E-03 | 8.22E-03 |
