## Supplemental Table 1-10 for "miRNA/mRNA analysis of increased TGF-β pathways drive epithelial-mesenchymal transition and regulatory T cell differentiation": Supp Table 8.docx

| **Supp Table 8. Significantly downregulated miRNAs in Endo+ compared to Endo- biopsies with FDR<0.05 (Sorted by ascending fold changes)** | | | |
| --- | --- | --- | --- |
| **miRNA symbol** | **Fold Change** | **P value** | **FDR** |
| miR-1285-5p | 0.15 | 3.79E-04 | 2.77E-03 |
| miR-223-3p | 0.19 | 7.53E-03 | 2.68E-02 |
| miR-141-3p | 0.22 | 5.87E-08 | 3.71E-06 |
| miR-362-3p | 0.22 | 2.09E-07 | 7.88E-06 |
| miR-1254 | 0.24 | 1.78E-04 | 1.61E-03 |
| miR-142-5p | 0.26 | 9.45E-03 | 3.23E-02 |
| miR-4734 | 0.27 | 1.21E-03 | 6.23E-03 |
| miR-429 | 0.29 | 8.84E-06 | 2.03E-04 |
| miR-200a-3p | 0.32 | 8.02E-07 | 2.25E-05 |
| miR-566 | 0.32 | 1.91E-03 | 8.76E-03 |
| miR-126-5p | 0.32 | 7.73E-04 | 4.45E-03 |
| miR-1273d | 0.36 | 3.74E-03 | 1.55E-02 |
| miR-34c-5p | 0.36 | 1.12E-02 | 3.59E-02 |
| miR-224-5p | 0.37 | 6.75E-04 | 4.07E-03 |
| miR-19a-3p | 0.37 | 1.88E-03 | 8.76E-03 |
| miR-374b-5p | 0.39 | 1.64E-03 | 7.98E-03 |
| miR-101-3p | 0.39 | 1.03E-02 | 3.46E-02 |
| miR-30b-5p | 0.40 | 2.07E-03 | 9.20E-03 |
| miR-19b-3p | 0.40 | 4.12E-04 | 2.89E-03 |
| miR-21-5p | 0.44 | 6.90E-03 | 2.53E-02 |
| miR-335-5p | 0.44 | 1.34E-03 | 6.77E-03 |
| miR-660-5p | 0.44 | 2.43E-04 | 2.12E-03 |
| miR-376c-3p | 0.45 | 2.15E-03 | 9.39E-03 |
| miR-194-5p | 0.46 | 1.70E-04 | 1.59E-03 |
| miR-18a-5p | 0.47 | 5.44E-04 | 3.53E-03 |
| miR-30e-5p | 0.47 | 1.60E-03 | 7.95E-03 |
| miR-106b-5p | 0.49 | 1.17E-04 | 1.34E-03 |
| miR-20a-5p | 0.50 | 4.86E-03 | 1.92E-02 |
| miR-20b-5p | 0.51 | 7.39E-08 | 3.74E-06 |
| miR-376a-3p | 0.51 | 1.63E-02 | 4.75E-02 |
| miR-148a-3p | 0.53 | 5.15E-04 | 3.52E-03 |
| miR-27a-3p | 0.55 | 9.72E-05 | 1.17E-03 |
| miR-455-5p | 0.56 | 1.05E-02 | 3.47E-02 |
| miR-193a-3p | 0.57 | 4.23E-03 | 1.70E-02 |
| miR-200c-3p | 0.59 | 1.26E-02 | 3.93E-02 |
| miR-17-5p | 0.60 | 1.65E-04 | 1.59E-03 |
| miR-186-5p | 0.62 | 2.05E-03 | 9.20E-03 |
| miR-130b-3p | 0.63 | 7.17E-03 | 2.59E-02 |
| miR-130a-3p | 0.63 | 1.14E-03 | 6.02E-03 |
| miR-27b-3p | 0.66 | 4.10E-03 | 1.67E-02 |
| miR-30c-5p | 0.68 | 1.48E-02 | 4.45E-02 |
| miR-148b-3p | 0.73 | 2.50E-03 | 1.07E-02 |
