## Supplemental Table 1-10 for "miRNA/mRNA analysis of increased TGF-β pathways drive epithelial-mesenchymal transition and regulatory T cell differentiation": Supp Table 9.docx

| **Supp Table 9. Ingenuity canonical pathways enriched by significantly downregulated miRNA-upregulated mRNA pairs in Endo+ compared to Endo- women with FDR<0.05 (Top 100)** | |
| --- | --- |
| **Pathways** | **FDR** |
| Pathogen Induced Cytokine Storm Signaling Pathway | 1.58E-14 |
| Regulation Of the Epithelial Mesenchymal Transition by Growth Factors Pathway | 1.58E-14 |
| Regulation of the Epithelial-Mesenchymal Transition Pathway | 2.51E-12 |
| IL-6 Signaling | 2.51E-12 |
| Oncostatin M Signaling | 2.51E-12 |
| JAK/STAT Signaling | 2.51E-12 |
| Th1 and Th2 Activation Pathway | 2.51E-12 |
| S100 Family Signaling Pathway | 7.94E-12 |
| PPARα/RXRα Activation | 7.94E-12 |
| FAK Signaling | 1.00E-11 |
| IL-10 Signaling | 1.10E-10 |
| Role of JAK1 and JAK3 in γc Cytokine Signaling | 1.23E-10 |
| IL-7 Signaling Pathway | 2.69E-10 |
| T Cell Exhaustion Signaling Pathway | 6.61E-10 |
| IL-17 Signaling | 6.76E-10 |
| Macrophage Classical Activation Signaling Pathway | 7.08E-10 |
| Macrophage Alternative Activation Signaling Pathway | 1.51E-09 |
| ERB2-ERBB3 Signaling | 1.55E-09 |
| Acute Myeloid Leukemia Signaling | 1.55E-09 |
| HMGB1 Signaling | 2.00E-09 |
| ID1 Signaling Pathway | 2.09E-09 |
| Prolactin Signaling | 2.45E-09 |
| IL-23 Signaling Pathway | 2.57E-09 |
| TGF-β Signaling | 2.63E-09 |
| Systemic Lupus Erythematosus in T Cell Signaling Pathway | 2.82E-09 |
| GM-CSF Signaling | 2.95E-09 |
| CREB Signaling in Neurons | 2.95E-09 |
| Th2 Pathway | 3.63E-09 |
| Insulin Receptor Signaling | 4.57E-09 |
| Mouse Embryonic Stem Cell Pluripotency | 4.68E-09 |
| NOD1/2 Signaling Pathway | 5.01E-09 |
| Telomerase Signaling | 5.13E-09 |
| IGF-1 Signaling | 5.13E-09 |
| MSP-RON Signaling in Cancer Cells Pathway | 5.13E-09 |
| Role of Macrophages, Fibroblasts and Endothelial Cells in Rheumatoid Arthritis | 5.13E-09 |
| Tumor Microenvironment Pathway | 5.13E-09 |
| HER-2 Signaling in Breast Cancer | 5.75E-09 |
| Renal Cell Carcinoma Signaling | 6.31E-09 |
| IL-3 Signaling | 7.08E-09 |
| Molecular Mechanisms of Cancer | 7.59E-09 |
| IL-9 Signaling | 8.91E-09 |
| MSP-RON Signaling in Macrophages Pathway | 1.20E-08 |
| Role Of Osteoclasts in Rheumatoid Arthritis Signaling Pathway | 1.41E-08 |
| Pulmonary Healing Signaling Pathway | 1.45E-08 |
| PDGF Signaling | 1.51E-08 |
| Renin-Angiotensin Signaling | 1.74E-08 |
| Ephrin Receptor Signaling | 1.74E-08 |
| IL-2 Signaling | 1.95E-08 |
| G-Protein Coupled Receptor Signaling | 2.51E-08 |
| ERBB Signaling | 2.57E-08 |
| HGF Signaling | 3.47E-08 |
| Myelination Signaling Pathway | 3.98E-08 |
| Granulocyte Adhesion and Diapedesis | 4.17E-08 |
| Pulmonary Fibrosis Idiopathic Signaling Pathway | 4.27E-08 |
| Osteoarthritis Pathway | 7.76E-08 |
| Cardiac Hypertrophy Signaling (Enhanced) | 8.51E-08 |
| Production of Nitric Oxide and Reactive Oxygen Species in Macrophages | 8.71E-08 |
| Angiopoietin Signaling | 8.91E-08 |
| Role of JAK family kinases in IL-6-type Cytokine Signaling | 9.77E-08 |
| Adrenomedullin signaling pathway | 1.10E-07 |
| PTEN Signaling | 1.20E-07 |
| Neuroinflammation Signaling Pathway | 1.86E-07 |
| CNTF Signaling | 2.00E-07 |
| Role of NANOG in Mammalian Embryonic Stem Cell Pluripotency | 2.34E-07 |
| Pancreatic Adenocarcinoma Signaling | 2.69E-07 |
| T Cell Receptor Signaling | 2.82E-07 |
| ERK/MAPK Signaling | 3.09E-07 |
| Actin Nucleation by ARP-WASP Complex | 3.24E-07 |
| Chronic Myeloid Leukemia Signaling | 3.47E-07 |
| Phagosome Formation | 4.07E-07 |
| Role of Osteoblasts, Osteoclasts and Chondrocytes in Rheumatoid Arthritis | 4.07E-07 |
| Thrombopoietin Signaling | 4.17E-07 |
| VEGF Signaling | 5.13E-07 |
| CLEAR Signaling Pathway | 5.13E-07 |
| Estrogen Receptor Signaling | 5.13E-07 |
| ERBB4 Signaling | 5.50E-07 |
| Paxillin Signaling | 9.77E-07 |
| Natural Killer Cell Signaling | 9.77E-07 |
| Agranulocyte Adhesion and Diapedesis | 9.77E-07 |
| IL-22 Signaling | 1.10E-06 |
| GDNF Family Ligand-Receptor Interactions | 1.41E-06 |
| Airway Pathology in Chronic Obstructive Pulmonary Disease | 1.51E-06 |
| Thyroid Cancer Signaling | 1.51E-06 |
| Epithelial Adherens Junction Signaling | 1.55E-06 |
| Neurotrophin/TRK Signaling | 1.62E-06 |
| Neuregulin Signaling | 1.66E-06 |
| Role of Tissue Factor in Cancer | 1.66E-06 |
| PAK Signaling | 1.86E-06 |
| FAT10 Cancer Signaling Pathway | 1.91E-06 |
| Fc Epsilon RI Signaling | 1.95E-06 |
| Ferroptosis Signaling Pathway | 3.39E-06 |
| Atherosclerosis Signaling | 3.63E-06 |
| EGF Signaling | 3.63E-06 |
| Regulation of eIF4 and p70S6K Signaling | 3.89E-06 |
| IL-12 Signaling and Production in Macrophages | 4.07E-06 |
| Ceramide Signaling | 4.17E-06 |
| Non-Small Cell Lung Cancer Signaling | 4.90E-06 |
| Endometrial Cancer Signaling | 5.25E-06 |
| SPINK1 General Cancer Pathway | 5.75E-06 |
| Role Of Chondrocytes in Rheumatoid Arthritis Signaling Pathway | 6.61E-06 |
