## Supplemental Table 1-10 for "miRNA/mRNA analysis of increased TGF-β pathways drive epithelial-mesenchymal transition and regulatory T cell differentiation": Supp Table 10.docx

| **Supp Table 10. Ingenuity canonical pathways enriched by significantly upregulated miRNA-downregulated mRNA pairs in Endo+ compared to Endo- women with FDR<0.05** | |
| --- | --- |
| **Pathways** | **FDR** |
| Sirtuin Signaling Pathway | 7.59E-08 |
| Ribonucleotide Reductase Signaling Pathway | 7.59E-08 |
| Cyclins and Cell Cycle Regulation | 4.37E-07 |
| DNA Methylation and Transcriptional Repression Signaling | 4.37E-07 |
| Granzyme A Signaling | 2.09E-06 |
| Role of CHK Proteins in Cell Cycle Checkpoint Control | 4.27E-06 |
| Kinetochore Metaphase Signaling Pathway | 5.13E-06 |
| Cell Cycle Regulation by BTG Family Proteins | 1.20E-05 |
| Neutrophil Extracellular Trap Signaling Pathway | 5.13E-05 |
| Estrogen-mediated S-phase Entry | 7.59E-05 |
| Cell Cycle: G1/S Checkpoint Regulation | 8.51E-05 |
| MicroRNA Biogenesis Signaling Pathway | 1.07E-04 |
| Senescence Pathway | 2.29E-04 |
| NAD Signaling Pathway | 2.40E-04 |
| Aryl Hydrocarbon Receptor Signaling | 3.24E-04 |
| DNA damage-induced 14-3-3σ Signaling | 3.24E-04 |
| Small Cell Lung Cancer Signaling | 3.98E-04 |
| ATM Signaling | 4.17E-04 |
| Cell Cycle: G2/M DNA Damage Checkpoint Regulation | 4.27E-04 |
| NER (Nucleotide Excision Repair, Enhanced Pathway) | 8.71E-04 |
| Coronavirus Pathogenesis Pathway | 9.12E-04 |
| Glioma Signaling | 9.12E-04 |
| Mitotic Roles of Polo-Like Kinase | 9.12E-04 |
| Pancreatic Adenocarcinoma Signaling | 9.12E-04 |
| Protein Kinase A Signaling | 2.00E-03 |
| Phagosome Maturation | 2.29E-03 |
| Necroptosis Signaling Pathway | 3.31E-03 |
| Glioblastoma Multiforme Signaling | 3.31E-03 |
| Non-Small Cell Lung Cancer Signaling | 3.98E-03 |
| p53 Signaling | 4.07E-03 |
| Coronavirus Replication Pathway | 4.57E-03 |
| BER (Base Excision Repair) Pathway | 5.62E-03 |
| Macrophage Alternative Activation Signaling Pathway | 5.62E-03 |
| PFKFB4 Signaling Pathway | 6.03E-03 |
| Melanoma Signaling | 6.03E-03 |
| ID1 Signaling Pathway | 7.08E-03 |
| p38 MAPK Signaling | 7.76E-03 |
| Myelination Signaling Pathway | 8.32E-03 |
| Ferroptosis Signaling Pathway | 8.71E-03 |
| Mitochondrial Dysfunction | 8.71E-03 |
| GADD45 Signaling | 1.02E-02 |
| White Adipose Tissue Browning Pathway | 1.15E-02 |
| Remodeling of Epithelial Adherens Junctions | 1.23E-02 |
| Semaphorin Neuronal Repulsive Signaling Pathway | 1.66E-02 |
| HOTAIR Regulatory Pathway | 1.91E-02 |
| Pyrimidine Deoxyribonucleotides De Novo Biosynthesis I | 1.91E-02 |
| Androgen Signaling | 1.95E-02 |
| Estrogen Receptor Signaling | 1.95E-02 |
| Huntington's Disease Signaling | 2.04E-02 |
| CLEAR Signaling Pathway | 2.24E-02 |
| Apelin Liver Signaling Pathway | 2.82E-02 |
| Salvage Pathways of Pyrimidine Ribonucleotides | 3.02E-02 |
| Sonic Hedgehog Signaling | 3.09E-02 |
| CREB Signaling in Neurons | 3.24E-02 |
| Mouse Embryonic Stem Cell Pluripotency | 3.24E-02 |
| Pulmonary Fibrosis Idiopathic Signaling Pathway | 3.31E-02 |
| Telomerase Signaling | 3.55E-02 |
| Nucleotide Excision Repair Pathway | 3.72E-02 |
| Autophagy | 3.72E-02 |
| Prostate Cancer Signaling | 3.72E-02 |
| ERK/MAPK Signaling | 3.89E-02 |
| Bladder Cancer Signaling | 4.07E-02 |
| Axonal Guidance Signaling | 4.47E-02 |
| Intrinsic Prothrombin Activation Pathway | 4.57E-02 |
| Osteoarthritis Pathway | 4.79E-02 |
