## Supplementary figures and images for "miRNA/mRNA analysis of increased TGF-β pathways drive epithelial-mesenchymal transition and regulatory T cell differentiation"

### Figure 1.jpeg

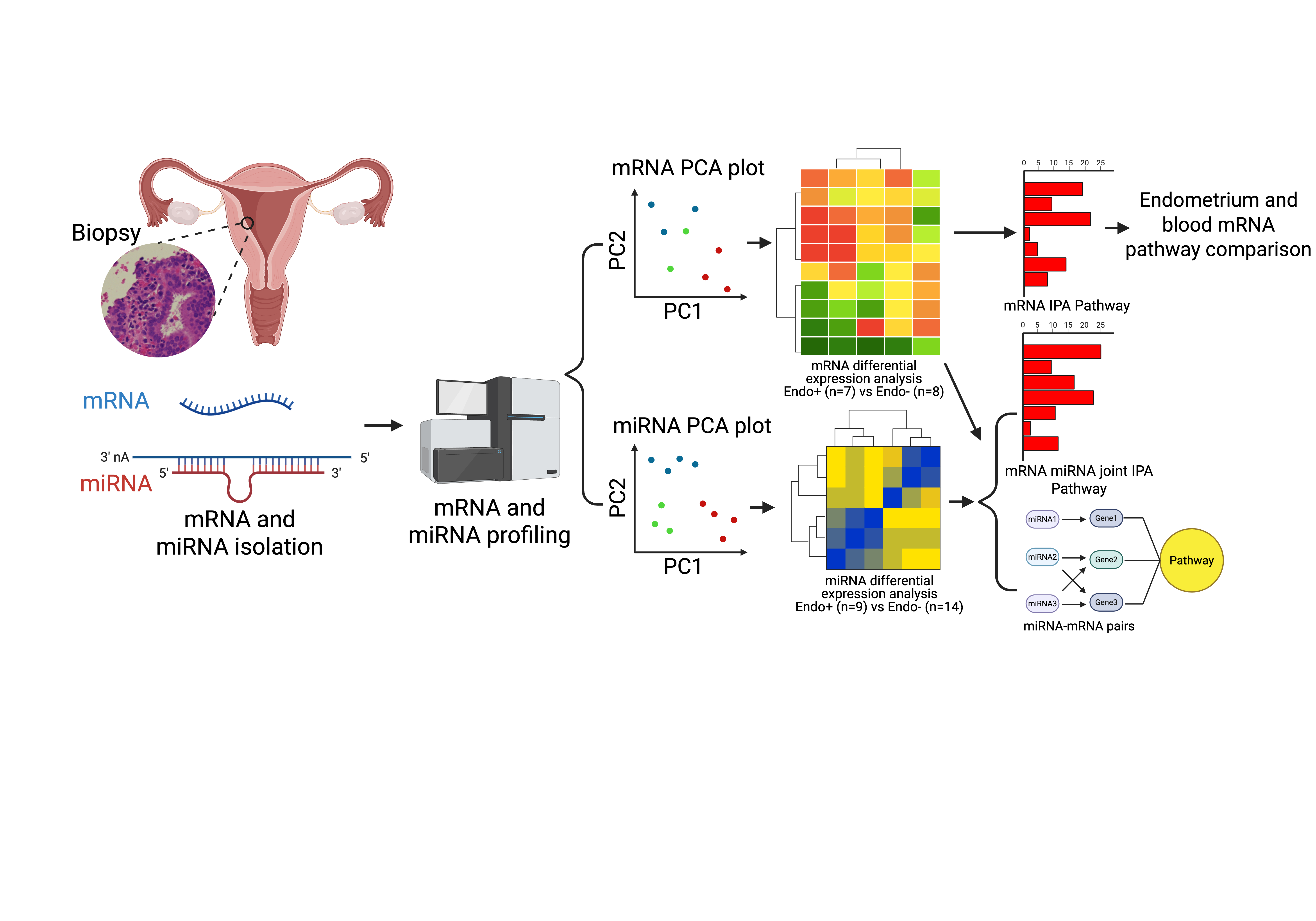

### Figure 2.tif

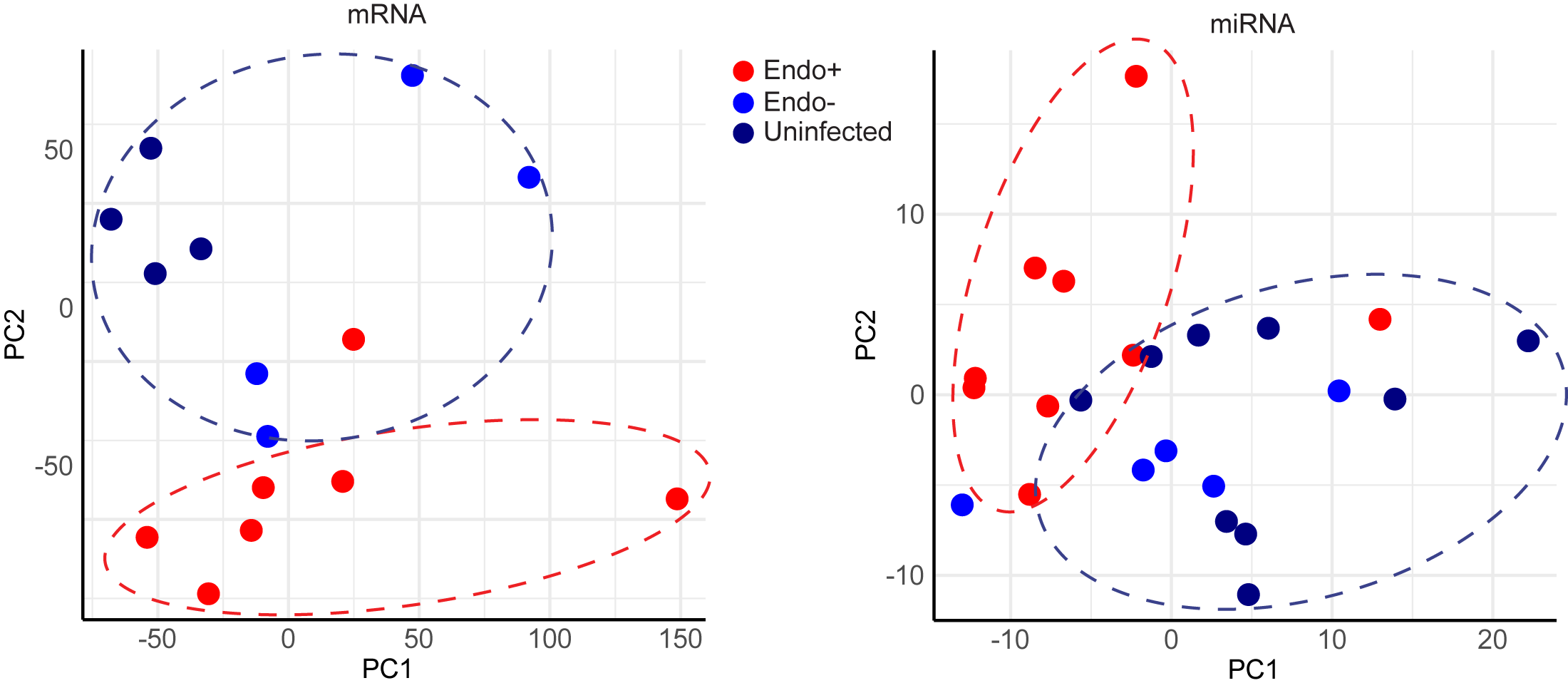

### Figure 3.tif

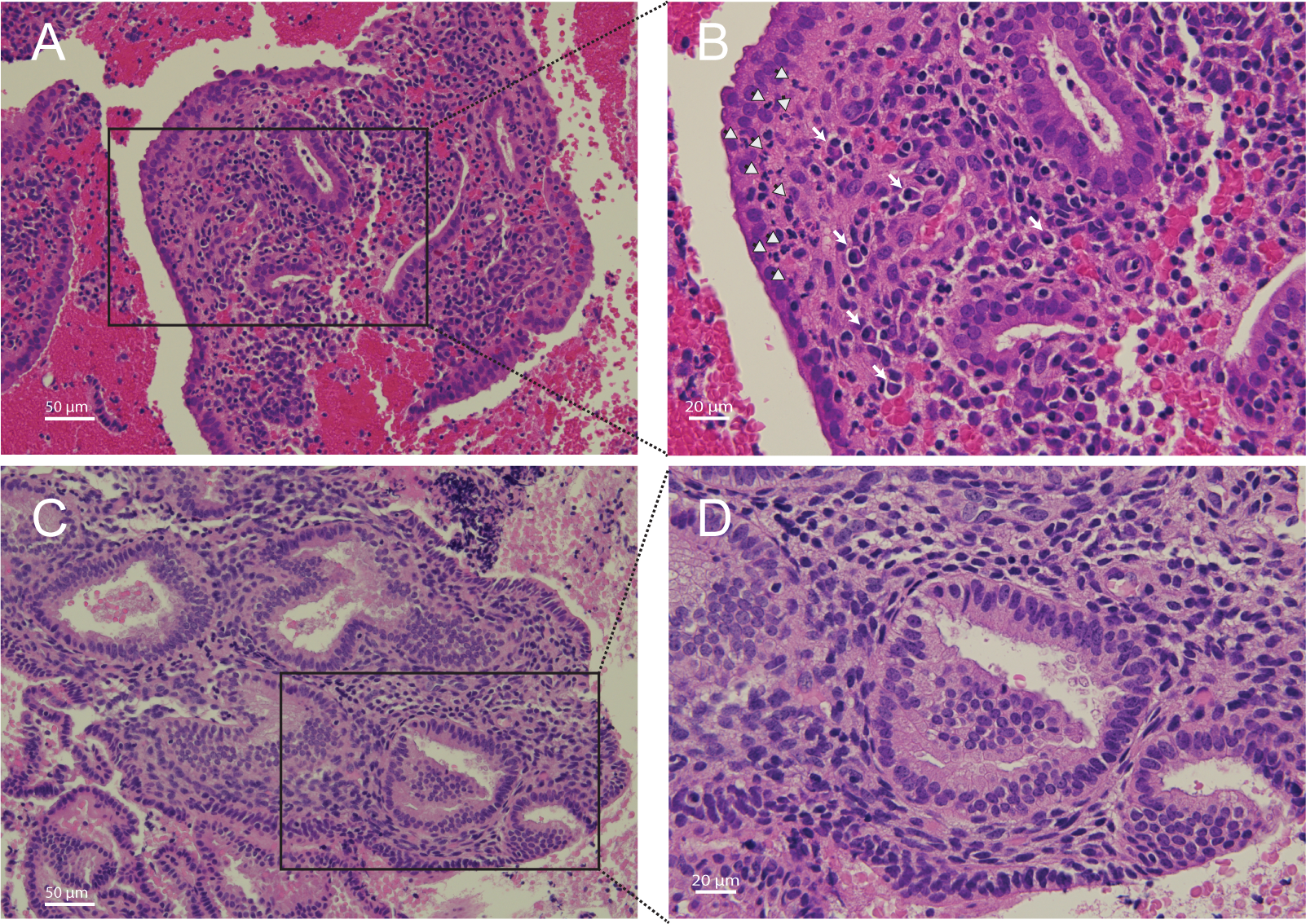

### Figure 4.tiff

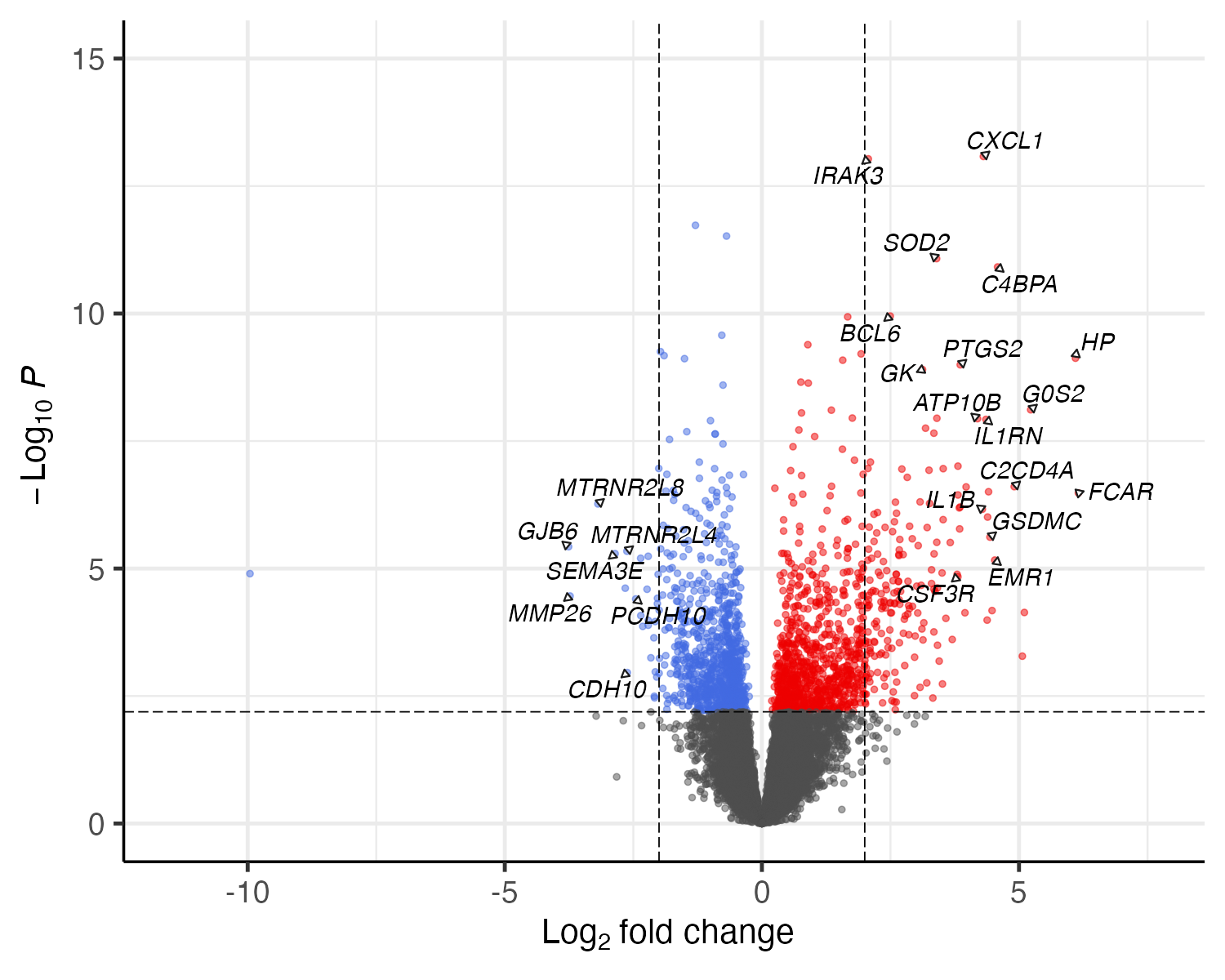

### Figure 5.tif

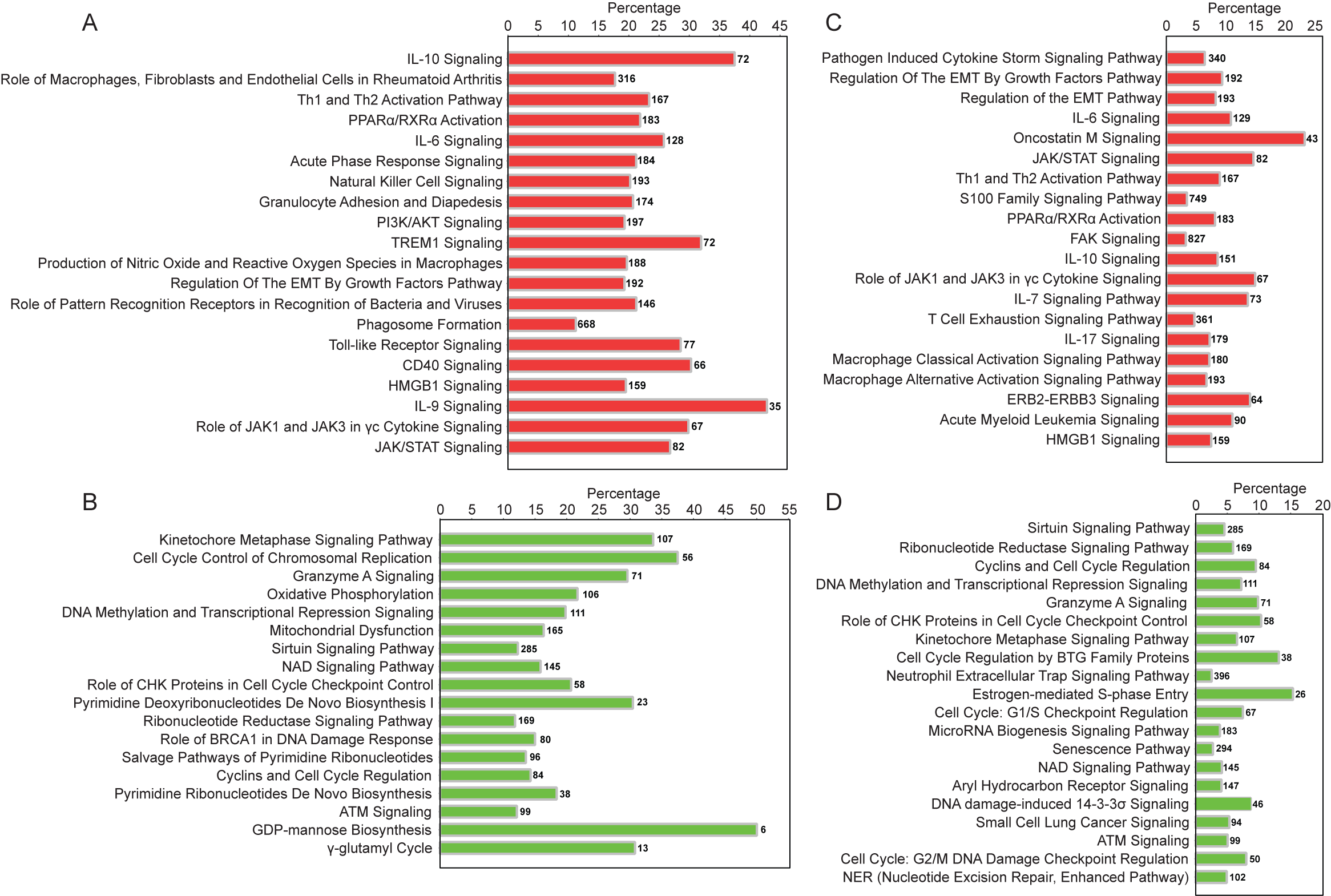

### Figure 6.tif

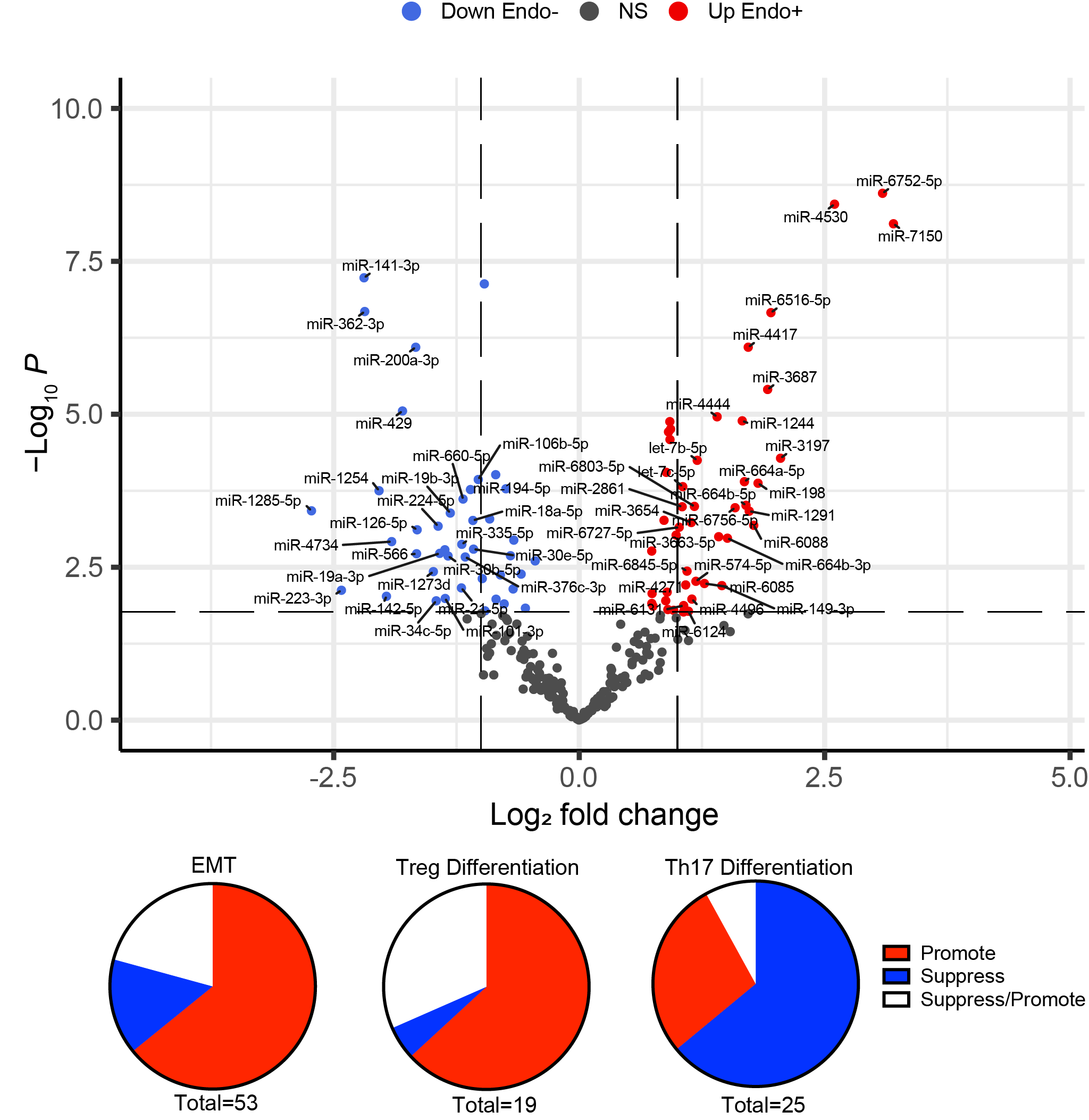

### Figure 7.tif

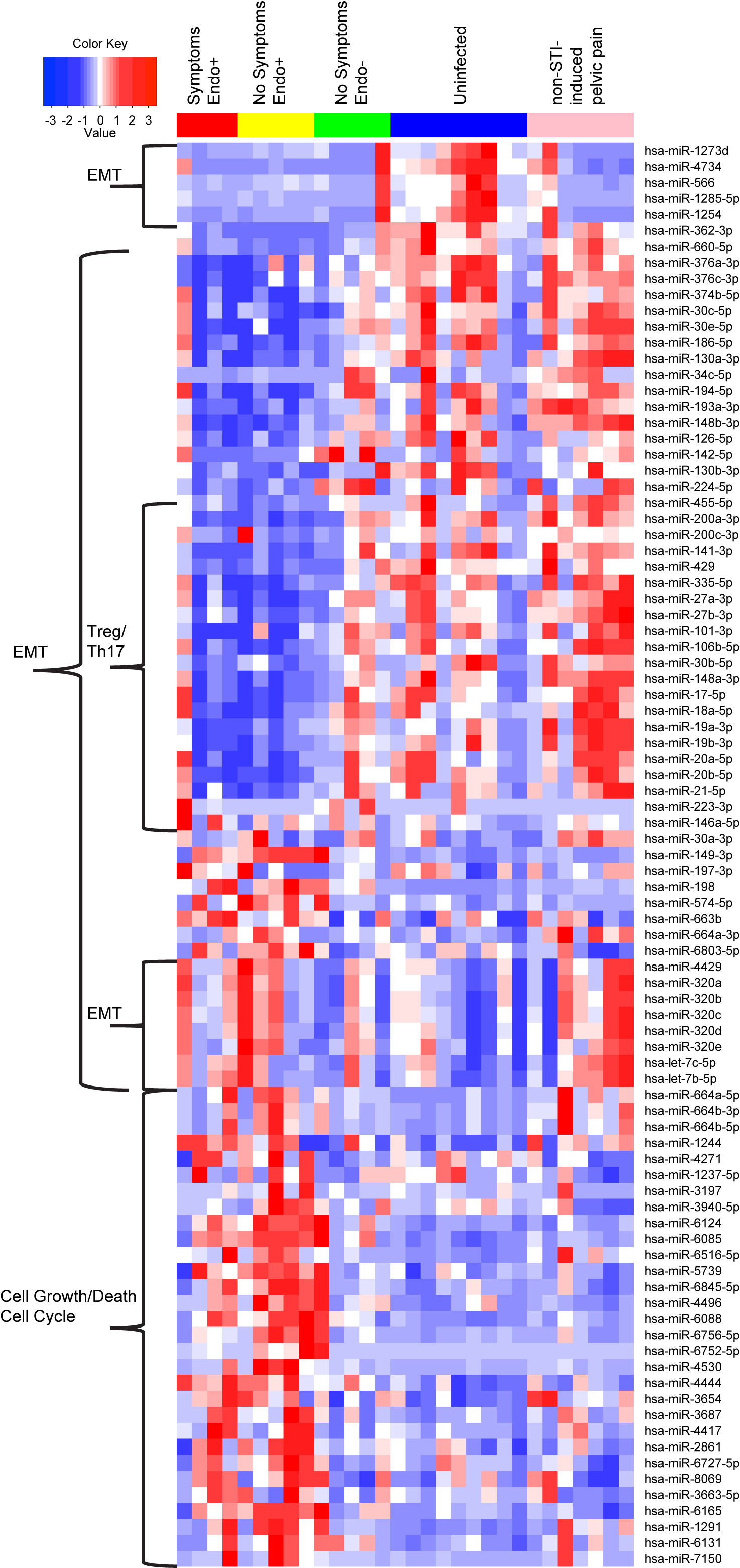
